# A quantitative model predicts how m^6^A reshapes the kinetic landscape of nucleic acid hybridization and conformational transitions

**DOI:** 10.1101/2020.12.25.424401

**Authors:** Bei Liu, Honglue Shi, Atul Rangadurai, Felix Nussbaumer, Chia-Chieh Chu, Kevin Andreas Erharter, David A. Case, Christoph Kreutz, Hashim M. Al-Hashimi

## Abstract

*N*^6^-methyladenosine (m^6^A) is a post-transcriptional modification that controls gene expression by recruiting proteins to RNA sites. The modification also slows biochemical processes through mechanisms that are not understood. Using NMR relaxation dispersion, we show that m^6^A pairs with uridine with the methylamino group in the *anti* conformation to form a Watson-Crick base pair that transiently exchanges on the millisecond timescale with a singly hydrogen-bonded low-populated (1%) mismatch-like conformation in which the methylamino group is *syn.* This ability to rapidly interchange between Watson-Crick or mismatch-like forms, combined with different *syn*:*anti* isomer preferences when paired (~1:100) versus unpaired (~10:1), explains how m^6^A robustly slows duplex annealing without affecting melting via two pathways in which isomerization occurs before or after duplex annealing. Our model quantitatively predicts how m^6^A reshapes the kinetic landscape of nucleic acid hybridization and conformational transitions, and provides an explanation for why the modification robustly slows diverse cellular processes.

## Introduction

*N*^6^-methyladenosine (m^6^A) (Fig. 1a) is an abundant RNA modification^1,2^ that helps control gene expression in a variety of physiological processes including cellular differentiation, stress response, viral infection, and cancer progression^3–5^. m^6^A is also the most prevalent form of DNA methylation in prokaryotes where it is used to distinguish benign host DNA from potentially pathogenic nonhost DNA^6^. Although under debate^7^, there is also evidence for m^6^A in mammalian DNA where it is proposed to play roles in transcription suppression and gene silencing^8,9^.

**Fig. 1.**
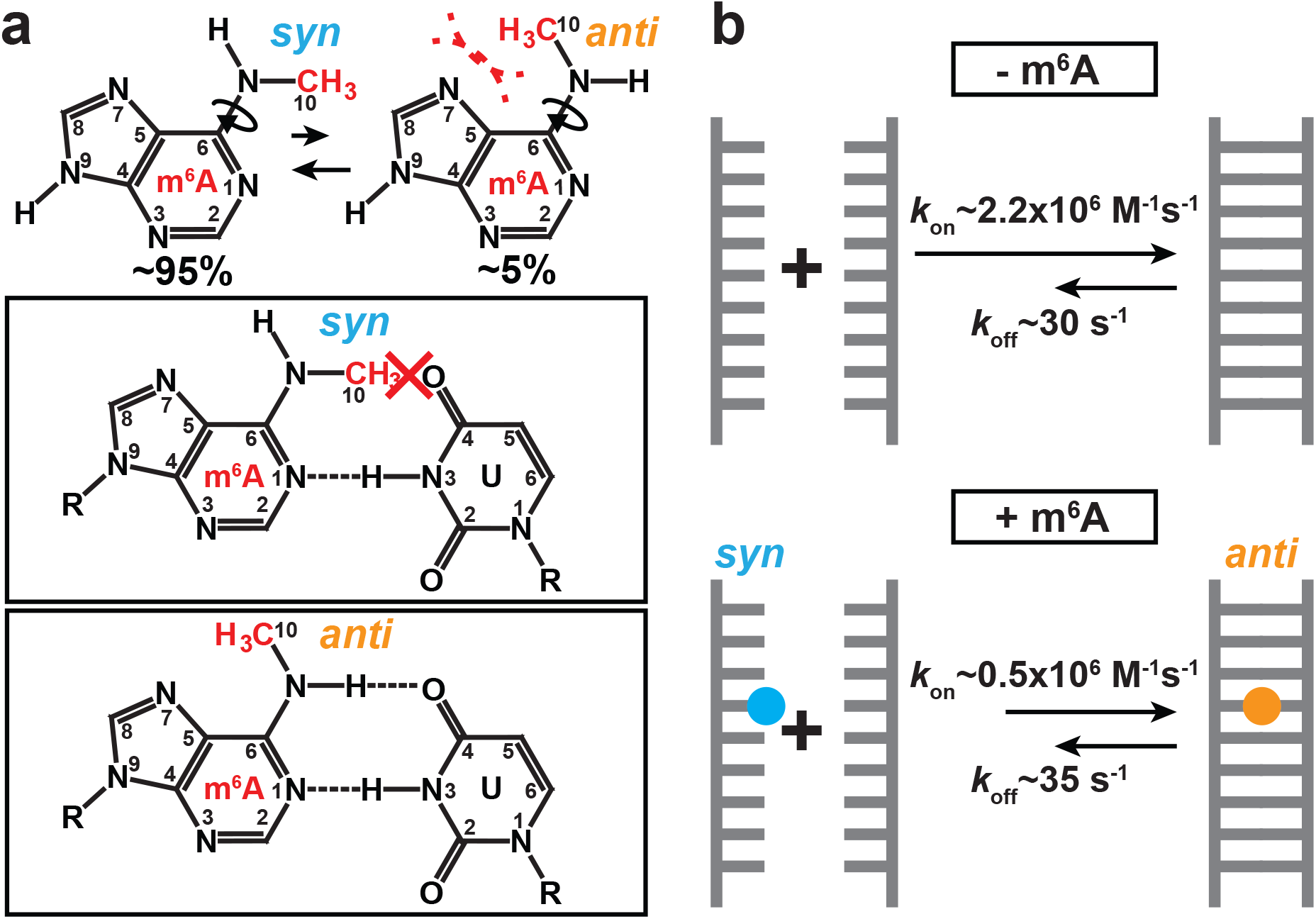
The *syn* and *anti* isomers of m^6^A. **a,** The m^6^A nucleobase shows a 20:1 preference for the *syn* isomer due to unfavorable steric interactions (shown in dashed red lines) in the *anti* isomer^12,25^. In a duplex, the *syn* isomer impedes Watson-Crick pairing, and the *anti* isomer becomes the dominant form. **b,** Apparent annealing (*k*_on_) and melting (*k*_off_) rate constants for unmethylated (-m^6^A) and methylated (+m^6^A) dsRNA. Rate constants shown were obtained from CEST measurements on dsGGACU with and without m^6^A at T = 65°C^21^.

In RNAs, m^6^A is thought to primarily function by recruiting proteins to specific modified sites (reviewed in^3–5^). However, there is also growing evidence that the modification can impact a range of biochemical processes by changing the behavior of the methylated RNAs^10,11^. For example, by destabilizing canonical double-stranded RNA (dsRNA)^12^, m^6^A has been shown to promote binding of proteins to single-stranded regions of RNAs (ssRNA)^10^. The modification has also been shown to slow biochemical processes that involve base pairing. For example, in mRNAs, m^6^A delays tRNA selection and reduces translation efficiency *in vitro*^13^ and *in vivo*^14^ by 3-15-fold. In mRNA introns, m^6^A slows splicing and promotes alternative splicing *in vivo*^15^. Additionally, m^6^A reduces the rate of NTP incorporation during DNA replication^16^ and reverse transcription^17^ *in vitro* by 2-13-fold.

Recently, using NMR relaxation-dispersion (RD)^18–20^, we showed that m^6^A preferentially slows the apparent rate of RNA duplex annealing by ~5-10-fold while having little effect on the apparent rate of duplex melting^21^ (Fig. 1b). This impact of m^6^A on hybridization kinetics stands in contrast to mismatches, which slow the rate of duplex annealing but also substantially increase the rate of duplex melting by up to ~100-fold^22–24^. How m^6^A selectively slows duplex annealing remains unknown. The comparable m^6^A induced slowdown observed for duplex annealing and a variety of biochemical processes indicates that a common mechanism might be at play^13,16,17^.

It has been known for many decades that the methylamino group of the m^6^A nucleobase can form two rotational isomers which interconvert on the millisecond timescale^25,26^ (Fig. 1a). The preferred *syn* isomer^12,25,26^ cannot form a canonical Watson-Crick base pair (bp) with uridine as the methyl group impedes one of the hydrogen-bonds (H-bonds) (Fig. 1a). Rather, when paired with uridine, the methylamino group rotates into the energetically disfavored *anti* isomer and forms a canonical m^6^A-U Watson-Crick bp (Fig. 1a). As isomerization is energetically disfavored, it has been proposed to explain how m^6^A destabilizes dsRNA via the so-called “spring-loading”^12^ mechanism despite forming a canonical Watson-Crick m^6^A-U bp.

Here, using NMR relaxation dispersion (RD), we show that m^6^A with the methylamino group in the *anti* conformation forms a Watson-Crick base pair with uridine that transiently exchanges on the millisecond timescale with an unusual singly hydrogen-bonded, low-populated (1%), and mismatch-like conformation through isomerization of the methylamino group to the *syn* conformation. This ability to rapidly interchange between Watson-Crick or mismatch forms, combined with different *syn*:*anti* isomers preferences when paired versus unpaired, explains how m^6^A robustly and selectively slows duplex annealing without affecting melting via two pathways in which isomerization occurs before or after duplex annealing. We develop a model that quantitatively predicts how m^6^A reshapes the kinetic landscape of nucleic acid hybridization, that could explain why the modification robustly slows a variety of cellular processes. The model also predicts that m^6^A more substantially slows fast intra-molecular RNA conformational transitions, and this prediction was verified experimentally by using NMR.

## Results

### Kinetics of m^6^A methylamino isomerization in ssRNA

We developed and tested a simple model that can explain how m^6^A slows duplex annealing while not affecting the melting rate. The model assumes that the minor *anti* isomer of m^6^A hybridizes with apparent annealing (*k*_on_) and melting (*k*_off_) rate constants similar to those of the unmethylated RNA. This assumption is reasonable given that like unmethylated adenine, the *anti* isomer forms a canonical m^6^A-U Watson-Crick bp when paired with uridine^11,12,25,26^. Since the *syn* isomer is incapable of Watson-Crick pairing with uridine, the model assumes that hybridization only proceeds via annealing of the single-strand containing the minor *anti* isomer (ssRNA^*anti*^) through a conformational selection (CS) type pathway^27,28^ (Fig. 2a). The apparent *k*_on_ would then be reduced relative to the unmethylated RNA because the methylamino group has to rotate from the major *syn* to the minor *anti* isomer prior to hybridization (Fig. 2a). However, because *anti* is the preferred isomer in the canonical duplex, and because hybridization is rate limiting under our experimental conditions (see below), the apparent *k*_off_ would remain equivalent to that of the unmethylated duplex.

**Fig. 2.**
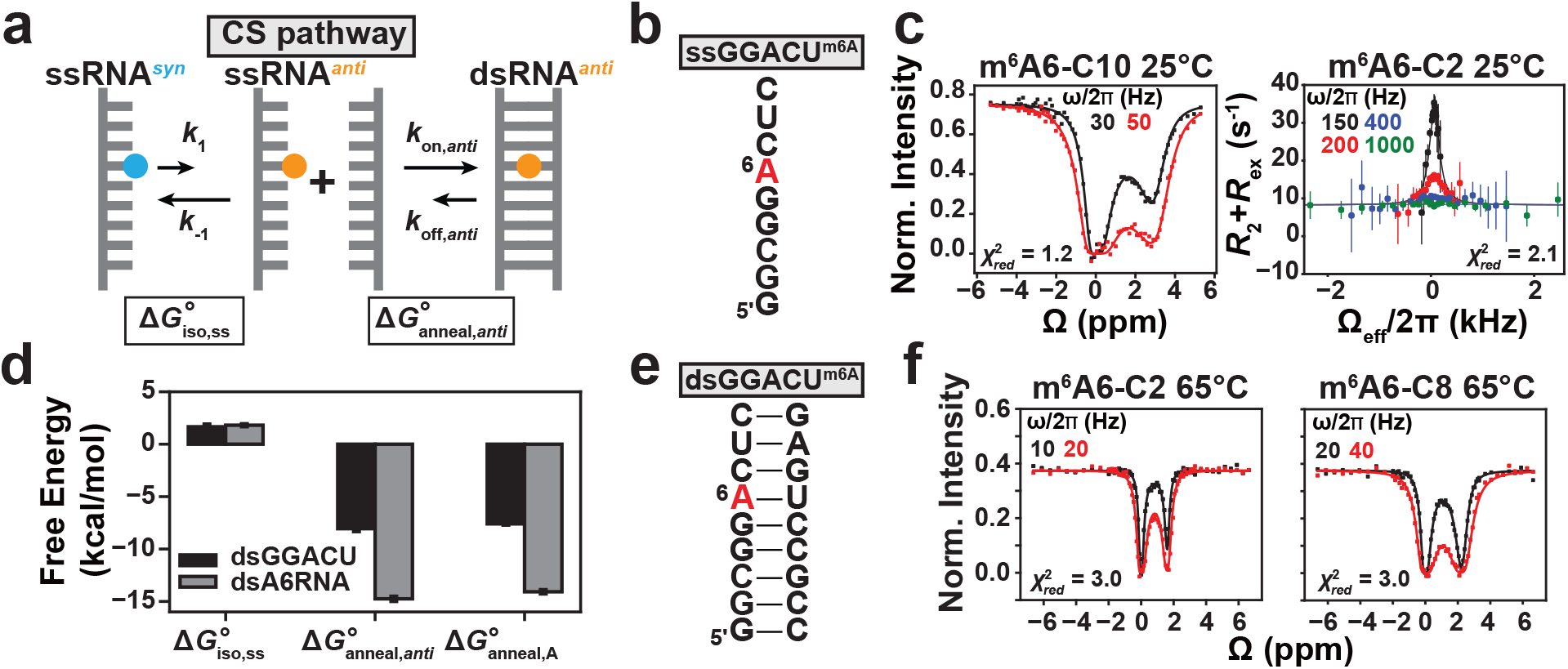
Testing a conformational selection kinetic model for m^6^A hybridization. **a**, The CS pathway. 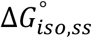 is the free energy of methylamino isomerization in ssRNA. 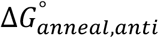 is the free energy of annealing the methylated ssRNA when m^6^A is *anti*. **b**, ssGGACU sequence with the m^6^A site highlighted in red. **c**, ^13^C CEST profile for m^6^A6-C10 and off-resonance ^13^C *R*_1ρ_ RD profile for m^6^A6-C2 in ssGGACU^m6A^. **d**, Free energy decomposition (Methods) of the CS pathway for dsGGACU^m6A^ at T = 65°C and dsA6RNA^m6A^ (Extended Data Fig. 1) at T = 20°C. 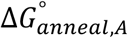 is the free energy of annealing unmethylated ssRNA and the value for dsGGACU was obtained from a prior study using RD measurements^21^, and for dsA6RNA was measured using UV melting experiments (Supplementary Table 4). The uncertainty in free energies were obtained from Monte-Carlo simulations as described in Methods for RD measurements, or from standard deviations for UV melting measurements. **e**, The dsGGACU^m6A^ duplex with the m^6^A site highlighted in red. **f**, ^13^C CEST profiles for m^6^A6-C2 and C8 in dsGGACU^m6A^ at T = 65°C (data obtained from a prior study^21^). Solid lines in panels **c** and **f** denote a 2-state and constrained 3-state fit to the CS pathway, using Bloch-McConnell equations as described in Methods. Buffer conditions for NMR experiments are described in Methods. RF field powers used for CEST and spin-lock powers used for *R*_1ρ_ are color-coded. Error bars for CEST (smaller than data points) and *R*_1ρ_ profiles were obtained from standard deviations and Monte-Carlo simulations, respectively as described in Methods.

To test this CS model, we first used NMR RD to measure the isomerization kinetics in a ssRNA containing the most abundant m^6^A consensus sequence^1,2^ in eukaryotic mRNAs (ssGGACU^m6A^, Fig. 2b). This was important given that prior kinetic measurements of isomerization were performed on the m^6^A nucleobase dissolved in organic solvents and the kinetics may differ in ssRNA under aqueous conditions^25^.

To enable the RD measurements, we used organic synthesis (Methods) to incorporate m^6^A ^13^C-labeled at the base C2 and C8, or methyl C10 carbons (Extended Data Fig. 1) into ssGGACU. We then performed NMR Chemical Exchange Saturation Transfer (CEST)^29–31^ and off-resonance spin relaxation in the rotating frame (*R*_1ρ_) experiments^18–20^ to measure the isomerization kinetics. Together, *R*_1ρ_ and CEST, which are optimized for different nuclei and exchange kinetics, allowed robust characterization of chemical exchange between the major ground-state (GS) *syn* methylamino and the low-populated and short-lived “excited-state” (ES)^32^ *anti* methylamino isomer in unpaired m^6^A.

In the m^6^A-C10 CEST profile (Fig. 2c), we observed a minor dip indicating that the methyl group in ssGGACU^m6A^ undergoes conformational exchange with a sparsely populated ES. The dip was observed at a chemical shift Δω_C10_ = ω_ES_ − ω_GS_ = 3 ppm, which was in good agreement with the value predicted for the *anti* isomer (Δω_C10_ = 3-5 ppm) using density functional theory (DFT) calculations^33^ (Methods). In addition, an RD peak was observed for m^6^A-C2 at Δω_C2_ = −0.6 ppm (Fig. 2c). The same C2 RD was observed in methylated but not unmethylated AMP, as expected if the RD is reporting on isomerization (Extended Data Fig. 2a).

Based on a 2-state fit of the m^6^A-C10 and m^6^A-C2 RD data (Fig. 2c), the population of the ssRNA^*anti*^ isomer in ssGGACU^m6A^ was ~9% and the exchange rate for isomerization (*k*_ex_ = *k*_1_ + *k*_−1_, where *k*_1_ and *k*_−1_ are the forward and backward rate constants, respectively) was ~600 s^−1^ at T = 25°C (Supplementary Table 1). The population was ~2-fold higher than the value measured in the nucleobase in organic solvent (Fig. 1a)^25^ while the exchange rate was ~20-fold faster, and in better agreement with values reported recently for ssDNA^34^ (at T = 45°C, Supplementary Table 1). Similar *syn*-*anti* isomerization kinetics were obtained for another different sequence (Extended Data Fig. 2b).

### m^6^A(*anti*)-U a nd A-U have similar thermodynamic stabilities in dsRNA

Before testing whether the CS model can predict the hybridization kinetics of methylated duplexes, we tested a thermodynamic prediction made by our model, namely that the energetics of annealing a single-strand containing the *anti* isomer of m^6^A should be similar to the energetics of annealing the unmethylated control. In this scenario, m^6^A destabilizes a duplex^12^ solely due to the conformational penalty 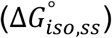 accompanying *syn* to *anti* isomerization in the ssRNA, which we have measured here for ssGGACU^m6A^ using NMR RD.

To test this prediction, we decomposed (Fig. 2a) the overall annealing energetics 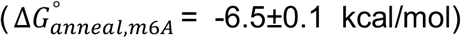 of methylated dsGGACU^m6A^ (Fig. 2e) measured previously using melting experiments^21^ into the sum of 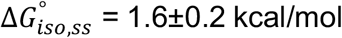 plus the desired annealing energetics 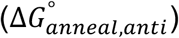 of m^6^A when it adopts the *anti* isomer,

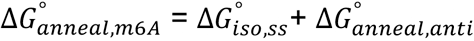

Indeed, we find that 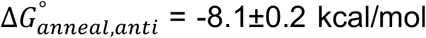 is similar to that measured for the unmethylated RNA 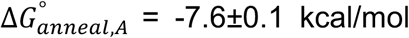, with the methyl group being only slightly stabilizing within error by 0.5±0.2 kcal/mol. A similar result was obtained for a different duplex (Fig. 2d) and a similar conclusion was also reached previously using the isomerization energetics measured in the nucleobase^25,26^. Therefore, with respect to the thermodynamics of annealing canonical duplexes, m^6^A in the *anti* isomer behaves similarly (within <0.5 kcal/mol) to unmethylated adenine and m^6^A primarily destabilizes dsRNA due to the conformational penalty accompanying isomerization, consistent with the previously proposed “spring-loading” mechanism^12^.

### Testing the conformational selection kinetic model

Next, we tested whether the CS kinetic model could explain the impact of m^6^A on the hybridization kinetics of the dsGGACU^m6A^ RNA measured recently using NMR RD^21^. These experiments were performed at T = 65°C under conditions in which the duplex was the GS, and the ssRNA comprising two species in rapid equilibrium (ssRNA^*syn*^⇌ssRNA^*anti*^) was the ES with population of ~25%. Based on a 2-state fit (dsRNA⇌ssRNA) of the m^6^A6-C2 and m^6^A6-C8 RD data (Extended Data Fig. 3a), m^6^A reduced the apparent rate of dsGGACU^m6A^ annealing 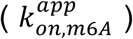 relative to the unmethylated control (*k*_*on*_) by 5-fold while having little impact on the melting rate 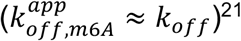.

We used the 3-state CS model to simulate the m^6^A6-C8 and m^6^A6-C2 RD profiles measured for the methylated dsGGACU^m6A^ duplex. The exchange parameters for the first isomerization step (ssRNA^*syn*^⇌ssRNA^*anti*^) were fixed to the values determined independently from RD measurements on ssGGACU^m6A^ (Extended Data Fig. 2c). *k*_*off,anti*_ was assumed to be equal to *k*_*off*_ measured for the unmethylated dsGGACU. This assumption is reasonable considering that hybridization is rate limiting under our experimental conditions, and given the similarity between the experimentally measured *k*_*off*_ for methylated and unmethylated duplexes^21^. The value of *k*_*on,anti*_ was slightly adjusted relative to *k*_*on*_ of the unmethylated control (*k*_*on,anti*_ ≈ 2 × *k*_*on*_) to take into account small differences in their annealing energetics (Fig. 2a). The remaining NMR exchange parameters (Δω, *R*_1_, *R*_2_ of GS and two ESs) for the hybridization and isomerization steps were fixed to the values obtained from the 2-state fit of the RD data measured for dsGGACU^m6A^ and ssGGACU^m6A^ (Methods).

Interestingly, this simulation with no adjustable parameters satisfactorily reproduced the RD data with 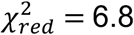. This can be compared with 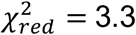 (Extended Data Fig. 3a) obtained from a 2-state fit of the RD data with six adjustable parameters. As a negative control, the agreement deteriorated considerably 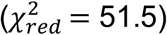 (Extended Data Fig. 3b) when decreasing the exchange rate by 20-fold to mimic values observed for the nucleobase in organic solvents^25^. A constrained 3-state fit to the RD data using the CS model in which the exchange parameters were allowed to vary within experimental error by one standard deviation, and in which the ratio (but not absolute magnitude) of *k*_*on,anti*_ and *k*_*off,anti*_ was constrained to preserve the free energy of the hybridization step improved the agreement to 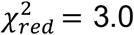 (Methods, Fig. 2f) and yielded *k*_*on,anti*_ ≈ 2 × *k*_*on*_ and *k*_*off,anti*_ ≈ *k*_*off*_ (Supplementary Table 2). Therefore, even when it to comes to hybridization kinetics, m^6^A in the *anti* isomer behaves similarly to unmethylated adenine.

These results provide a plausible explanation for the unique impact of m^6^A on RNA hybridization kinetics at T = 65°C. m^6^A does not impact the apparent melting rate because the dominant isomer in the duplex is *anti* and it melts at a rate comparable to that of the unmethylated RNA. On the other hand, m^6^A slows the apparent annealing rate by ~5-fold due to the ~10-fold lower equilibrium population of the ssRNA^*anti*^ intermediate relative to the unmethylated ssRNA control and because the ssRNA^*anti*^ intermediate anneals at a 2-fold faster rate relative to its unmethylated counterpart.

### A new hybridization intermediate at T = 55°C

To test the robustness of the CS model, we repeated the RD measurements of hybridization kinetics of dsGGACU^m6A^ at a lower temperature of T = 55°C. Based on a 2-state fit of the adenine C8 RD data, which only reports on hybridization and not this new ES (Extended Data Fig. 4a), m^6^A reduced the apparent annealing rate by 20-fold while minimally (~1.6 fold) impacting the apparent melting rate under these conditions (Extended Data Fig. 4b).

Interestingly, we observed evidence for a new ES, which manifested as a second minor dip in the m^6^A-C2 CEST profile (Fig. 3a). This new ES dip at Δω_C2_ ~2 ppm was also observed at lower temperatures in another dsRNA (dsA6RNA^m6A^) sequence context (Extended Data Fig. 5 and Supplementary Table 1). The fact that this new ES was not observed in ssGGACU^m6A^ indicated that it very likely was a dsRNA conformation. The new ES was likely not observed at higher temperature T = 65°C (Fig. 2f)^21^ because it was masked by the higher RD contribution from the more populated ssRNA ES.

**Fig. 3.**
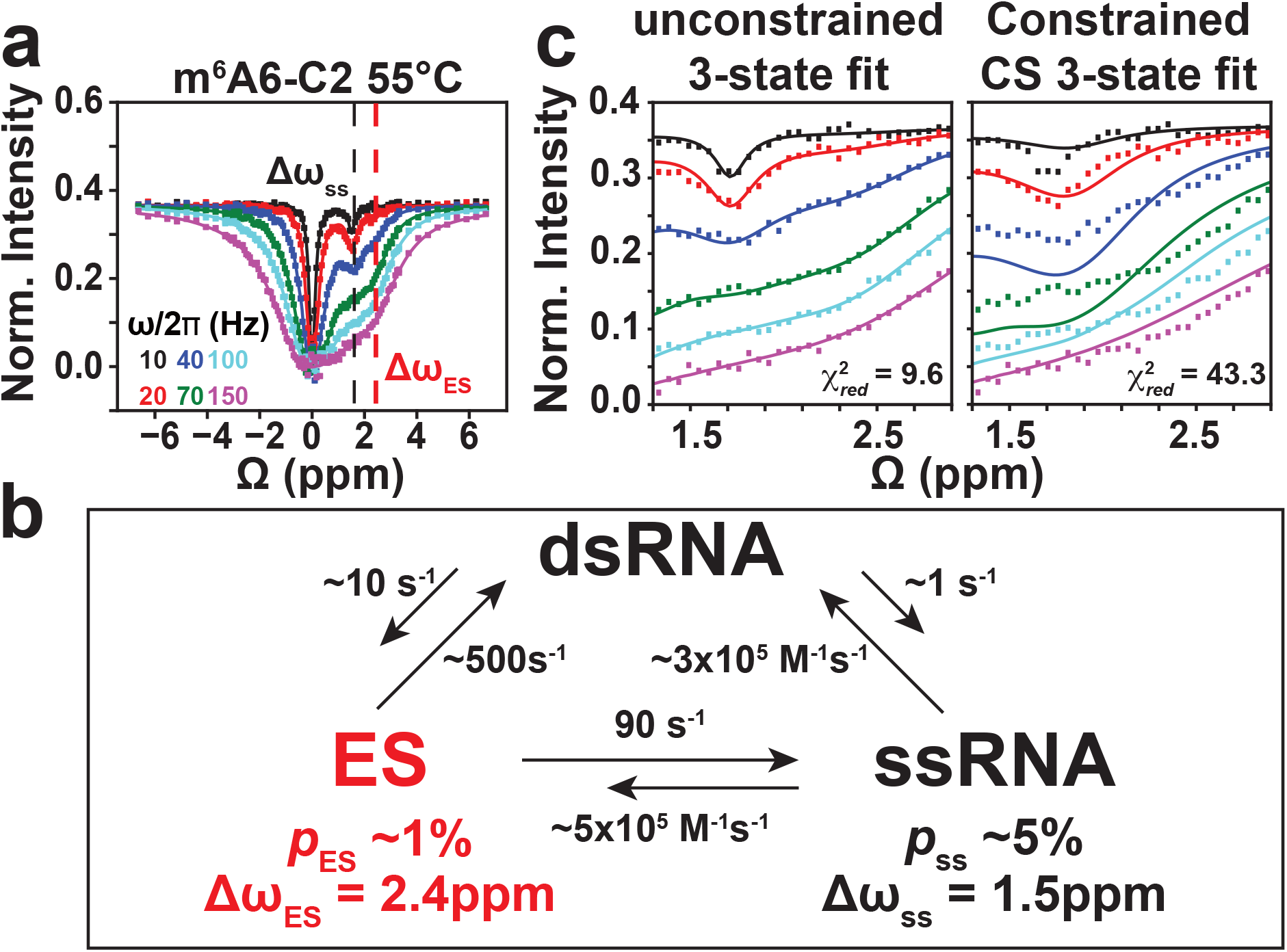
A new hybridization intermediate for dsGGACU^m6A6^ at T = 55°C. **a**, ^13^C CEST profile for m^6^A6 C2 in dsGGACU^m6A6^ at T = 55°C shows a second dip at Δω_ES_ that is distinct from the ssRNA ES at Δω_ss_. **b**, Exchange parameters (Supplementary Table 3) from 3-state fit to the RD data using a triangular model. **c**, Zoom in to the m^6^A6 C2 CEST profiles comparing results from an unconstrained 3-state fit to the Bloch-McConnell equations assuming the triangular model and a constrained 3-state fit assuming a linear CS model. Error bars for CEST profiles (smaller than data points) were obtained using standard deviation of 3 measurements of peak intensity with zero relaxation delay as described in Methods. RF field powers used for CEST are color-coded.

The m^6^A-C2 RD data (Fig. 3a) could be satisfactorily fit to a 3-state model which includes dsRNA, ssRNA, and the new ES. Among several 3-state topologies tested^35^ (see Extended Data Fig. 4d), the best agreement was obtained with models that place the new ES on-pathway between the dsRNA and ssRNA (Fig. 3b). Therefore, these results provide direct evidence for a new dsRNA on-pathway hybridization intermediate and the CS pathway alone cannot fully explain the hybridization kinetics at T = 55°C. Indeed, simulations using the CS model did not reproduce the m^6^A-C2 RD data at 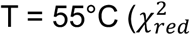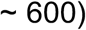 (Extended Data Fig. 4c) and neither did a constrained 3-state fit to the CS model 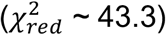 (Fig. 3c) because the model fails to account for the RD contribution from the new ES.

### The new dsRNA hybridization intermediate features a m^6^(*syn*)A···U stabilized by a single H-bond

Understanding how m^6^A selectively slows annealing of dsGGACU at T = 55°C by 20-fold without affecting the melting rate requires that we characterize the newly identified intermediate, which can be part of a new hybridization pathway distinct from the CS pathway.

Although never observed previously, one possibility is that the new intermediate is a dsRNA conformation in which the methylamino group rotates into the energetically favored *syn* isomer. Although such a conformation is predicted to be highly energetically disfavored, given the loss of at least one Watson-Crick H-bond, this loss in energetic stability would be partly compensated for by a gain in stability of ~−1.5 kcal/mol from restoring the energetically favored *syn* isomer. Such an intermediate would allow for an induced-fit (IF) type hybridization pathway, in which isomerization of the methylamino group occurs following and not before initial duplex formation (see Fig. 5a).

To test this proposed conformation for the ES, we performed an array of NMR RD experiments using a stable hairpin variant of dsGGACU^m6A^ (hpGGACU^m6A^, Fig. 4a) with a much higher melting temperature (Tm is predicted to be ~80°C), designed to eliminate any background RD contribution from the ssRNA across a range of temperatures. Interestingly, we observed 2-state RD for both m^6^A-C10 (Fig. 4b) and m^6^A-C2 (Extended Data Fig. 6a) at T = 55°C. A global fit of the data yielded an ES population (~1%), *k*_ex_ (~500 s^−1^), and Δω_C2_ = 2.5 ppm that were in very good agreement with the values (Supplementary Table 3) measured for the on-pathway ES hybridization intermediate in dsGGACU^m6A^. The Δω_C10_ and Δω_C2_ values were also in very good agreement with values predicted for m^6^(*syn*)A···U based on DFT calculations (Fig. 4g). Additional support that in the ES the methylamino group is *syn* comes from the kinetic rate constants of inter-conversion (Supplementary Note 1).

**Fig. 4.**
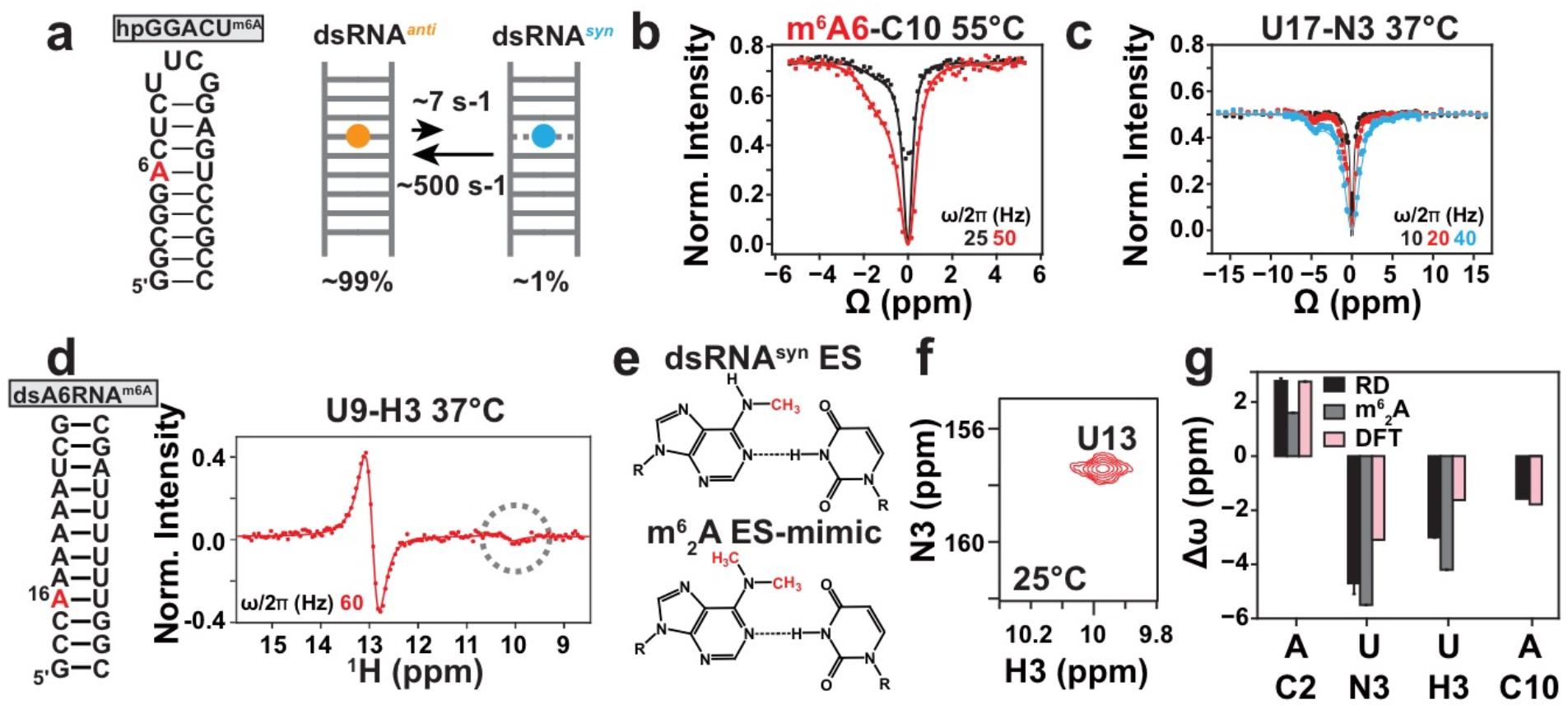
Characterizing the conformation of the new ES intermediate. **a**, The hpGGACU^m6A^ hairpin construct with the m^6^A site highlighted in red (left) and exchange parameters between dsRNA^*anti*^ and dsRNA^*syn*^ measured at T = 55°C (right). **b**, ^13^C CEST profile measured for m^6^A6-C10 in hpGGACU^m6A^ at T = 55°C. **c**, ^15^N CEST profile measured for U17-N3 in hpGGACU^m6A^ at T = 37°C. **d**, The dsA6RNA^m6A^ duplex (left) and ^1^H CEST profile for U9-H3 at T = 37°C (right). The minor peak is highlighted in the gray circle. **e**, Chemical structures of proposed dsRNA^*syn*^ ES and m^6^_2_A ES-mimic. **f**, 2D [^15^N, ^1^H] HSQC spectra of U13-N3 ^15^N site-labeled dsGGACU^m62A^ at T = 25°C. **g**, Comparison of the chemical shift differences (Δω_ES-GS_ = ω_ES_ − ω_GS_) measured using RD in hpGGACU^m6A^ (A C2/C10, U N3) and dsA6^m6A^ (U H3) at T = 37°C (RD), when taking the difference between the chemical shifts measured for dsGGACU^m62A^ and dsGGACU^m6A^ (m^6^_2_A) and calculated using DFT as the difference between an m^6^(syn)A···U conformational ensemble and a Watson-Crick m^6^A(*anti*)-U bp (DFT) (Methods). Values for m^6^_2_A C10 are not shown because it is the site of modification. Solid lines in panel **b**, **c**, **d** denote a fit to the Bloch-McConnell equations to a 2-state exchange model (Methods). RF field powers for CEST profiles are color coded. Error bars for CEST profiles (smaller than data points) were obtained from standard deviations as described in Methods. Error bars in Δω (panel **g**) was obtained using Monte-Carlo simulations as described in Methods.

To gauge the nature of the Watson-Crick (m^6^A)N1···H3-N3(U) H-bond in the ES, we performed additional RD experiments targeting the N3 and H3 atoms of the partner Uridine. We observed ^15^N (Fig. 4c) and ^1^H (Fig. 4d) RD only for the uridine partner of m^6^A (Extended Data Fig. 6a), and 2-state fit of the data yielded exchange parameters similar to those obtained from the carbon C2/C10 data (Extended Data Fig. 6a), indicating that they are reporting on the same ES. The Δω_N3_ = −4.8 ppm and Δω_H3_ = −3 ppm values indicated substantial weakening of the remaining H-bond in the ES^36^ (Fig. 4e). Indeed, a structural model for the m^6^(*syn*)A···U ES conformation that predicts the ES chemical shifts well based on DFT (Fig. 4g), features a slightly (by 0.4 Å) elongated (m^6^A)N1···H3-N3(U) H-bond (Extended Data Fig. 6b). Note that while a minor peak was not observed in the ^1^H CEST profile for U17-H3 in hpGGACU^m6A^, simulations indicate that this could be due to the 2-fold lower ES population (Extended Data Fig. 6c and Supplementary Table 1).

These results establish that the m^6^A methylamino group can also isomerize even in the context of a duplex m^6^(*anti*)A-U Watson-Crick bp and show that the preferences for the *syn:anti* isomers is inverted from ~10:1 in the unpaired single-strand to ~1:100 in the paired dsRNA.

### Chemical shift fingerprinting the m^6^(*syn*)A ···U ES using m^6^_2_A

To further verify the unusual m^6^(*syn*)A···U conformation proposed for the ES, we stabilized this species and rendered it the dominant conformation by replacing the m^6^A amino proton with a second methyl group so as to eliminate the GS Watson-Crick H-bond (Fig. 4e). This *N*^6^,*N*^6^-dimethyl adenine (m^6^_2_A) modification (Fig. 4e) is also a naturally occurring RNA modification^37^.

Comparison of NMR spectra of dsGGACU with and without m^6^_2_A showed that the modification primarily affected the methylated bp while minimally impacting other neighboring bps (Extended Data Fig. 7a). Both the m^6^_2_A-C2 and U-N3 chemical shifts of the m^6^_2_A modified dsGGACU (dsGGACU^m62A^) were in very good agreement with those measured for the ES in dsGGACU^m6A^ using RD (Fig. 4g). In addition, we observed an upfield shifted imino proton resonance (at ~10 ppm) which could unambiguously be assigned via site labelling to the m^6^A partner U13-H3 (Fig. 4f and Extended Data Fig. 7a). This along with NOE-based distance connectivity (Extended Data Fig. 7a) indicate that the m^6^(*syn*)A···U ES likely retains a weaker (m^6^A6)N1···H-N3(U13) Watson-Crick H-bond although we cannot rule out that the H-bond is mediated by water (see Extended Data Fig. 7d). Similar chemical shift agreement including for Δω_H3_ was obtained for m^6^_2_A in dsA6RNA (Extended Data Fig. 7b).

Taken together, these data provide strong support for a singly H-bonded m^6^(*syn*)A···U bp (Fig. 4e) which is distinct from the bp open state (Extended Data Fig. 8 and Supplementary Note 2). To our knowledge, this alternative m^6^A-specific conformational state has not been documented previously.

### m^6^(*syn*)A···U behaves like a mismatch

Although we initially dismissed hybridization pathways in which the major *syn* isomer hybridizes to form a dsRNA intermediate, our data indicate that this is indeed possible because m^6^A can pair with uridine to form the m^6^(*syn*)A···U conformation. Several lines of evidence indicate that m^6^(*syn*)A···U behaves like a mismatch when it comes to hybridization kinetics.

Like many mismatches^38^, m^6^(*syn*)A···U loses a H-bond and is destabilized relative to the Watson-Crick m^6^(*anti*)A-U bp by ~3 kcal/mol. In addition, based on the 3-state fit of the RD data measured for dsGGACU^m6A^ at T = 55°C (Fig. 3b), the m^6^(*syn*)A···U containing duplex intermediate anneals at a ~20-fold slower rate compared to the unmethylated control, whereas it melts with an ~80-fold faster rate. These changes in hybridization kinetics relative to the unmethylated control are also in line with those previously reported when introducing single mismatches to dsRNA^22–24^.

We were able to verify the mismatch-like hybridization kinetics of m^6^(*syn*)A···U containing duplex by using NMR RD to measure the hybridization kinetics of the dsGGACU^m62A^ ES-mimic (Extended Data Fig. 7c). For dsGGACU^m62A^, *k*_on_ was ~16-fold slower while *k*_off_ was ~100-fold faster relative to the unmethylated RNA. Therefore, depending on the isomer, m^6^A can behave either like a Watson-Crick (*anti*) or mismatch (*syn*) when paired to the same partner uridine.

### Kinetic model for m^6^A hybridization via conformation selection and induced fit

The RD data measured for dsGGACU^m6A^ at T = 55°C provided direct evidence for hybridization via an IF pathway. Since the RD data measured at T = 65°C is consistent with hybridization via CS, with no evidence for flux along IF, we tested a general model that includes both pathways (CS+IF) (Fig. 5a).

**Fig. 5.**
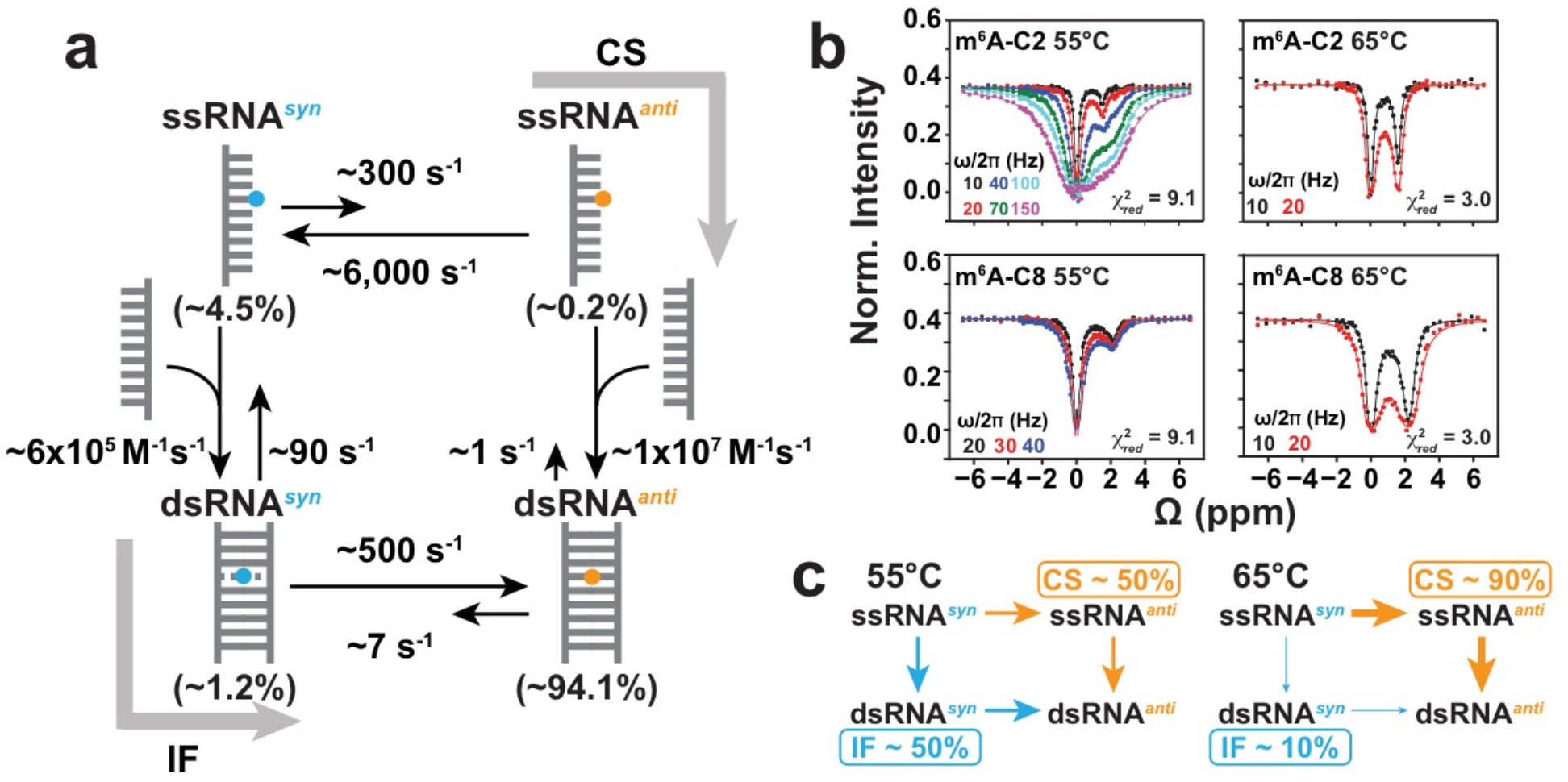
Testing a 4-state CS+IF kinetic model. **a**, Schematic of the CS+IF model with populations and kinetic rate constants measured at T = 55°C for dsGGACU^m6A^. **b**, Constrained 4-state (CS+IF model) shared fit (solid lines) of the m^6^A C2 and C8 ^13^C CEST profiles to the Bloch-McConnell equations for dsGGACU^m6A^ at T = 55°C and 65°C. 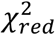 values were obtained from global fitting m^6^A-C2 and m^6^A-C8 CEST data. RF field powers for CEST profiles are color coded. Error bars in CEST profiles (smaller than data points) were obtained using standard deviation of 3 measurements of peak intensity with zero relaxation delay as described in Methods. **c**, Equilibrium flux through CS and IF pathways at T = 55°C and 65°C.

We used the 4-state CS+IF model along with the exchange parameters (Δω, *R*_1_ and *R*_2_ values) determined independently (Methods) to simulate the RD data measured for dsGGACU^m6A^ at T = 55°C. The exchange parameters associated with isomerization in ssRNA were again fixed to the values obtained from temperature dependent RD measurements on ssGGACU^m6A^ (Extended Data Fig. 2c). *k*_*off,anti*_ was again assumed equal to *k*_*off*_ and *k*_*on,anti*_ deduced by using the melting free energy obtained from RD measurements (Methods) (Fig. 2a). *k*_*on,syn*_ and *k*_*off,syn*_ describing the hybridization of ssRNA^*syn*^ and methyl isomerization in dsRNA were fixed to the values obtained from the 3-state fit of the RD data for dsGGACU^m6A^ (Fig. 3b).

Indeed, the RD profiles simulated for m^6^A-C2 using the 4-state model were in much better agreement 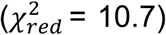 (Extended Data Fig. 9a) with the experimental data relative to simulations using the CS model 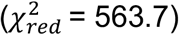 (Extended Data Fig. 4c) or constrained 3-state fits to the CS model 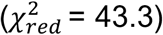 (Fig. 3c). A constrained fit of the RD data to the 4-state model (Methods) improved the agreement further 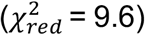 (Extended Data Fig. 9a) to a level comparable to the 3-state fit (Fig. 3a). The 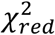 values from globally fitting both m^6^A-C2 and m^6^A-C8 show similar trends (Fig. 5b).

These results provide a plausible explanation for how m^6^A selectively slows dsGGACU^m6A^ annealing at T = 55°C via both the CS and IF pathways. Based on optimized kinetic rate constants obtained from the constrained 4-state fit of the RD data, the flux (Methods) was ~50:50 through the CS and IF pathways at T = 55°C (Fig. 5c). Along the CS pathway, m^6^A reduces the apparent rate of annealing due to the ~20-fold lower population of the ssRNA^*anti*^ intermediate. However, as described for the data measured at T = 65°C, m^6^A does not affect melting because the dominant isomer in the duplex is *anti* which behaves similarly to unmethylated adenine. Along the IF pathway, m^6^A reduces the apparent rate of annealing by 20-fold because m^6^(*syn*)A···U behaves as a mismatch, reducing hybridization rate to form the dsRNA^*syn*^ intermediate by 20-fold. Like a mismatch-containing duplex, this intermediate melts at a rate ~100-fold faster relative to the unmethylated duplex. However, the intermediate does not accelerate the apparent melting rate of the methylated duplex along the IF pathway relative to the unmethylated control because its equilibrium population is only ~1%.

We also re-analyzed the RD data measured at T = 65°C and obtained good agreement with the constrained 4-state fit 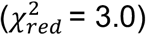 (Fig. 5b). The level of agreement is similar to that obtained using the constrained 3-state fit to the CS model (Fig. 2f), which is expected considering that majority (90%) of the flux is through the CS pathway (Fig. 5c).

### A quantitative model predicts how m^6^A reshapes the hybridization kinetics of DNA and RNA duplexes

To test the generality and robustness of our proposed mechanism, we developed and tested a quantitative CS+IF model that predicts how methylating a central adenine residue impacts the hybridization kinetics for any duplex. The model assumes that the temperature dependent isomerization kinetics in ssRNA and dsRNA do not vary, consistent with the small deviations (<2-fold) seen with sequence, as supported by our data (Supplementary Table 1). The model assumes that *k*_*off,anti*_ = *k*_*off*_ and *k*_*on,anti*_ is deduced based on the known energetics of annealing the m^6^A containing duplex. The value of *k*_*on,syn*_ was assumed to be 20-fold slower than the unmethylated RNA and *k*_*off,syn*_ was then deduced by closing the thermodynamic cycle (Methods). Using these rate constants and the CS+IF model, kinetic simulations (Methods) were used to predict 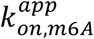 and 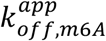.

We used the model to predict the 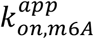 and 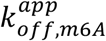 values recently reported^21^ for two duplexes (dsGGACU^m6A^ and dsHCV^m6A^) under a range of different salt (Mg^2+^ and Na^+^) concentrations and temperatures and for a new dataset involving dsHCV^m6A^ at T = 55°C in 3 mM Mg^2+^ (Extended Data Fig. 5). Across these duplexes and conditions, m^6^A slowed the apparent annealing by ~5-fold to ~20-fold while minimally impacting the melting rate (<2-fold). As shown in Fig. 6a, a good correlation (*R*^2^ = 0.8-0.9) was observed between the measured and predicted 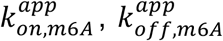, as well as the overall impact on the apparent annealing and melting rates induced by methylation, with all deviations being <1.5-fold.

**Fig. 6.**
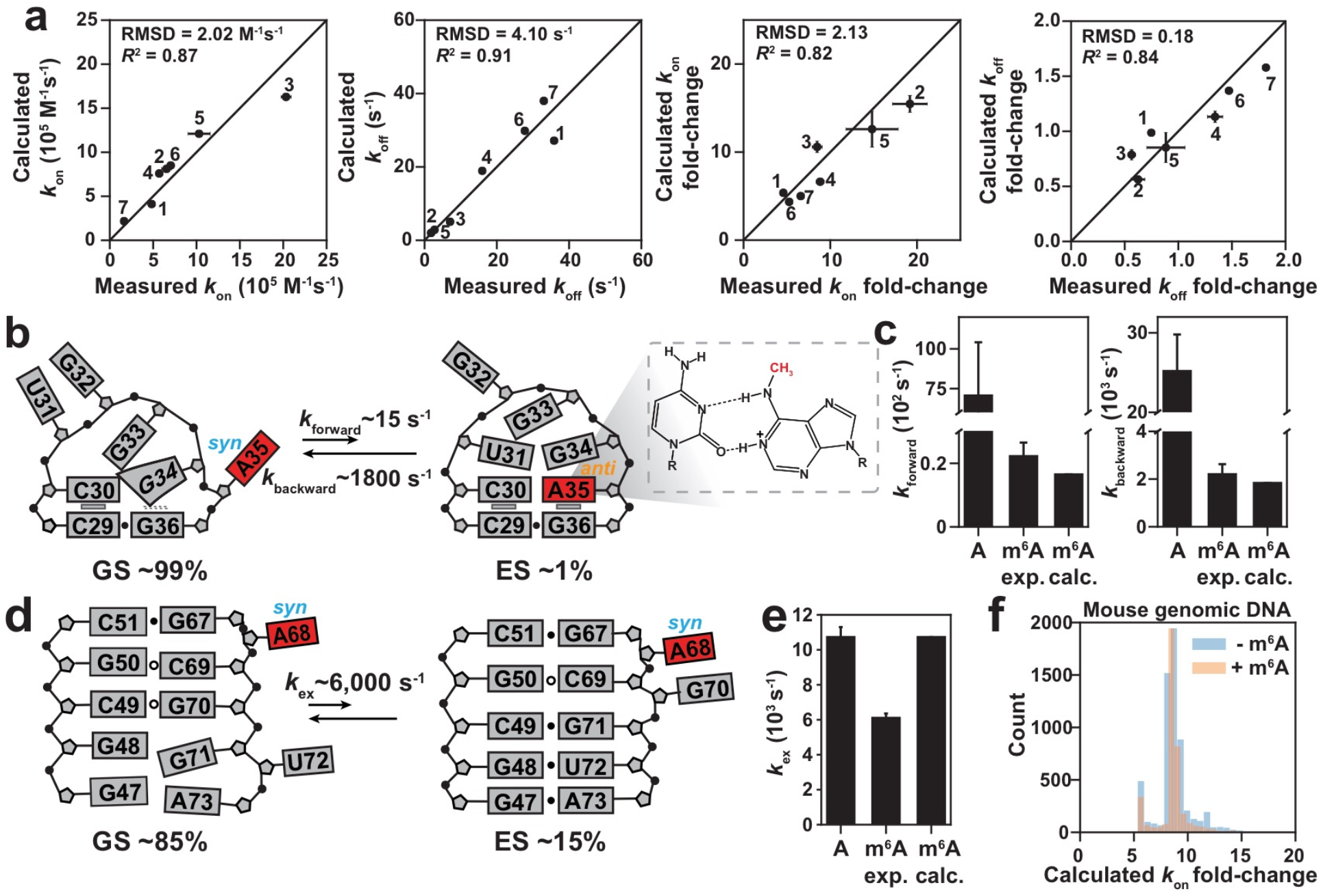
Testing the predictive power of the CS+IF model. **a**, Comparison of experimentally measured and predicted apparent *k*_on_, *k*_off_ and the fold-change relative to unmethylated duplex (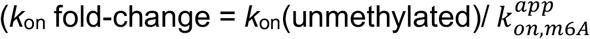 and 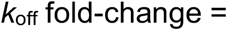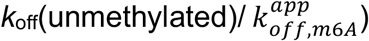) for RNA and DNA duplexes. Each point corresponds to a different duplex and/or experimental condition. All buffers contained 40 mM Na^+^, unless stated otherwise: (1) dsGGACU^m6A^ at T = 65°C, (2) at T = 55°C, (3) with 3 mM Mg^2+^ at T = 65°C; (4) dsHCV^m6A^ with 3 mM Mg^2+^ at T = 60°C, (5) with 3 mM Mg^2+^ at T = 55°C, (6) with 3 mM Mg^2+^ and 100 mM Na^+^ at T = 60°C; (7) dsA6DNA^m6A^ at T = 50°C. Similar correlations were observed using RD simulation-based prediction method shown in Extended Data Fig. 9b. **b**, Secondary structures of GS and ES in the apical loop of HIV-TAR with m^6^A35 (highlighted in red), showing the chemical structure of the m^6^A^+^-C bp. **c**, Comparison of *k*_forward_ and *k_backward_* for unmethylated TAR (A), experimentally measured (m^6^A exp.) and predicted (m^6^A calc.) for methylated TAR. **d**, Secondary structures of GS and ES of methylated RREIIB. **e**, Comparison of *k*_ex_ of unmethylated RRE (A), experimentally measured (m^6^A exp.) and predicted (m^6^A calc.) for methylated RRE. **f**, Predicting the m^6^A-induced slowdown effect on 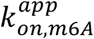 of 12-mers (Methods) for m^6^A sites^9^ (orange) and random DNA (blue) in the mouse genome. Error bars in panel **a**, **c**, **e** were obtained using a Monte-Carlo scheme as described in Methods.

In all the above examples, the equilibrium flux was primarily (~50-95%) via the CS pathway. The differences in the m^6^A induced slowdown (~5-20 fold) of annealing across different duplexes is primarily driven by differences in the annealing rate of ssRNA^*anti*^ along the CS pathway relative to that of unmethylated RNA, with the slowdown being more substantial the more stable the unmethylated duplex (Extended Data Fig. 9c). It should be noted that the slowdown is predicted to be even more substantial when hybridization is fast and isomerization of methylamino group becomes rate-limiting, as observed for an RNA conformational transition, as described below.

As an additional test, we used the model to predict the impact of m^6^A on the apparent hybridization kinetics of an A-rich duplex DNA (dsA6DNA, Extended Data Fig. 5). Based on the unmethylated duplex hybridization kinetics measured previously^21^, the model predicts that m^6^A should reduce the apparent 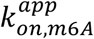 by ~6-fold while having little effect (<2 fold) on 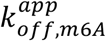. We used NMR RD measurements (Extended Data Fig. 5) on methylated dsA6DNA to test these predictions and the results show that m^6^A reduces 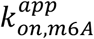 by ~8-fold while having little effect (<2 fold) on 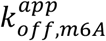, in good agreement with the predictions (Fig. 6a).

Finally, we extended our model to also predict NMR CEST data by imposing additional constraints on NMR exchange parameters (Δω, *R*_1_ and *R*_2_) needed to simulate the RD data (Methods). In addition to providing a rationale for the kinetic basis of the m^6^A induced hybridization slow down, such a model would also validate the existence of the IF and CS intermediates in diverse sequence contexts under a variety of experimental conditions. Thus, we subjected all of the above RD data to a constrained 4-state fit to the CS+IF model. A reasonable fit 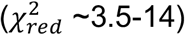 could be obtained in all cases (Extended Data Fig. 9d). This suggests that m^6^A induced hybridization slowdown in DNA is likely mediated by similar IF and CS intermediates as RNA.

### Testing kinetic model on RNA conformational transitions

Beyond duplex hybridization, our kinetic model predicts that m^6^A should also slow intra-molecular conformational dynamics in which m^6^A transitions between an unpaired conformation with the methylamino group predominantly *syn*, to a paired conformation in which the methylamino group is predominantly *anti*. In addition, the model predicts that the slowdown can be much more substantial for conformational transitions that are much faster than the hybridization kinetics measured under our experimental conditions.

To test these predictions, we methylated A35 in the apical loop of transactivation response element (TAR) (Fig. 6b) from human immunodeficiency virus type-1 (HIV-1)^39^ and examined whether m^6^A reduces the rate constant of a previously described intra-molecular conformational transition in which unpaired A35 in the GS forms a wobble A35^+^-C30 mismatch in the ES^40^. As in the Watson-Crick A-U bp, the methylamino group needs to be *anti* to form one of the H-bonds in the m^6^A^+^-C wobble (Fig. 6b). TAR therefore also allowed us to test the generality of the model to non-Watson-Crick bps.

We prepared a TAR NMR sample containing m^6^A35 and ^13^C8-labeled G34 as an RD probe^40^. Based on the chemical shift perturbations, m^6^A destabilized the TAR ES relative to the GS by ~2 kcal/mol, in a manner analogue to duplex destabilization^12^ (Extended Data Fig. 10a, Methods). The CS+IF kinetic model predicts that m^6^A will reduce *k*_ex_, *k*_1_ and *k*_−1_ for the TAR conformational transition by ~17-fold, ~400-fold and ~14-fold respectively. The much greater m^6^A induced reduction in forward rate constant relative to hybridization arises because the TAR conformational transition is intrinsically faster, and this pushes the isomerization step in the dominant CS pathway away from equilibrium, leading to a slowdown much greater than that due to the equilibrium population (~10%) of the ssRNA^*anti*^ CS intermediate when hybridization is limiting. Here, the IF pathway is highly disfavored (flux <1%) because the ES with m^6^A in the *syn* conformation is predicted to be highly energetically disfavored.

Based on NMR RD measurements (Extended Data Fig. 10b), m^6^A reduced *k*_ex_, *k*_1_ and *k*_−1_ by ~15-fold, ~300-fold and ~12-fold in very good agreement with predictions from our model (Fig. 6c). The TAR experimental RD data could be satisfactorily fit to a constrained 3-state fit to the CS model with 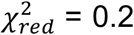 (Extended Data Fig. 10c) comparable to that obtained from an unconstrained 2-state fit. These results indicate that m^6^A can also slowdown RNA conformational transitions and potentially to a much greater degree than observed in our duplex hybridization experiments.

As a negative control, m^6^A minimally (<2-fold) affects exchange rate of conformational transition in the HIV-1 Rev response element stem IIB (RREIIB, Fig. 6d)^41^ in which the m^6^A remains unpaired in the two conformations (Fig. 6e, Extended Data Fig. 10d and Supplementary Note 3).

## Discussion

Our results help explain how m^6^A selectively and robustly slows annealing while minimally impacting the rate of duplex melting under our experimental conditions. The minor ssRNA^*anti*^ isomer hybridizes with kinetic rate constants similar to unmethylated adenine. m^6^A slows the apparent annealing rate along the CS pathway relative to the unmethylated control due to the low equilibrium population of the ssRNA^*anti*^ isomer. Once in a duplex, *anti* is the dominant isomer and m^6^A does not substantially impact the apparent rate of duplex melting along the CS pathway. The major ssRNA^*syn*^ isomer can also hybridize via an IF pathway to form a singly H-bonded bp and with kinetic rate constants similar to that of a mismatch-containing duplex. This intermediate forms slowly, explaining why m^6^A also slows the apparent annealing rate along the IF pathway. However, because its equilibrium population is only ~1%, the intermediate does not accelerate the apparent melting rate along the IF pathway. While we have focused on relatively short duplexes with m^6^A located in the middle, the impact of the modification on the hybridization kinetics will likely vary and be diminished when placed near the terminal ends, as observed for mismatches^22^.

By treating the two m^6^A isomers as two modular elements that have Watson-Crick or mismatch-like kinetic properties independent of sequence context^42^, we were able to build a model that can predict the impact of m^6^A on the overall hybridization kinetics and RNA conformational dynamics from component reactions. The power of such a quantitative and predictive kinetic model is that it obviates the need to carry out time-consuming kinetics experiments to measure the universe of kinetics data that is of biological interest.

For example, when combined with an existing computational model that can predict the hybridization kinetics of unmethylated DNA duplexes from sequence^43^, our model could be used to predict how a central m^6^A impacts the hybridization kinetics of any arbitrary DNA duplex. This allowed us to predict the impact of m^6^A on hybridization kinetics for all ~6,000 m^6^A sites reported in the mouse genome^9^ (Fig. 6f). Our model may also aid the design and implementation of studies which harness the kinetic effects of m^6^A as a chemical tool that can bring conformational transitions within detection or aid kinetic studies of RNA and DNA biochemical mechanisms.

Our model also makes a number of interesting biological predictions. The model predicts that m^6^A should slow any process in which the unpaired m^6^A in the predominantly *syn* isomer has to transition into a conformation in which m^6^A is predominantly *anti*. This should include all templated processes that create canonical A-U Watson-Crick bps and many mismatches (A^+^(*anti*)-C(*anti*), A(*anti*)-G(*anti*), A^+^(*anti*)-G(*syn*)), in which the methylamino group adopts the *anti* conformation. m^6^A is found in a variety of RNAs involved in processes that require base pairing, including R-loop formation^44^, microRNA RNA target recognition^45^, snoRNA-pre-rRNA base pairing^46^, snRNA-pre-mRNA base pairing^47^, and the assembly of the spliceosome^48^ and ribosome^49^. The model also predicts that the m^6^A-induced slowdown could exceed 1000-fold for fast conformational transitions such as the folding of short hairpins and this could have important consequences on RNA folding, conformational switches, RNA protein recognition, and processes that occur co-transcriptionally. Further studies are needed to examine whether m^6^A does indeed slow these processes and whether this has any biological consequences.

The newly uncovered mismatch-like m^6^(*syn*)A···U bp is interesting not only because of its role in hybridization kinetics, but also because it could potentially prime the methylamino group for recognition by reader proteins, which recognize the methylamino group in a *syn* conformation^50^. Upon surveying ~50,000 unmethylated A-U bps in PDB, we found 428 bps that share the conformational signatures of the singly H-bonded m^6^A···U bp (Methods). More than 60% of these bps are found in non-canonical regions, such as junctions, terminal ends, tertiary structural elements, and protein-bound RNA (Extended Data Fig. 7e). It will be interesting to examine whether the mismatch-like m^6^(*syn*)A···U forms as the dominant conformation in certain structural contexts where it may facilitate recognition by reader proteins both by locally destabilizing the bp so that m^6^A is more accessible and by adopting a preformed *syn* conformation.

## Methods

### Sample preparation

#### AMP and m^6^AMP

Unlabeled adenosine and *N*^6^-methyladenosine 5’-monophosphate monohydrate (AMP and m^6^AMP) were purchased from Sigma-Aldrich (A2252 and M2780). Powders were directly dissolved in NMR buffer (25 mM sodium chloride, 15 mM sodium phosphate, 0.1 mM EDTA and 10% D_2_O at pH 6.8 with or without 3 mM Mg^2+^). The final concentrations of AMP and m^6^AMP were 50 mM.

#### Oligonucleotides

Unmethylated, methylated (*N*^6^-methylated adenosine, *N*^6^,*N*^6^-dimethyl adenosine), and ^13^C or ^15^N-site labeled (^15^N3-labeled U, ^13^C8,^13^C2-labeled A/m^6^A and ^13^C10-labeled m^6^A) RNA oligonucleotides were synthesized using a MerMade 6 Oligo Synthesizer employing 2’-tBDSilyl protected phosphoramidites and 1 μmol standard synthesis columns (1000 Å) (BioAutomation). Unlabeled m^6^A, m^6^_2_A, rU and n-acetyl protected rC, rA, rG phosphoramidites were purchased from Chemgenes. ^15^N3-labeled U, ^13^C8,^13^C2-labeled rA/m^6^A phosphoramidites were synthesized in-house according to published procedures^21,51^. ^13^C10-labeled m^6^A phosphoramidite was synthesized as described in Supplementary Note 4. RNA oligonucleotides were synthesized with the option to retain the final 5’-protecting group, 4,4’-dimethoxytrityl (DMT). Synthesized oligonucleotides were cleaved from columns using 1 ml AMA (1:1 ratio of 30% ammonium hydroxide and 30% methylamine) followed by 2-hour incubation at room temperature. The solution was then air-dried and dissolved in 115 μl DMSO, 60 μl TEA, and 75ul TEA.3HF, followed by 2.5 hour incubation at T = 65°C for 2’-O deprotection. The solutions were then quenched using Glen-Pak RNA quenching buffer and loaded onto Glen-Pak RNA cartridges (Glen Research Corporation) for purification and subsequently ethanol precipitated. Following ethanol precipitation, RNA oligonucleotides were dissolved in water (200-500 μM for duplex samples, 50 μM for hairpin samples) and annealed by heating an equimolar amount of complementary single strands or hairpins at T = 95°C for 10 min followed by cooling at room temperature for 2 hours for duplex samples or 30 min on ice for hairpin samples. Extinction coefficients for concentration calculation were obtained from the atdbio online calculator (https://www.atdbio.com/tools/oligo-calculator).

The extinction coefficients for modified single strands were assumed to be equal to that of their unmodified counterparts (modified bases are estimated to affect the extinction coefficient for the oligos used here by <10% based on reference values in Basanta-Sanchez *et al*). All samples were buffer exchanged using centrifugal concentrators (Amicon Ultra-15 3-kDa cut-off EMD Millipore) into NMR buffer (25 mM sodium chloride, 15 mM sodium phosphate, 0.1 mM EDTA and 10% D_2_O at pH 6.8 with or without 3 mM Mg^2+^).

The ^13^C8,^13^C2-labeled m^6^dA ssA6DNA oligonucleotide was synthesized in-house using a MerMade 6 oligo synthesizer. The ^13^C8,^13^C2-labeled m^6^dA phosphoramidite was synthesized as described in Supplementary Note 5. Standard DNA phosphoramidites (n-ibu-dG, bz-dA, ac-dC, dT) were purchased from Chemgenes. DNA oligonucleotides were synthesized with the option to retain the final 5’-DMT group. Synthesized oligonucleotides were cleaved from columns using 1 ml AMA followed by 2-hour incubation at room temperature. The DNA sample were then purified using Glen-Pak DNA cartridges and ethanol precipitated. The complementary ssDNA of the m^6^A containing ssDNA is uniformly ^13^C/^15^N-labeled and was synthesized and purified by *in vitro* primer extension as described previously^21^. DNA duplexes were prepared and buffer exchanged in a manner analogous to that described above for RNA duplexes.

### Definition of rate constants

1. *k*_1_ and *k*_−1_ are the forward and backward rate constants for methylamino isomerization in ssRNA, respectively.
2. *k*_2_ and *k*_−2_ are the forward and backward rate constants for methylamino isomerization in dsRNA, respectively.
3. *k*_*on*_ and *k*_*off*_ are the annealing and melting rate constants, respectively for unmethylated RNA.
4. *k*_*on,anti*_ and *k*_*off,anti*_ are the annealing and melting rate constants, respectively when m^6^A adopts *anti* conformation in both ssRNA and dsRNA.
5. *k*_*on,syn*_ and *k*_*off,syn*_ are the annealing and melting rate constants, respectively when m^6^A adopts *syn* conformation in both ssRNA and dsRNA.
6. 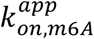 and 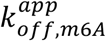 are the apparent annealing and melting rate constants, respectively for m^6^A methylated RNA.
7. *k*_*forward*_ and *k*_*backward*_ are the forward and backward rate constants, respectively for conformational transitions measured using RD.

### NMR experiments

#### Resonance assignments

All NMR experiments (except for the imino proton exchange experiment) were performed on a Bruker Avance III 600 MHz spectrometer equipped with a 5mm triple-resonance HCPN cryogenic probe. Resonance assignments for hpGGACU^m6A^ have been reported previously^51^. Resonance assignments for m^6^_2_A modified dsGGACU and dsA6 were obtained using 2D [^1^H,^1^H] NOESY experiments with 150 ms mixing time along with 2D [^13^C, ^1^H] and [^15^N, ^1^H] HSQC experiments. The assignments for ssGGACU^m6A^, ssA6RNA^m6A^, dsGGACU A/m^6^A, dsA6DNA^m6A^, dsHCV A/m^6^A could be readily obtained since the samples were site-specifically labelled. The assignments for AMP and m^6^AMP were obtained from a prior study^25^ (Extended Data Fig. 1). Data was processed using NMRpipe software package^52^ and analyzed using SPARKY (T.D. Goddard and D.G. Kneller, SPARKY 3, University of California, San Francisco).

#### ^13^C and ^15^N *R*_1ρ_ relaxation dispersion

^13^C and ^15^N *R*_1ρ_ experiments were performed using 1D *R*_1ρ_ schemes as described previously^53–55^. The spin-lock powers (ω/2π Hz) and offsets (Ω_eff_/2π Hz, where Ω_eff_ = ω_obs_ − ω_rf_, where ω_obs_ is the Larmor frequency of the spin and ω_rf_ is the carrier frequency of the applied spin-lock) are listed in Supplementary Table 5. The spin-lock was applied for a maximal duration (<120 ms for ^15^N and <60 ms for ^13^C) to achieve ~70% loss of peak intensity at the end of relaxation delay.

#### Analysis of *R*_1ρ_ data

1D peak intensities were measured using NMRpipe^52^. *R*_1ρ_ values for a given spin-lock power and offset combination were calculated by fitting the intensities at each delay time to a mono-exponential decay as described previously^33^. A Monte-Carlo approach was used to calculate *R*_1ρ_ uncertainties^56^. Alignment of initial magnetization during the Bloch-McConnell fitting was performed based on the *k*_ex_/Δω_major_ ratio as described previously^18^. Chemical exchange parameters were obtained by fitting experimental *R*_1ρ_ values to numerical solutions of the Bloch-McConnell (B-M) equations^57^ that describe n-site chemical exchange^33^. Errors in exchange parameters were determined using a Monte-Carlo approach as described previously^33^. When available, *R*_1ρ_ data measured for the same exchange process under the same condition were globally fitted, sharing ES population and exchange rate constants. Reduced chi-square 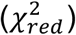 was calculated to assess the goodness of fitting as described previously^18^. In general, similar exchange parameters were obtained from individual fitting and global fitting. All exchange parameters are summarized in Supplementary Table 1.

#### Estimate p_ES_ of methylated TAR from chemical shifts

The RD signal of methylated TAR is weak probably due to small *p*_ES_ and fast *k*_ex_ (Extended Data Fig. 10b). We used chemical shift perturbation based method^58^ as an alternative approach to estimate the population of ES^40^ (*p*_*ES,m6A*_) of methylated TAR. Specifically, in methylated TAR, ω_*obs*_ = ω_*GS*_ × (1 − *p*_*ES,m6A*_) + ω_*ES*_ × *p*_*ES,m6A*_. ω_*GS*_ and ω_*ES*_ are chemical shifts of GS and ES of unmethylated TAR and were determined previously^58^. Based on 2D [^13^C, ^1^H] HSQC spectra, G34-C8 peak shifts towards GS (Extended Data Fig. 10a) and the calculated *p*_*ES,m6A*_ ~1%.

#### ^13^C and ^15^N CEST

^13^C and ^15^N CEST experiments were performed using 1D schemes as described previously without equilibration of GS and ES magnetization prior to the relaxation delay^21^. The radiofrequency (RF) field strengths (ω/2π Hz) and offset combinations (Ω/2π Hz, where Ω = ω_rf_ − ω_obs_) used in CEST measurements are listed in Supplementary Table 6. The relaxation delay for all CEST experiments was 200 ms.

#### 2D CEST for ^13^C methyl probes

The pulse sequence for the ^13^C methyl CEST was derived by modifying the 2D CEST experiment for ^13^C from Zhang *et al*^29^ in accordance with considerations described in Kay *et al*^31^ outlining a 2D CEST experiment for ^13^C methyl groups. The following changes were made to the CEST experiment from Zhang *et al*^29^.

- Given that the samples for methyl CEST in this study were site-specifically ^13^C labeled at the methyl group, we removed shaped pulse c that was used to refocus carbon-carbon scalar couplings.
- The delay τ between ^13^C pulses of phase ϕ2 and ϕ3, and ϕ3 and ϕ5 was set to be as close as possible to the optimal value of 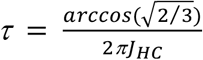 where *J*_HC_ is the scalar coupling between the methyl carbon and protons, for optimal transfer of in-phase methyl carbon magnetization to anti-phase, as described by Kay *et al*., while ensuring that the delays between the pulses in the sequence were positive. *J*_HC_ was measured using an F1 coupled 2D [^13^C, ^1^H] HSQC experiment.
- The τ delay flanking shaped pulse b was set to be equal to 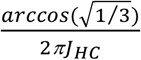. The duration of shaped pulse b was shortened as needed so as to ensure that the delays between the pulses in the sequence were positive.
- A gradient pulse was inserted between the ^13^C and ^1^H π/2 pulses after T1 evolution, as described by Kay *et al*^31^., to purge transverse magnetization.

#### Analysis of the CEST data

1D or 2D peak intensities were calculated using NMRpipe^52^. The intensity error for all offsets for a given spin lock power was set to be equal to the standard deviation of 3 measurements of peak intensity with zero relaxation delay under the same spin lock power. The intensities were normalized to the average intensity of the three zero delay measurements. Exchange parameters were then obtained by fitting experimental intensity values to numerical solutions of the B-M equations and RF field inhomogeneity was taken into account during CEST fitting as described previously^59^. No equilibration of GS magnetization was assumed when integrating the B-M equations for non-methyl probes^59^, while equilibration was assumed for the methyl CEST given that the sequence employs non-selective hard pulses. Fits of CEST data were carried out assuming unequal *R*_2_ or assuming equal *R*_2_ for duplex melting^21^ and other ES measurements, respectively. Alignment of the initial magnetization during CEST fitting was chosen as described previously^59^. Errors in exchange parameters were determined using a Monte-Carlo approach as described previously^60^. Global fitting of CEST data was carried out for the same exchange process under identical condition. 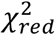 was calculated to assess the goodness of fitting as described previously^18^. Note that the different 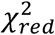 values for different fits are most likely due to differences in the quality of the NMR data and poor estimation of the real experimental uncertainty (Supplementary Table 1). Model selection (3-state with triangular, linear or starlike topology, Extended Data Fig. 4d) was carried out by calculating Akaike’s (wAIC) and Bayesian information criterion (wBIC) weights for each model and selecting the model with the highest relative probability as described previously^33^.

#### ^1^H CEST experiment

A TROSY-based spin-state selective ^1^H CEST experiment^61^ was carried as described previously (Wang *et al*., Chemical Shift Prediction of RNA Imino Groups: Application toward Characterizing RNA Excited States. In revision.). The power of the *B*_1_ field was set to be 60 Hz or 120 Hz and the offset of the *B*_1_ field ranged from 8.5 pm to 15.5 ppm with a step of 30 Hz. The relaxation delay was 400 ms. The ^1^H CEST data were collected in a pseudo-3D mode and were analyzed using NMRPipe^52^. The intensities in the N^α^ and N^β^ CEST profiles were normalized to a reference intensity with *B*_1_ frequency = −20 ppm. The N^β^ CEST profile was then subtracted from the N^α^ CEST profile to result in a difference CEST profile, from which the Δω of the ES was fitted with pre-determined fitting parameters such as *p*_ES_, *k*_ex_, and ^15^N *R*_1_ from the ^13^C/^15^N *R*_1ρ_ experiments. Errors in the CEST intensity profiles were estimated based on the scatter in regions of 1D profiles that did not contain any intensity dips. The Python package *ChemEx* (https://github.com/gbouvignies/chemex) is used to carry out fitting.

#### Imino proton exchange experiment

Experiments were carried out on a 700 MHz Bruker NMR spectrometer quipped with a HCN room-temperature probe to measure the proton exchange between imino proton and water^62^, following the same pulse programs and protocols as described in a prior study^63^. Briefly, the water proton longitudinal relaxation rate constant *R*_1_ was first measured using a standard saturation-recovery method^63^. A pre-saturation pulse was used for solvent suppression. The relaxation delay time for measuring water proton *R*_1_ was set to be 0.0, 0.4, 0.8, 1.2, 1.6, 2.0, 2.4, 2.8, 3.2, 3.6, 4.0, 4.4, 4.8, 5.2, 6.0, 7.0, 8.0, 9.0, 10.0, 12.0 and 15.0 s. The apparent solvent exchange rate constant of the imino protons was then measured using an inversion-recovery scheme by initially selectively inverting the bulk water magnetization followed by detecting transfer of the water magnetization to the imino proton during solvent exchange. A sinc-shaped π-pulse was optimized and used to invert the water magnetization. A binominal water-suppression scheme was used to suppress water. The delay times used to measure water and imino proton exchange rate constants are listed in Supplementary Table 7.

The apparent exchange rate (*k*_*ex*_) of imino and water proton was obtained by fitting the imino magnetization as a function of exchange time upon solvent exchange according to equation (1),

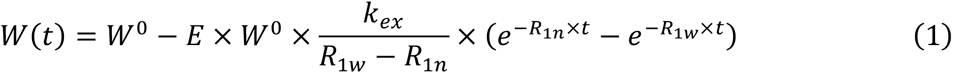

where *W*(*t*) is the imino peak volume as a function of exchange time *t*, *W*^0^ is the initial peak volume (at t = 0 s), *E* is the efficiency of the inversion pulse, *k*_*ex*_ is the apparent solvent exchange rate constant between imino and water proton, *R*_1*w*_ is water proton *R*_1_, *R*_1*n*_ is the summation of imino proton *R*_1_ and exchange rate constant *k*_*ex*_. In the equation, *R*_1*w*_ and *E* values are fixed parameters that are pre-determined, while *k*_*ex*_ and *R*_1*n*_ are fitted parameters. The error of the fitted parameters is the standard fitting error which is the square root of the diagonal elements of the covariance matrix. The efficiency of the selective shape pulse used for water inversion (E) was calculated by the equation (2):

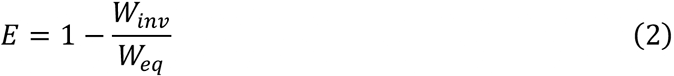

where the *W*_*inv*_ and *W*_*eq*_ represents the peak volumes of the water proton with and without the shape pulse for inversion, respectively (at zero delay time and without binominal water suppression).

#### Determining the methylamino isomerization rate constants from temperature depended RD measurements for methylated ssRNA and dsRNA

The observed temperature dependence of *k*_1_, *k*_−1_ in m^6^AMP and ssRNA (Extended Data Fig. 2c), and *k*_2_, *k*_−2_ in dsRNA (Extended Data Fig. 6d) determined using RD were fit to a modified van’t Hoff equation that accounts for statistical compensation effects and assumes a smooth energy surface as described previously^54^:

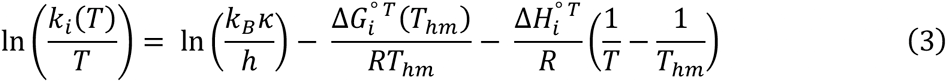

Where *k*_i_ (i = 1, −1 or 2, −2) is the rate constant, 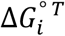 and 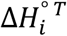 are the free energy and enthalpy of activation (i = 1,2) or deactivation (i = −1, −2) respectively, *R* is the universal gas constant (kcal/mol/K), *T* is temperature (K), and *T*_hm_ is the harmonic mean of the experimental temperatures (*T*_*i*_ in K) computed as 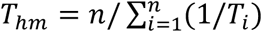, *k*_B_ is the Boltzmann’s constant, *κ* is the transmission coefficient (assumed to be 1). The goodness-of-fit indicator *R*^2^ between the measured and fitted rate constants was calculated as follows: 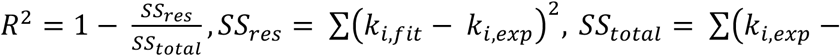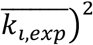. *k*_*i,fit*_ and *k*_*i,exp*_ (i = 1, −1 or 2, −2) are fitted and experimentally measured rate constants. 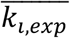 is the mean of all *k*_*i,exp*_. Errors of fitting for 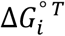 and 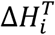 were calculated as the square root of the diagonal elements of the covariance matrix. Given these fitted 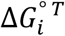 and 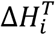 values, *k*_i_ at T = 55°C and 65°C used for kinetic modeling were computed using Equation (3).

#### Measuring the kinetics of duplex hybridization from CEST data

*k*_off_ (s^−1^) and *k*_on_ (M^−1^s^−1^) for duplex hybridization were determined based on the forward rate (*k*_forward_) and backward (*k*_backward_) rate constants obtained from a 2-state fit of the dsHCV/dsHCV^m6A^ A11-C8 and dsA6DNA m^6^A16-C2 RD data (2-state fit of other constructs were reported previously^21^) and 3-state fit of m^6^A-C2 dsGGACU^m6A^ at T = 55°C:

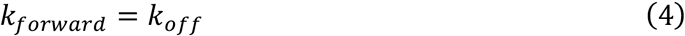

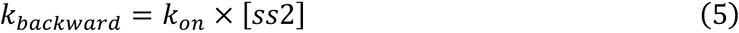

[*ss*2] is the free concentration of the complementary single strand.

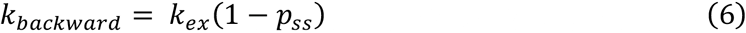

*p*_*ss*_ is the single strand population. The annealing rate constant *k*_on_ is given by:

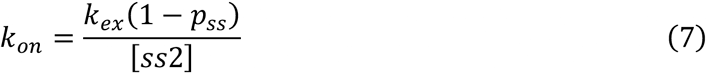

The uncertainty in [*ss*2], and *p*_ss_ and *k*_ex_ from CEST measurements were propagated to determine the uncertainty in of *k*_on_. From 2-state CEST fit, [*ss*2] = *C*_*t*_ × *p*_*ss*_, *C*_*t*_ is the total concentration of the duplex, obtained using the extinction coefficient as described in the ‘Sample preparation’ section. The uncertainty of *C*_*t*_ was assumed to be 20 %^21^. [*ss*2] from a 3-state fit were calculated as described in the energetic decomposition section below.

### UV melting experiments

UV melting experiments were conducted on a PerkinElmer Lambda 25 UV/VIS spectrometer with a RTP 6 Peltier Temperature Programmer and a PCB 1500 Water Peltier System. At least three measurements were carried out for each sample (3 µM in NMR buffer without D_2_O) with a volume of 400 μL in a Teflon-stoppered 1 cm path length quartz cell. The absorbance at 260 nm (A_260_) was monitored at temperatures ranging from 15°C to 95°C, at a ramp rate of 1.0°C/min. The melting temperature (T_m_) and standard enthalpy change (ΔH°) of hybridization reaction for duplexes were obtained by fitting the absorbance of the optical melting experiment to equation (8) and (9)^64^,

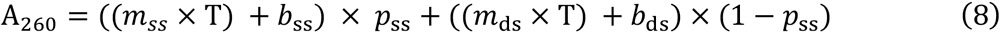

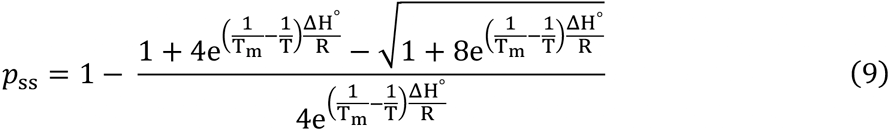

where *m*_*ss*_, *b*_*ss*_, *m*_*ds*_ and *b*_*ds*_ are coefficients describing the temperature dependence of the molar extinction coefficient of single strand and double strands, respectively, T is the temperature (K), *R* is the gas constant (kcal/mol/K) and *p*_ss_ is the population of the single strand. Standard entropy change (ΔS°) and ΔG° of double strand hybridization were therefore computed from equation (10) and (11).

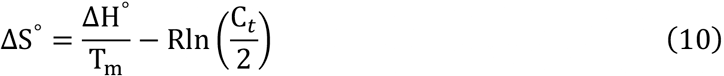

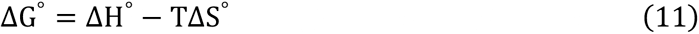

*C*_t_ is the total concentration of duplex. The uncertainty in T_m_ and ΔH° were obtained based on standard deviation in triplicate measurements which were propagated to the uncertainty of ΔS° and ΔG°.

### MD simulations

To generate an ensemble of RNA duplexes with different m^6^A geometries, we performed MD simulations on dsGGACU with the m^6^A-U bp in either *syn* or *anti* conformations, or an m^6^_2_A···U bp. All MD simulations were performed using the ff99 AMBER force field with bsc0 and χ_OL3_ corrections for RNA, using periodic boundary conditions as implemented in the AMBER MD package. Starting structures for MD of unmethylated dsGGACU were generated by building an idealized A-RNA duplex using the *fiber* module of the 3DNA suite of programs^65^. The starting structures for dsGGACU^m6A^ with an m^6^A-U bp in either the *anti* or *syn* conformation were generated by replacing the *anti* and *syn* adenine amino hydrogen atoms in the idealized unmethylated dsGGACU structure with a methyl group. The starting structure for the dsGGACU duplex with the m^6^_2_A-U bp was generated by replacing both of the amino hydrogen atoms of the adenine in the idealized unmethylated dsGGACU structure with methyl groups. All starting structures were solvated with an octahedral box of SPC/E water molecules with box size chosen such that the boundary was at least 10 Å away from any of the DNA atoms. Na^+^ ions treated using the Joung-Cheatham parameters were then added to neutralize the charge of the system. The system was then energy minimized in two stages with the solute heavy atoms (except for the atoms comprising the m^6^_2_A···U bp and the m^6^(*syn*)A···U bp) being fixed (with a restraint of 500 kcal/mol/Å^2^) during the first stage. Heating, equilibration and production runs (500 ns) were performed as described previously^66^. To maintain the methyl group in the *syn* conformation during the MD simulation of the dsGGACU duplex with the m^6^(*syn*)A···U bp, a torsion angle restraint was applied on the angle spanning the methyl carbon-N6-C6-C5 atoms of m^6^A. The restraint was chosen to be square welled between 160° and 200°, parabolic between 159-160° and 200-201°, and linear beyond 201° and less than 159°, with a force constant of 32 kcal/mol/Å^2^. Force field parameters for m^6^A were derived from those in Aduri *et al* ^67^. In particular, the atom types and charges for the methyl group were taken from those by Aduri *et al*, while retaining atom types and charges (apart from N6, see below) for the remaining atoms from those of adenine in the AMBER ff99bsc0χOL3 force field. Charges on the amino N6 atom of m^6^A were adjusted to maintain a net charge for the m^6^A nucleoside of −1. An analogous procedure was followed to generate the parameters for the m^6^_2_A nucleoside. Missing force field parameters were generated using the *antechamber* and *parmchk* utilities of the AMBER suite (16.0).

### Automated fragmentation quantum mechanics/molecular mechanics (AF-QM/MM) chemical shift calculations

We generated mono-nucleoside models of m^6^A with the N1-C6-N6-methyl carbon dihedral angle ranging from 0° to 360° in steps of 20° (*syn* conformation is 0° whereas *anti* conformation is 180°). Coordinates of the m^6^A residue were derived from Aduri *et al* ^67^. We subjected the various mono-nucleoside models and all the RNA duplex MD ensembles (each with N = 100) to QM/MM chemical shift calculations using a fragmentation procedure as described previously^68^. The parameters of geometric minimization for RNA structures were described in a prior study^69^. For all the RNA duplex ensembles, the chemical shift calculations were solely focused on A6 and U13 residues in dsGGACU; therefore, each conformer in the RNA duplex ensembles was broken into only two quantum fragments centered on A6 or U13, respectively, whereas for all the mono-nucleoside models, each quantum fragment was the single mono-nucleoside. We then used a distribution of point charges on the fragment surface to represent the effects of RNA which is outside the quantum fragment and solvent^70^. The local dielectric ∊ value was set to be 1, 4 and 80 for RNA inside quantum fragment, RNA outside quantum fragment and solvent, respectively. We then performed the GIAO chemical shift calculations for each quantum fragment with the OLYP functional and the pcSseg-0 basis set, using demon-2k program (http://www.demon-software.com/public_html/download.html). Reference shieldings were computed for TMS and nitromethane at the same level of theory.

### Free energy decomposition along the CS pathway

The free energy of annealing the methylated duplex can be decomposed into two steps (CS pathway):

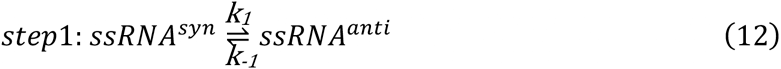

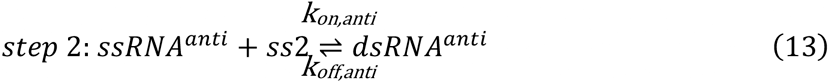

*k*_1_ and *k*_−1_ were determined from 2-state fits or temperature dependence of the RD data (see ‘<u>Determining the methylamino isomerization rate constants from temperature depended RD measurements for methylated ssRNA and dsRNA</u>’ section above):

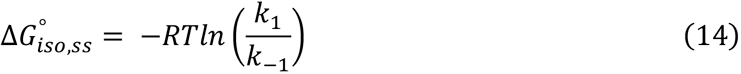

The apparent free energy of annealing methylated dsRNA was determined using:

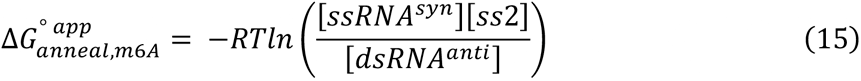

in which the concentrations of the relevant species were measured based on 2-state fits of the RD data^21^:

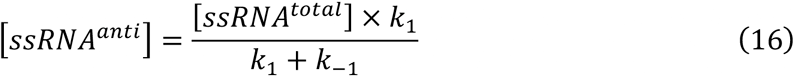

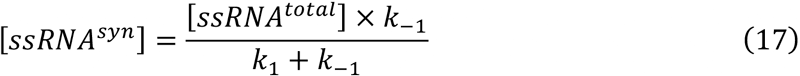

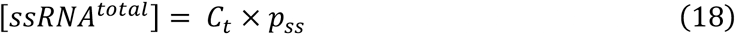

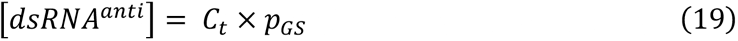

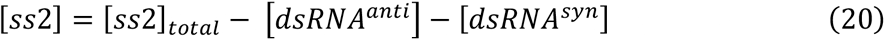

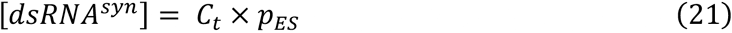

In which *p*_*ss*_ and *p*_*GS*_ are the populations of the ssRNA^total^ (ssRNA^*syn*^+ssRNA^*anti*^) and dsRNA^*anti*^ species obtained from the RD data. [*ss*2]_*total*_ is the total complementary strand concentration. Note that at T = 65°C, dsRNA^*syn*^ has a negligible contribution to RD profiles, [dsRNA^*anti*^] = 0, while at T = 55°C, dsRNA^*syn*^ population (*p*_*ES*_) was obtained from 3-state fit of the m^6^A-C2 CEST data for dsGGACU^m6A^. Also note that the 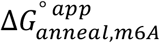 here differs slightly (by ~0.1 kcal/mol) from the prior study^21^, where ssRNA^*syn*^ and ssRNA^*anti*^ were not distinguished.

The free energy of annealing ssRNA^*anti*^ is given by:

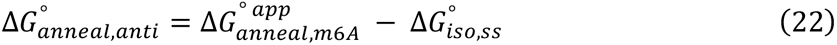

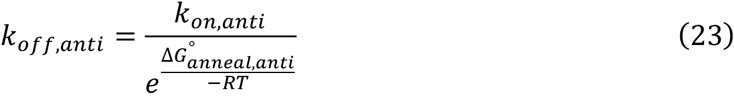

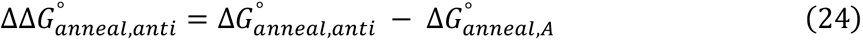

At T = 55°C, 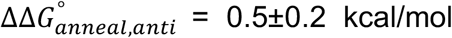, and the m^6^A methyl group in *anti* conformation slightly destabilizes the duplex, whereas it stabilized it by a comparable amount at T = 65°C 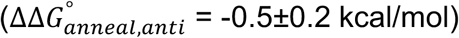.

### B-M simulations and constrained fits

The simulations and constrained fits were performed by numerically integrating the appropriate B-M equations as described previously^33^. Briefly, the simulations were performed by directly predicting RD profiles for a given set of exchange parameters which are defined below. In the constrained fitting, the same numerical integration was used to fit exchange parameters applying specific constraints as detailed below.

#### 3-state CS simulations and constrained fits for the dsGGACU^m6A^ RD data measured at T = 65°C

These analyses used the following input exchange parameters:

1. *k*_1_ and *k*_−1_ were obtained from the temperature dependent RD measurements on ssGGACU^m6A^ (Extended Data Fig. 2c).
2. *k*_*off,anti*_ was assumed equal to *k*_*off*_ measured for the unmethylated dsGGACU and *k*_*on,anti*_ was obtained from the energetic decomposition described above.
3. The longitudinal (*R*_1_) and transverse (*R*_2_) relaxation rate constants for all three species (ssRNA^*syn*^, ssRNA^*anti*^ and dsRNA^*anti*^) were obtained from 2-state fits of the CEST RD data probing duplex melting at T = 65°C^21^. *R*_1_(ssRNA^*anti*^) = *R*_1_(ssRNA^*syn*^) = *R*_1_(dsRNA^*anti*^) = *R*_1,GS_ = *R*_1,ES_. *R*_2_(ssRNA^*anti*^) = *R*_2_(ssRNA^*syn*^) = *R*_2,ES_. *R*_2_(dsRNA^*anti*^) = *R*_2,GS_.
4. The equilibrium populations 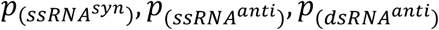 were obtained from kinetic simulations (see differential equations below) that were sufficiently long to ensure equilibration. The same equilibrium populations were obtained from analytical expressions outlined in^71^.

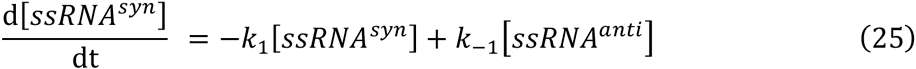

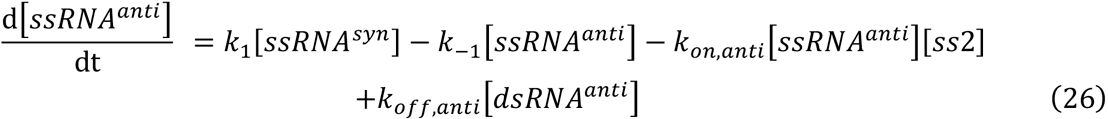

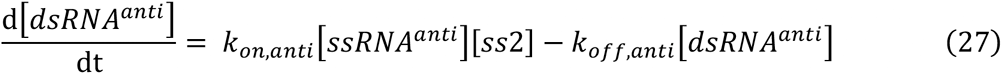

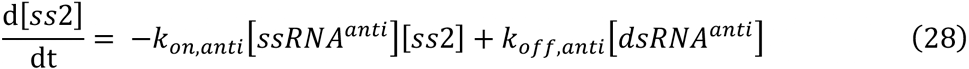
5. Δω of ssRNA^*syn*^ and ssRNA^*anti*^ for C2: 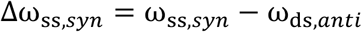, in which 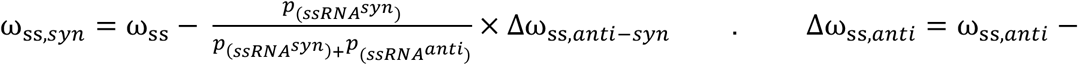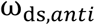, in which 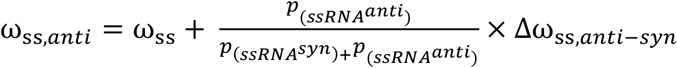. ω_*ss*_ and ω_*ds,anti*_ were obtained from 2D HSQC spectra (Extended Data Fig. 1) and Δω_*ss,anti-syn*_ was obtained from ssGGACU^m6A^ RD measurements at T = 25°C and was assumed to be temperature independent, as supported by the data collected in this study (Extended Data Fig. 2a, Supplementary Table 1). Since C8 is not sensitive to methylamino isomerization (Extended Data Fig. 2), Δω_*ss,anti*_ = 0, while Δω_*ss,syn*_ is obtained from 2-state fit of the CEST RD data probing duplex melting at T = 65°C^21^.

The above parameters were fixed to simulate the CEST profiles using a 3-state Bloch-McConnell equation^33^. For the constrained 3-state fit, the ratio (but not absolute magnitude) of *k*_*on,anti*_ to *k*_*off,anti*_ was constrained to preserve the free energy of the hybridization step. All other parameters (population, *k*_1_, *k*_−1_, Δω, *R*_1_ and *R*_2_ for all species) were allowed to float by an amount determined by the uncertainty (one standard deviation). When possible, global constrained 3-state B-M fits were carried out on both m^6^A C8 and C2 CEST data (Fig. 2f). 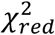 was calculated to assess the goodness of fitting as described previously^18^.

#### 4-state CS+IF simulations and constrained fits for dsGGACU^m6A^ RD data at T = 55°C

These analyses used the following input exchange parameters:

1. All of the exchange parameters related to the CS pathway (*k*_1_, *k*_−1_, *k*_*on,anti*_, *k*_*off,anti*_, Δω_*ss,syn*_, Δω_*ss,anti*_, *R*_1_(ssRNA^*anti*^), *R*_1_(ssRNA^*syn*^), *R*_1_(dsRNA^*anti*^ *R*_2_(ssRNA^*anti*^), *R*_2_(ssRNA^*syn*^) and *R*_2_(dsRNA^*anti*^)) were obtained as described in the previous section for the 3-state CS analysis.
2. *k*_2_, *k*_−2_, *k*_*on,syn*_, *k*_*off,syn*_, C2 Δω_ds,*syn*_, C2 *R*_1_(dsRNA^*syn*^) = *R*_1_(dsRNA^*anti*^) = *R*_1,GS_, and *R*_2_(dsRNA^*syn*^) = *R*_2_(dsRNA^*anti*^) = *R*_2,GS_ were obtained from a 3-state fit to the dsGGACU^m6A^ m^6^A-C2 RD data (Fig. 3a and Supplementary Table 3) using the triangular topology. C8 Δω_ds,_*syn* = 0 because C8 is not sensitive to methylamino isomerization (Extended Data Fig. 6a). C8 *R*_1_(dsRNA^*syn*^) = *R*_1_(dsRNA^*anti*^) = *R*_1,GS_, and *R*_2_(dsRNA^*syn*^) = *R*_2_(dsRNA^*anti*^) = *R*_2,GS_ were obtained from a 2-state fit to the dsGGACU^m6A^ m^6^A-C8 RD data (Supplementary Table 1)
3. The population of all 4 species was obtained from 4-state kinetic simulations using the eight rate constants (*k*_1_, *k*_−1_, *k*_*on,anti*_, *k*_*off,anti*_, *k*_−2_, *k*_2_, *k*_*on,syn*_, *k*_*off,syn*_) based on the CS+IF model (see differential equations below). The same equilibrium populations were obtained from analytical expressions outlined in^71^.

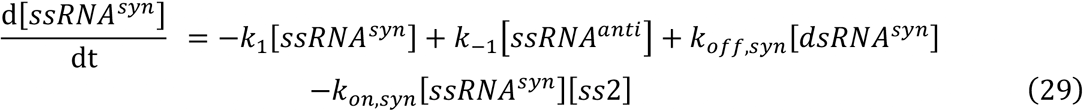

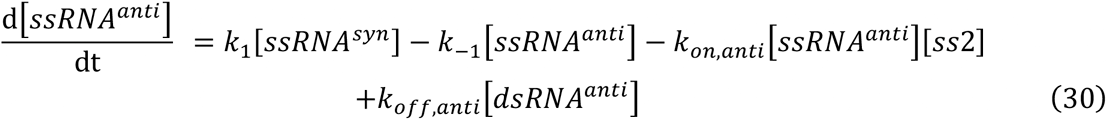

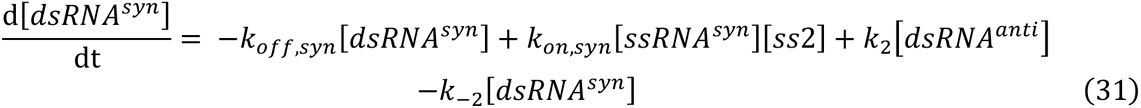

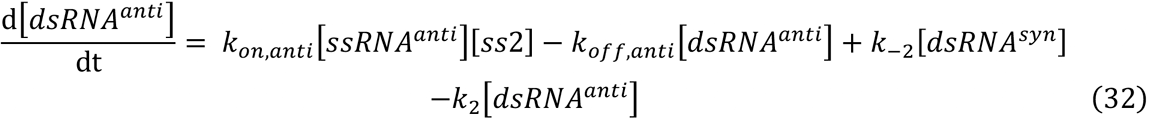

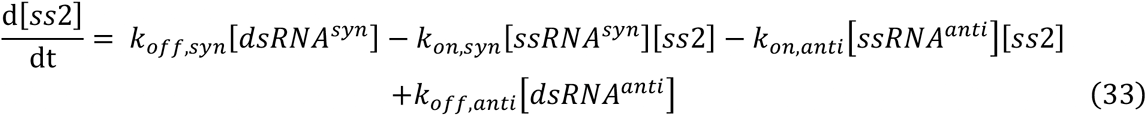

The exchange parameters descried above were then used to simulate CEST profile using a 4-state B-M equation (see below) as described previously^60^:

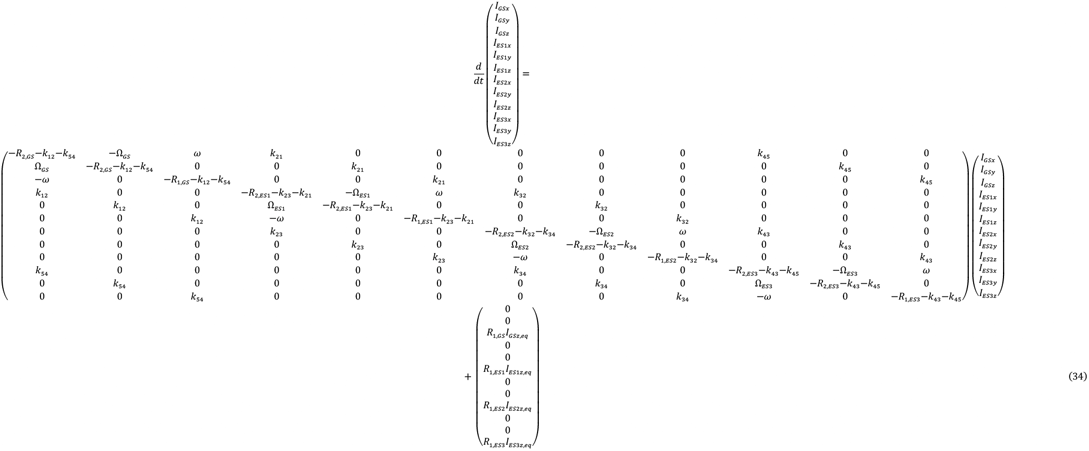

{*GS*/*ES*i}{*x*/*y*/*z*} (i = 1, 2, 3) denotes the magnetization of the GS or ESs in the specified direction. *R*_2,GS_, *R*_2,ES1_, *R*_2,ES2_ and *R*_2,ES3_ are the transverse relaxation rate constants for the GS (dsRNA^*anti*^), ES1 (dsRNA^*syn*^), ES2 (ssRNA^*syn*^) and ES3 (ssRNA^*anti*^) respectively. *R*_1,GS_, *R*_1,ES1_, *R*_1,ES2_ and *R*_1,ES3_ are corresponding longitudinal relaxation rate constants. ω is the RF field power; k_{ij}_ and k_{ji}_ are the forward and backward rate constants of reactions shown in Fig. 5a. Specifically, *k*_12_ = *k*_2_, and *k*_21_ = *k*_−2_ are the forward and backward rate constants of methylamino isomerization in dsRNA. *k*_23_ = *k*_*off,syn*_, *k*32 = *k*_*on,syn*_[*ss*2]. *k*_34_ = *k*_1_ and *k*_43_ = *k*_−1_ are the forward and backward rate constants of methylamino isomerization in ssRNA. *k*_45_ = *k*_*on,anti*_[*ss*2], *k*_54_ = *k*_*off,anti*_. *I*_{*GS/ES*i}z,eq_ (i = 1, 2, 3) denotes the longitudinal magnetization of the GS or ESs at the start of the experiment. Ω_i_ (i = 1, 2, 3, 4) are the offset frequencies of the GS, or ESs resonances in the rotating frame of the RF field, defined as described previously^59^.

We carried out two independent constrained 4-state fits at T = 55°C that differ with regards to how *k*_*on,syn*_ and *k*_*off,syn*_ were defined. In one case, *k*_*on,syn*_ was assumed to be equal to the *k*_*SS→ES*_ rate constant obtained from a 3-state fit to the CEST data measured for dsGGACU^m6A^ at T = 55°C (Fig. 3b) using the triangular topology. Note that this is an approximation since the ssRNA represents the major ssRNA^*syn*^ and minor ssRNA^*anti*^ species in fast exchange. *k*_*off,syn*_ was then calculated by closing the thermodynamic cycle:

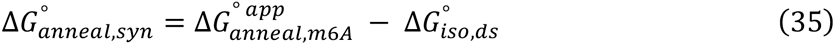

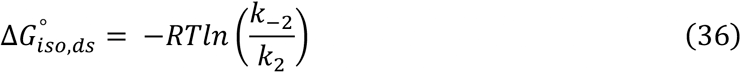

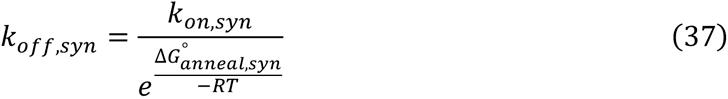

All other exchange parameters were then allowed to float by an amount determined by the experimental uncertainty (one standard deviation). In the second case, only the ratio (but not absolute magnitude) of *k*_*on,syn*_ to *k*_*off,syn*_ was constrained to preserve the free energy of the hybridization step. The fitted *k*_*on,syn*_ and *k*_*off,syn*_ values were similar using these two independent methods. The results from the second method were reported in Fig. 5a and Supplementary Table 2. When possible, global constrained 4-state B-M fits were carried out on both m^6^A C8 and C2 CEST data. 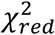 was calculated to assess the goodness of fitting as described previously^18^.

#### 4-state constrained fits for the CS+IF model for dsGGACU^m6A^ at T = 65°C

Because the dsRNA^*syn*^ ES was not directly detected at T = 65°C, the RD data was analyzed as described for T = 55°C with exception that *k*_2_ and *k*_−2_ were measured in hpGGACU^m6A^ at T = 65°C using *R*_1ρ_ RD (Extended Data Fig. 6a), *k*_*on,syn*_ was assumed to be equal to *k*_*on*_/20. This 20-fold slowdown in annealing of ssRNA^*syn*^ relative to unmethylated ssRNA was observed for dsGGACU^m6A^ at T = 55°C. *k*_*off,syn*_ was then calculated by closing the thermodynamic cycle (equations 37). Similar results were obtained when assuming *k*_*off,syn*_ is equal to *k*_*off*_ × 80 as observed for dsGGACU^m6A^ at T = 55°C, and closing the cycle (equations 37) to calculate *k*_*on,syn*_.

#### 4-state constrained fits for the CS+IF model for dsHCV^m6A^ and dsA6DNA^m6A^

RD data measured for dsHCV^m6A^ and dsA6DNA^m6A^ were analyzed in a similar manner as described in the previous sections.

1. *k*_1_, *k*_−1_ and *k*_2_, *k*_−2_ were assumed to be the same as those measured in GGACU^m6A^ constructs using temperature dependent RD measurements (Extended Data Fig. 2c and 6d).
2. *R*_1_(ssRNA^*anti*^) = *R*_1_(ssRNA^*syn*^) = *R*_1_(dsRNA^*anti*^) = *R*_1,GS_ = *R*_1,ES_. *R*_2_(ssRNA^*anti*^) = *R*_2_(ssRNA^*syn*^) = *R*_2,ES_. *R*_2_(dsRNA^*anti*^) = *R*_2,GS_. *R*_1,ES_ and *R*_2,GS_ were obtained from a 2-state fit to the RD data probing duplex melting (Supplementary Table 1).
3. Δω_ss,*anti*_ = Δω_ds,*syn*_ = 0 for A11-C8 in dsHCV^m6A^ since A11 is not the m^6^A site. Δω_ss,*syn*_ was assumed to be equal to the Δω value for A11-C8 in ssRNA obtained from a 2-state fit of the A11-C8 RD data^21^.
4. Δω_ss,*syn*_ and Δω_ss,*anti*_ for m^6^A16-C2 in dsA6DNA^m6A^ were determined as described in CS 3-state simulation for dsGGACU^m6A^ at T = 65°C, assuming Δω_*ss,anti-syn*_ of ssA6DNA^m6A^ is the same as that of ssGGACU^m6A^. Δω_ds,*syn*_ was assumed to be equal to that measured for hpGGACU^m6A^ at T = 55°C (Supplementary Table 1).

### Flux calculations

Flux through the of CS (*F*_CS_) and IF (*F*_IF_) pathways were calculated as the harmonic mean of the forward rates along the CS and IF pathways as described previously^27^:

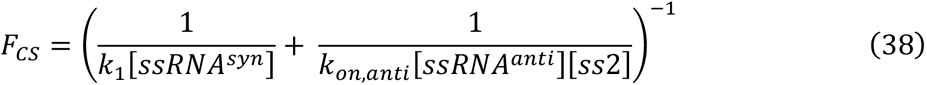

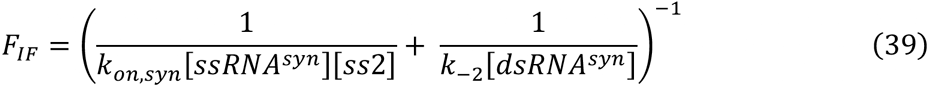

All concentrations are equilibrium concentrations obtained using constrained 4-state fit of CEST data (Fig. 5c) or CS+IF kinetic modeling.

### Model to predict apparent *k*_on_ and *k*_off_ for methylated RNA/DNA duplexes and TAR

The 4-state CS+IF model was used to simulate time traces describing the evolution of all four species as a function of time starting from 100% ssRNA^*syn*^ at t = 0. Similar results were obtained when performing simulations starting with an equilibrium population of ssRNA^*syn*^ (*k*_−1_/(*k*_1_ + *k*_−1_)) and ssRNA^*anti*^ (*k*_1_/(*k*_1_ + *k*_−1_)). *k*_1_, *k*_−1_, *k*_−2_, *k*_2_ were all assumed equal to the corresponding values measured for ssGGACU^m6A^ and dsGGACU^m6A^ at the appropriate temperature based on the temperature dependent RD measurements (Extended Data Fig. 2c and 6d). *k*_*off,anti*_ was assumed to be equal to *k*_*off*_, and *k*_*on,anti*_ was deduced from closing the thermodynamic cycle (equation 23). *k*_*on,syn*_ and *k*_*off,syn*_ were obtained using two different approaches and yielded similar predictions for the apparent *k*_on_ and *k*_off_ for methylated RNA/DNA duplexes and TAR. In one case, *k*_*on,syn*_ = *k*_*on*_/20, and *k*_*off,syn*_ was deduced from closing the thermodynamic cycle (equations 37). Alternatively, *k*_*off,syn*_ = *k*_*off*_ × 80 and *k*_*on,syn*_ was deduced from closing the thermodynamic cycle (equations 37). The predictions shown in Fig. 6a were obtained using the former approach. 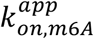 and 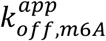 were obtained by fitting simulated time course of [dsRNA^*syn*^] + [dsRNA^*anti*^] at multiple time points to numerical solutions of equation (40) and (41) for a 2-state hybridization model *ss*1 + *ss*2 ⇌ *ds*, 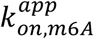 and 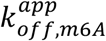 are the annealing and melting constants respectively.

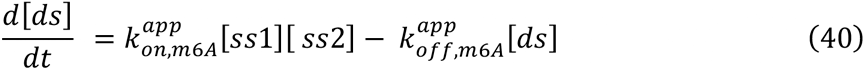

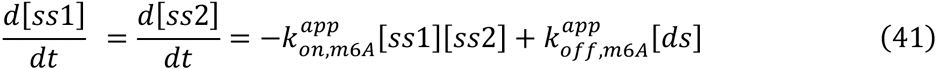

Similar results were obtained when fitting simulated time course of [dsRNA^*anti*^] only. However, it should be noted that for certain kinetic regimes outside those examined here, particularly when *k*_*on,syn*_ is ultra-fast, there can be substantial accumulation of the dsRNA^*syn*^. In this scenario, the system is poorly defined with the apparent 2-state approximation and separate rate constants are needed to describe the evolution of all species. In addition, similar results were obtained from fitting the traces to the appropriate 2-state 2^nd^ order kinetic equation (see ref^72^). Finally, similar results were obtained when simulating m^6^A-C8 RD profiles using 4-state CS+IF model together with exchange parameters (Δω, *R*_1_ and *R*_2_ values for all species) derived from dsGGACU^m6A^ 55°C m^6^A-C8 CEST data, then fitting the data to a 2-state model. Note C8 instead of C2 was used as the probe because the 2-state fit results vary depending on the three Δω values used in C2 CEST simulation. On the other hand, varying the one Δω value used in C8 CEST simulation does not affect the 2-state fit results. As the choice of exchange parameters (*R*_1_ and *R*_2_ values) had a minor effect on the 2-state fit results, we show results from the kinetic simulations in Fig. 6a and that from the 2-state fitting to the simulated C8 RD data in Extended Data Fig. 9b.

A similar approach was used to compute the apparent *k*_*forward*_ and *k*_*backward*_ rate constants for methylated TAR except that *k*_1_, *k*_−1_ were assumed to be equal to the values measured for m^6^AMP, which is a better mimic of the environment of the flipped out and unstacked A35 in TAR than ssRNA. Apparent *k*_*forward*_ and *k*_*backward*_ rate constants were obtained by fitting simulated time course of [*ES*] at multiple time points to the equation 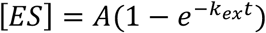, where A is a pre-exponential factor. Note that for the energetics decomposition and kinetic simulations of TAR, the [*SS*2] term in all equations above was removed since the TAR conformational transition is a first order reaction.

### Predict m^6^A-induced slowdown of DNA hybridization in the mouse genome

We used our 4-state CS+IF model to predict the hybridization kinetics for 12-mer DNA duplex representing 5,950 m^6^A sites in the mouse genome^9^ in which m^6^A was positioned at the 6^th^ nucleotide. *k*_*on*_ of unmethylated DNA was predicted as described previously^43^ (http://nablab.rice.edu/nabtools/kinetics.html). The free energy 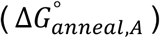 of each sequence was predicted using the MELTING program (https://www.ebi.ac.uk/biomodels-static/tools/melting/). *k*_*off*_ was then deduced by closing the thermodynamic cycle. In all cases, the thermodynamic destabilization of the duplex by m^6^A 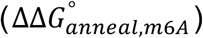 was assumed to be 1 kcal/mol based on prior studies^12,73^ and our measurements (Supplementary Table 4). 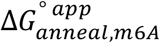 was obtained from 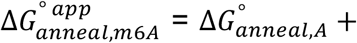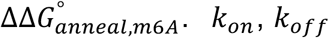 and 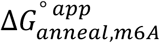 were then used as inputs to predict 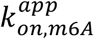 and 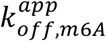 as described in the previous sections. The concentration of dsDNA was assumed to be 1 mM and T = 37°C. We also used this approach to predict the impact of m^6^A on RNA hybridization kinetics at T = 37°C using rate constants for hybridization of unmethylated RNA reported previously^22^ at T = 37°C and assuming that m^6^A destabilizes dsRNA by 1 kcal/mol^12^. m^6^A was predicted to slow *k*_*on*_ by ~5-fold while having a minor effect (<2-fold) on *k*_*off*_, consistent with our measurements at higher temperatures.

### Survey of single H-bonded A-U bps in PDB structures

To identify singly H-bonded A-U bp conformations that mimic the m^6^(*syn*)A···U ES, we conducted a structural survey of the RCSB Protein Data Bank (PDB)^74^. All X-ray (with resolution <= 3.0 Å) and NMR biological assemblies containing RNA molecules (including naked RNA, RNA protein complex etc.) were downloaded from RCSB PDB on Aug 2017 and processed by X3DNA-DSSR^75^ to generate a searchable database containing RNA structural information. Potential candidates of single H-bonded A-U bp were identified by applying the following filters in the database: (1) A-U bps are unmethylated; (2) The Leontis-Westhof (LW) classification^76^ is “cWW”; (3) Both A and U are not in *syn* conformation at glycosidic bond; (4) A-U bps contain A(N1)-U(N3) H-bond (distance between A(N1) and U(N3) is less than 3.5 Å) but do not contain A(N6)-U(O4) H-bond (distance between A(N6) and U(O4) is larger than 3.5 Å). We then manually inspected all the single H-bonded A-U bps, removed misregistered bps, and classified the structure context of all the resulting bps into the following categories (Extended Data Fig. 7e):

1. Junction: A-U bp that is next to an internal bulge, a mismatch or an apical loop.
2. Junction-1/2/3: 1/2/3 bp away from the junction.
3. Tertiary: involved in tertiary interactions.
4. Terminal: at terminal ends.
5. Terminal −1/2/3: 1/2/3 bp away from the terminal end.
6. duplex: A-U bp at canonical duplex context

## Data availability

The data that support this study are contained in the published article (and its Supplementary Information) or are available from the corresponding author on reasonable request.

## Code availability

In-house Python scripts used to perform kinetic simulations and predictions are provided at https://github.com/alhashimilab/m6A_hybridization_kinetics. The force field parameters for m^6^A and m^6^_2_A used in MD simulations and PDB files of these structures that were submitted to the DFT calculations are provided at https://github.com/alhashimilab/m6A_ES.

## Acknowledgements

We thank members of the Al-Hashimi laboratory for assistance and critical comments on the manuscript. We would like to thank Prof. Terrence Oas (Duke University) for advice about kinetic simulations and calculations and Prof. Qi Zhang for providing the 2D [^13^C, ^1^H] CEST pulse sequence based on which the methyl CEST sequence was derived. This work was supported by US National Institute for General Medical Sciences (1R01GM132899) and US National Institute of Health (R01GM089846) to H.M.A., the Austrian Science Fund (FWF, project P30370 and P32773) and the Austrian Research Promotion Agency FFG (West Austrian BioNMR, 858017) to C.K., and the National Institute for Allergy and Infectious Diseases (U54 AI150470) to D.A.C.

## Author contributions

B.L., H.S., and H.M.A. conceived the project and experimental design. B.L. prepared NMR samples, performed NMR experiments, and analyzed NMR data with the help from H.S., A.R. and C.C.C. F.N., K.A.E. and C.K. prepared (^13^CH_3_)-m^6^A RNA phosphoramidite and ^13^C8,^13^C2-labeled m^6^dA phosphoramidite. B.L. performed kinetic simulations and predictions. H.S. performed proton CEST and imino proton exchange experiments. A.R. performed MD simulations. H.S. and D.A.C. performed AF-QM/MM chemical shift calculations. B.L. and H.S. performed the PDB survey. H.M.A. and B.L. wrote the manuscript with critical input from H.S., A.R.

## References

1 Meyer, K. D. et al. Comprehensive analysis of mRNA methylation reveals enrichment in 3’ UTRs and near stop codons. Cell 149, 1635–1646, doi:10.1016/j.cell.2012.05.003 (2012).

2 Dominissini, D. et al. Topology of the human and mouse m6A RNA methylomes revealed by m6A-seq. Nature 485, 201–206, doi:10.1038/nature11112 (2012).

3 Fu, Y., Dominissini, D., Rechavi, G. & He, C. Gene expression regulation mediated through reversible m(6)A RNA methylation. Nat Rev Genet 15, 293–306, doi:10.1038/nrg3724 (2014).

4 Roundtree, I. A., Evans, M. E., Pan, T. & He, C. Dynamic RNA Modifications in Gene Expression Regulation. Cell 169, 1187–1200, doi:10.1016/j.cell.2017.05.045 (2017).

5 Zaccara, S., Ries, R. J. & Jaffrey, S. R. Reading, writing and erasing mRNA methylation. Nat Rev Mol Cell Biol 20, 608–624, doi:10.1038/s41580-019-0168-5 (2019).

6 Vanyushin, B. F., Belozersky, A. N., Kokurina, N. A. & Kadirova, D. X. 5-methylcytosine and 6-methylamino-purine in bacterial DNA. Nature 218, 1066–1067, doi:10.1038/2181066a0 (1968).

7 Douvlataniotis, K., Bensberg, M., Lentini, A., Gylemo, B. & Nestor, C. E. No evidence for DNA N (6)-methyladenine in mammals. Sci Adv 6, eaay3335, doi:10.1126/sciadv.aay3335 (2020).

8 Li, Z. et al. N(6)-methyladenine in DNA antagonizes SATB1 in early development. Nature 583, 625–630, doi:10.1038/s41586-020-2500-9 (2020).

9 Wu, T. P. et al. DNA methylation on N(6)-adenine in mammalian embryonic stem cells. Nature 532, 329–333, doi:10.1038/nature17640 (2016).

10 Liu, N. et al. N(6)-methyladenosine-dependent RNA structural switches regulate RNA-protein interactions. Nature 518, 560–564, doi:10.1038/nature14234 (2015).

11 Huang, L., Ashraf, S., Wang, J. & Lilley, D. M. Control of box C/D snoRNP assembly by N6-methylation of adenine. EMBO Rep 18, 1631–1645, doi:10.15252/embr.201743967 (2017).

12 Roost, C. et al. Structure and thermodynamics of N6-methyladenosine in RNA: a spring-loaded base modification. J Am Chem Soc 137, 2107–2115, doi:10.1021/ja513080v (2015).

13 Choi, J. et al. N(6)-methyladenosine in mRNA disrupts tRNA selection and translation-elongation dynamics. Nat Struct Mol Biol 23, 110–115, doi:10.1038/nsmb.3148 (2016).

14 Slobodin, B. et al. Transcription Impacts the Efficiency of mRNA Translation via Co-transcriptional N6-adenosine Methylation. Cell 169, 326–337 e312, doi:10.1016/j.cell.2017.03.031 (2017).

15 Louloupi, A., Ntini, E., Conrad, T. & Orom, U. A. V. Transient N-6-Methyladenosine Transcriptome Sequencing Reveals a Regulatory Role of m6A in Splicing Efficiency. Cell Rep 23, 3429–3437, doi:10.1016/j.celrep.2018.05.077 (2018).

16 Du, K. et al. Epigenetically modified N(6)-methyladenine inhibits DNA replication by human DNA polymerase eta. DNA Repair (Amst) 78, 81–90, doi:10.1016/j.dnarep.2019.03.015 (2019).

17 Aschenbrenner, J. et al. Engineering of a DNA Polymerase for Direct m(6) A Sequencing. Angew Chem Int Ed Engl 57, 417–421, doi:10.1002/anie.201710209 (2018).

18 Rangadurai, A., Szymaski, E. S., Kimsey, I. J., Shi, H. & Al-Hashimi, H. M. Characterizing micro-to-millisecond chemical exchange in nucleic acids using off-resonance R1rho relaxation dispersion. Prog Nucl Magn Reson Spectrosc 112-113, 55–102, doi:10.1016/j.pnmrs.2019.05.002 (2019).

19 Palmer, A. G., 3rd & Massi, F. Characterization of the dynamics of biomacromolecules using rotating-frame spin relaxation NMR spectroscopy. Chem Rev 106, 1700–1719, doi:10.1021/cr0404287 (2006).

20 Palmer, A. G., 3rd. Chemical exchange in biomacromolecules: past, present, and future. J Magn Reson 241, 3–17, doi:10.1016/j.jmr.2014.01.008 (2014).

21 Shi, H. et al. NMR Chemical Exchange Measurements Reveal That N(6)-Methyladenosine Slows RNA Annealing. J Am Chem Soc 141, 19988–19993, doi:10.1021/jacs.9b10939 (2019).

22 Cisse, II, Kim, H. & Ha, T. A rule of seven in Watson-Crick base-pairing of mismatched sequences. Nat Struct Mol Biol 19, 623–627, doi:10.1038/nsmb.2294 (2012).

23 Xu, S. C. et al. Real-time reliable determination of binding kinetics of DNA hybridization using a multi-channel graphene biosensor. Nature Communications 8, doi:ARTN 1490210.1038/ncomms14902 (2017).

24 Tawa, K. & Knoll, W. Mismatching base-pair dependence of the kinetics of DNA-DNA hybridization studied by surface plasmon fluorescence spectroscopy. Nucleic Acids Research 32, 2372–2377, doi:10.1093/nar/gkh572 (2004).

25 Engel, J. D. & von Hippel, P. H. Effects of methylation on the stability of nucleic acid conformations: studies at the monomer level. Biochemistry 13, 4143–4158 (1974).

26 Engel, J. D. & von Hippel, P. H. Effects of methylation on the stability of nucleic acid conformations. Studies at the polymer level. J Biol Chem 253, 927–934 (1978).

27 Hammes, G. G., Chang, Y. C. & Oas, T. G. Conformational selection or induced fit: a flux description of reaction mechanism. Proc Natl Acad Sci U S A 106, 13737–13741, doi:10.1073/pnas.0907195106 (2009).

28 Sekhar, A. et al. Conserved conformational selection mechanism of Hsp70 chaperone-substrate interactions. Elife 7, doi:10.7554/eLife.32764 (2018).

29 Zhao, B., Hansen, A. L. & Zhang, Q. Characterizing slow chemical exchange in nucleic acids by carbon CEST and low spin-lock field R(1rho) NMR spectroscopy. J Am Chem Soc 136, 20–23, doi:10.1021/ja409835y (2014).

30 Vallurupalli, P., Bouvignies, G. & Kay, L. E. Studying “invisible” excited protein states in slow exchange with a major state conformation. J Am Chem Soc 134, 8148–8161, doi:10.1021/ja3001419 (2012).

31 Bouvignies, G. & Kay, L. E. A 2D (1)(3)C-CEST experiment for studying slowly exchanging protein systems using methyl probes: an application to protein folding. J Biomol NMR 53, 303–310, doi:10.1007/s10858-012-9640-7 (2012).

32 Mulder, F. A., Mittermaier, A., Hon, B., Dahlquist, F. W. & Kay, L. E. Studying excited states of proteins by NMR spectroscopy. Nat Struct Biol 8, 932–935, doi:10.1038/nsb1101-932 (2001).

33 Kimsey, I. J., Petzold, K., Sathyamoorthy, B., Stein, Z. W. & Al-Hashimi, H. M. Visualizing transient Watson-Crick-like mispairs in DNA and RNA duplexes. Nature 519, 315–320, doi:10.1038/nature14227 (2015).

34 Abramov, G., Velyvis, A., Rennella, E., Wong, L. E. & Kay, L. E. A methyl-TROSY approach for NMR studies of high-molecular-weight DNA with application to the nucleosome core particle. Proc Natl Acad Sci U S A 117, 12836–12846, doi:10.1073/pnas.2004317117 (2020).

35 Koss, H., Rance, M. & Palmer, A. G., 3rd. General expressions for R1rho relaxation for N-site chemical exchange and the special case of linear chains. J Magn Reson 274, 36–45, doi:10.1016/j.jmr.2016.10.015 (2017).

36 Bhaswati Goswami, B. L. G., and Roger A. Jones. Nitrogen-15-Labeled Oligodeoxynucleotides. 5. Use of 15N NMR To Probe H-Bonding in an 06MeG-T Base Pair. J. Am. Chem. Soc 115, 3832–3833 (1993).

37 Van Charldorp, R., Heus, H. A. & Van Knippenberg, P. H. Adenosine dimethylation of 16S ribosomal RNA: effect of the methylgroups on local conformational stability as deduced from electrophoretic mobility of RNA fragments in denaturing polyacrylamide gels. Nucleic Acids Res 9, 267–275, doi:10.1093/nar/9.2.267 (1981).

38 Aboul-ela, F., Koh, D., Tinoco, I., Jr. & Martin, F. H. Base-base mismatches. Thermodynamics of double helix formation for dCA3XA3G + dCT3YT3G (X, Y = A,C,G,T). Nucleic Acids Res 13, 4811–4824, doi:10.1093/nar/13.13.4811 (1985).

39 Bannwarth, S. & Gatignol, A. HIV-1 TAR RNA: the target of molecular interactions between the virus and its host. Curr HIV Res 3, 61–71, doi:10.2174/1570162052772924 (2005).

40 Dethoff, E. A., Petzold, K., Chugh, J., Casiano-Negroni, A. & Al-Hashimi, H. M. Visualizing transient low-populated structures of RNA. Nature 491, 724–728, doi:10.1038/nature11498 (2012).

41 Chu, C. C., Plangger, R., Kreutz, C. & Al-Hashimi, H. M. Dynamic ensemble of HIV-1 RRE stem IIB reveals non-native conformations that disrupt the Rev-binding site. Nucleic Acids Res 47, 7105–7117, doi:10.1093/nar/gkz498 (2019).

42 Bisaria, N., Greenfeld, M., Limouse, C., Mabuchi, H. & Herschlag, D. Quantitative tests of a reconstitution model for RNA folding thermodynamics and kinetics. Proc Natl Acad Sci U S A 114, E7688–E7696, doi:10.1073/pnas.1703507114 (2017).

43 Zhang, J. X. et al. Predicting DNA hybridization kinetics from sequence. Nat Chem 10, 91–98, doi:10.1038/nchem.2877 (2018).

44 Abakir, A. et al. N(6)-methyladenosine regulates the stability of RNA:DNA hybrids in human cells. Nat Genet 52, 48–55, doi:10.1038/s41588-019-0549-x (2020).

45 Konno, M. et al. Distinct methylation levels of mature microRNAs in gastrointestinal cancers. Nat Commun 10, 3888, doi:10.1038/s41467-019-11826-1 (2019).

46 Decatur, W. A. & Fournier, M. J. RNA-guided nucleotide modification of ribosomal and other RNAs. J Biol Chem 278, 695–698, doi:10.1074/jbc.R200023200 (2003).

47 Seraphin, B., Kretzner, L. & Rosbash, M. A U1 snRNA:pre-mRNA base pairing interaction is required early in yeast spliceosome assembly but does not uniquely define the 5’ cleavage site. EMBO J 7, 2533–2538 (1988).

48 Will, C. L. & Luhrmann, R. Spliceosome structure and function. Cold Spring Harb Perspect Biol 3, doi:10.1101/cshperspect.a003707 (2011).

49 Klinge, S. & Woolford, J. L., Jr. Ribosome assembly coming into focus. Nat Rev Mol Cell Biol 20, 116–131, doi:10.1038/s41580-018-0078-y (2019).

50 Xu, C. et al. Structural basis for selective binding of m6A RNA by the YTHDC1 YTH domain. Nat Chem Biol 10, 927–929, doi:10.1038/nchembio.1654 (2014).

51 Liu, B. et al. A potentially abundant junctional RNA motif stabilized by m(6)A and Mg(2). Nat Commun 9, 2761, doi:10.1038/s41467-018-05243-z (2018).

52 Delaglio, F. et al. NMRPipe: a multidimensional spectral processing system based on UNIX pipes. J Biomol NMR 6, 277–293, doi:10.1007/BF00197809 (1995).

53 Nikolova, E. N., Gottardo, F. L. & Al-Hashimi, H. M. Probing transient Hoogsteen hydrogen bonds in canonical duplex DNA using NMR relaxation dispersion and single-atom substitution. J Am Chem Soc 134, 3667–3670, doi:10.1021/ja2117816 (2012).

54 Nikolova, E. N. et al. Transient Hoogsteen base pairs in canonical duplex DNA. Nature 470, 498–502, doi:10.1038/nature09775 (2011).

55 Hansen, A. L., Nikolova, E. N., Casiano-Negroni, A. & Al-Hashimi, H. M. Extending the range of microsecond-to-millisecond chemical exchange detected in labeled and unlabeled nucleic acids by selective carbon R(1rho) NMR spectroscopy. J Am Chem Soc 131, 3818–3819, doi:10.1021/ja8091399 (2009).

56 Bothe, J. R., Stein, Z. W. & Al-Hashimi, H. M. Evaluating the uncertainty in exchange parameters determined from off-resonance R1rho relaxation dispersion for systems in fast exchange. J Magn Reson 244, 18–29, doi:10.1016/j.jmr.2014.04.010 (2014).

57 Mcconnell, H. M. Reaction Rates by Nuclear Magnetic Resonance. J Chem Phys 28, 430–431, doi:Doi 10.1063/1.1744152 (1958).

58 Abou Assi, H. et al. 2’-O-Methylation can increase the abundance and lifetime of alternative RNA conformational states. Nucleic Acids Res, doi:10.1093/nar/gkaa928 (2020).

59 Rangadurai, A., Shi, H. & Al-Hashimi, H. M. Extending the Sensitivity of CEST NMR Spectroscopy to Micro-to-Millisecond Dynamics in Nucleic Acids Using High-Power Radio-Frequency Fields. Angew Chem Int Ed Engl 59, 11262–11266, doi:10.1002/anie.202000493 (2020).

60 Vallurupalli, P., Sekhar, A., Yuwen, T. & Kay, L. E. Probing conformational dynamics in biomolecules via chemical exchange saturation transfer: a primer. J Biomol NMR 67, 243–271, doi:10.1007/s10858-017-0099-4 (2017).

61 Yuwen, T. & Kay, L. E. Longitudinal relaxation optimized amide (1)H-CEST experiments for studying slow chemical exchange processes in fully protonated proteins. J. Biomol. NMR 67, 295–307, doi:10.1007/s10858-017-0104-y (2017).

62 Gueron, M., Kochoyan, M. & Leroy, J. L. A single mode of DNA base-pair opening drives imino proton exchange. Nature 328, 89–92, doi:10.1038/328089a0 (1987).

63 Szulik, M. W., Voehler, M. & Stone, M. P. NMR analysis of base-pair opening kinetics in DNA. Curr Protoc Nucleic Acid Chem 59, 7 20 21–18, doi:10.1002/0471142700.nc0720s59 (2014).

64 Bloomfield, V. A. et al. Nucleic Acids: Structure, Properties, and Functions. (University Science Books, 2000).

65 Lu, X. J. & Olson, W. K. 3DNA: a software package for the analysis, rebuilding and visualization of three-dimensional nucleic acid structures. Nucleic acids research 31, 5108–5121, doi:10.1093/nar/gkg680 (2003).

66 Rangadurai, A. et al. Why are Hoogsteen base pairs energetically disfavored in A-RNA compared to B-DNA? Nucleic Acids Res 46, 11099–11114, doi:10.1093/nar/gky885 (2018).

67 Aduri, R. et al. AMBER Force Field Parameters for the Naturally Occurring Modified Nucleosides in RNA. J Chem Theory Comput 3, 1464–1475, doi:10.1021/ct600329w (2007).

68 Swails, J., Zhu, T., He, X. & Case, D. A. AFNMR: automated fragmentation quantum mechanical calculation of NMR chemical shifts for biomolecules. J Biomol NMR 63, 125–139, doi:10.1007/s10858-015-9970-3 (2015).

69 Shi, H. et al. Rapid and accurate determination of atomistic RNA dynamic ensemble models using NMR and structure prediction. Nat Commun 11, 5531, doi:10.1038/s41467-020-19371-y (2020).

70 Richardson, W. H., Peng, C., Bashford, D., Noodleman, L. & Case, D. A. Incorporating solvation effects into density functional theory: Calculation of absolute acidities. Int J Quantum Chem 61, 207–217, doi:Doi 10.1002/(Sici)1097-461x(1997)61:2<207::Aid-Qua3>3.3.Co;2-4 (1997).

71 Orlovsky, N. I., Al-Hashimi, H. M. & Oas, T. G. Exposing Hidden High-Affinity RNA Conformational States. J Am Chem Soc 142, 907–921, doi:10.1021/jacs.9b10535 (2020).

72 Meagher, N. E. & Rorabacher, D. B. Mathematical Treatment for Very Rapid 2nd-Order Reversible Kinetics as Measured by Stopped-Flow Spectrophotometry with Corrections for the Cell Concentration Gradient. J Phys Chem-Us 98, 12590–12593, doi:DOI 10.1021/j100099a022 (1994).

73 Guo, Q., Lu, M. & Kallenbach, N. R. Effect of hemimethylation and methylation of adenine on the structure and stability of model DNA duplexes. Biochemistry 34, 16359–16364, doi:10.1021/bi00050a016 (1995).

74 Berman, H. M. et al. The Protein Data Bank. Nucleic Acids Res 28, 235–242, doi:10.1093/nar/28.1.235 (2000).

75 Lu, X. J., Bussemaker, H. J. & Olson, W. K. DSSR: an integrated software tool for dissecting the spatial structure of RNA. Nucleic Acids Res 43, e142, doi:10.1093/nar/gkv716 (2015).

76 Leontis, N. B. & Westhof, E. Geometric nomenclature and classification of RNA base pairs. RNA 7, 499–512, doi:10.1017/s1355838201002515 (2001).

