## Supplementary Information for "A quantitative model predicts how m^6^A reshapes the kinetic landscape of nucleic acid hybridization and conformational transitions"

### Extended Data Figures

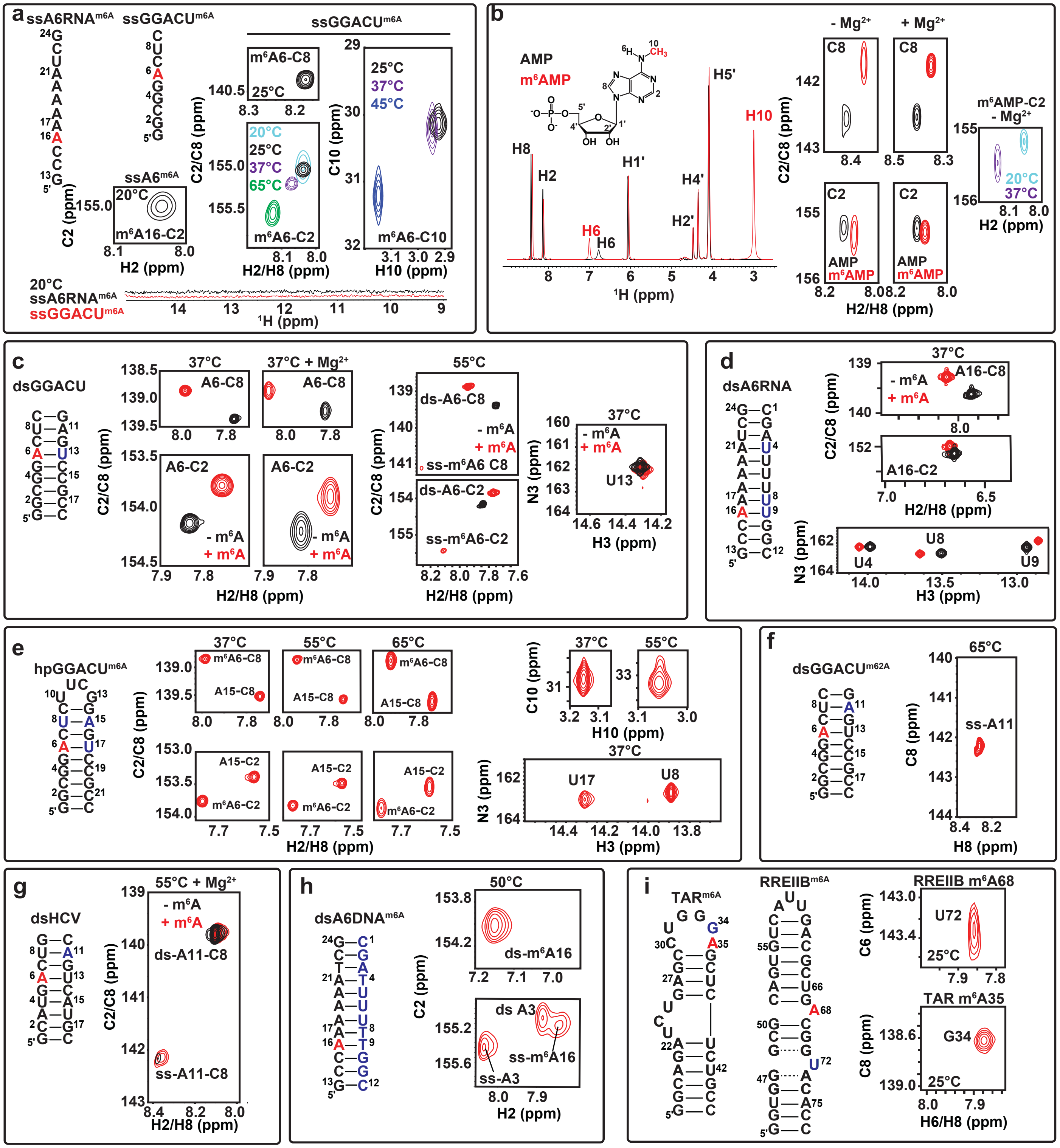


**Extended Data Fig. 1. NMR spectra of RNA and DNA constructs used for RD measurements.** m$^6$A/m$^6_2$A sites are in red and other $^{13}$C/$^{15}$N labeled sites are in blue. Shown are 1D and/or 2D HSQC spectra of **a**, $^{13}$C site-labeled (m$^6$A6-C2/C8 or C10) ssA6RNA$^{m6A}$ and ssGGACU$^{m6A}$; **b**, Unlabeled AMP and m$^6$AMP; **c**, $^{13}$C site-labeled (m$^6$A6/A6-C2/C8) and $^{15}$N site-labeled (U13-N3) dsGGACU with and without m$^6$A; **d**, $^{13}$C site-labeled (m$^6$A16/A16-C2/C8) and $^{15}$N site-labeled (U4-N3, U8-N3 and U9-N3)

dsA6RNA with and without $m^6A$; **e**, $^{13}C$ site-labeled ($m^6A6$-C2/C8, A15-C2/C8) and $^{15}N$ site-labeled (U17-N3 and U8-N3) or only $m^6A6$-C10 $^{13}C$ site-labeled $hpGGACU^{m6A}$; **f**, $^{13}C$ site-labeled (A11-C2/C8) $m^6{}_2A$ modified $dsGGACU^{m62A}$; **g**, $^{13}C$ site-labeled (A11-C2/C8) dsHCV with and without unlabeled $m^6A6$; **h**, $m^6A$ containing strand $^{13}C$ site-labeled ($m^6A6$-C2/C8) and the uniformly $^{13}C/^{15}N$ labeled complementary strand of $dsA6DNA^{m6A}$; **i**, $^{13}C$ site-labeled (G34-C8 for TAR and U72-C6 for RREIIB) methylated HIV-1 TAR and RREIIB. The buffers used for NMR measurements contain 15 mM sodium phosphate, 25 mM sodium chloride, 0.1 mM EDTA at pH 6.8 in a 90% $H_2O$:10% $D_2O$ mixture. +$Mg^{2+}$ corresponds to 3 mM $Mg^{2+}$ in the buffer.

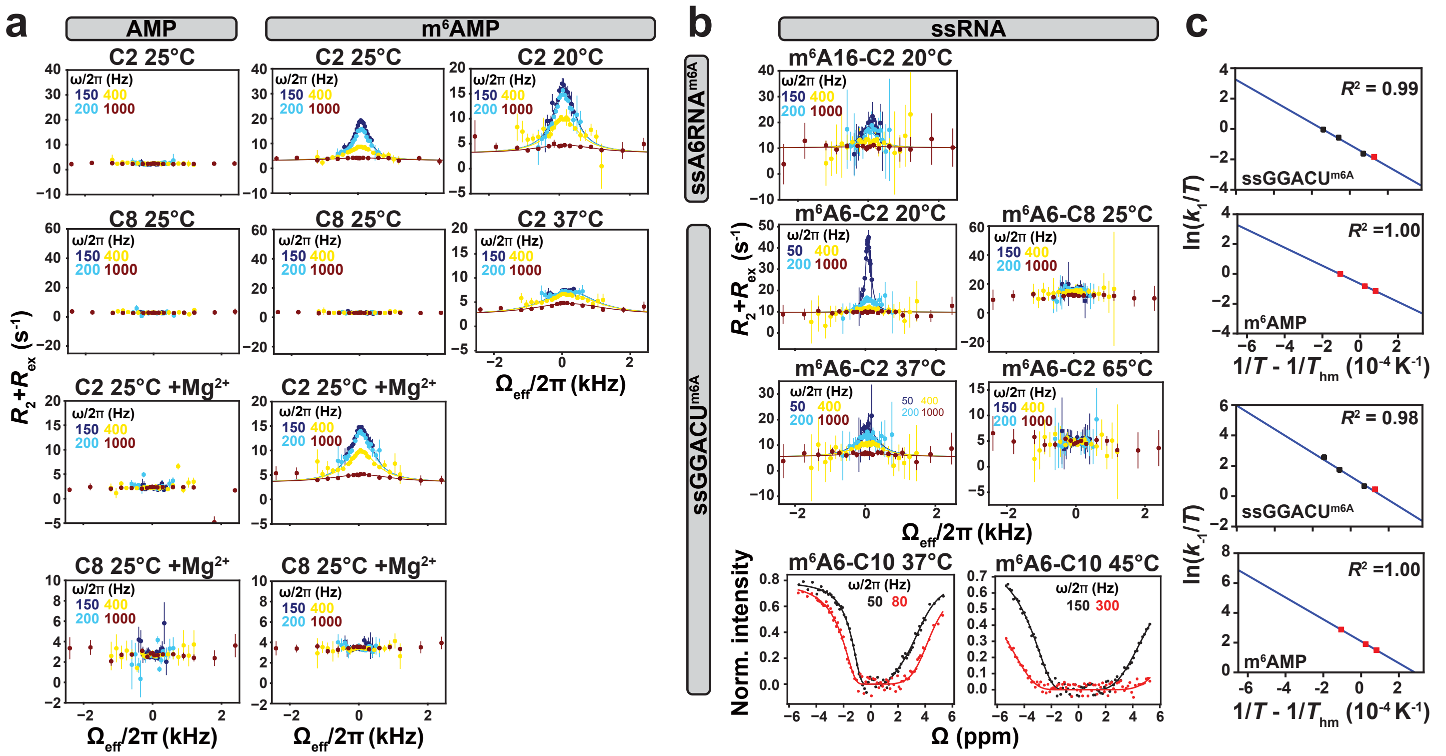


**Extended Data Fig. 2. RD measurements of methylamino isomerization in unpaired RNA.** **a**, $R_{1\rho}$ RD profiles for AMP (negative control) and m$^6$AMP at T = 20°C-37°C in the absence or presence of 3 mM $Mg^{2+}$. **b**, $R_{1\rho}$ RD and CEST profiles for ssA6RNA$^{m6A}$ and ssGGACU$^{m6A}$ at T = 20°C-65°C. **c**, van't Hoff plot showing temperature dependence of the forward ($k_1$) and reverse ($k_{-1}$) rate constants for methylamino isomerization in ssGGACU$^{m6A}$ and m$^6$AMP. Data points in black and red were measured using C10 CEST and C2 $R_{1\rho}$, respectively. $R^2$ denotes coefficient of determination (Methods). RF powers for CEST and spin-lock powers for $R_{1\rho}$ are color coded. Solid lines denote fits of the data to the Bloch-McConnell equations (Methods). Error bars for CEST profiles (smaller than data points) were obtained using standard deviation of 3 measurements of peak intensity with zero relaxation delay as described in Methods. Error bars for $R_{1\rho}$ profiles were obtained using Monte-Carlo simulations as described in Methods. Error bars in panel **c** were determined by propagating the error in $k_1$ and $k_{-1}$ obtained from CEST and $R_{1\rho}$ experiments. Buffer used for RD measurements was composed of 15 mM sodium phosphate, 25 mM sodium chloride, 0.1 mM EDTA in a 90% $H_2O$:10% $D_2O$ mixture at pH 6.8. +$Mg^{2+}$ corresponds to 3 mM $Mg^{2+}$ in the buffer.

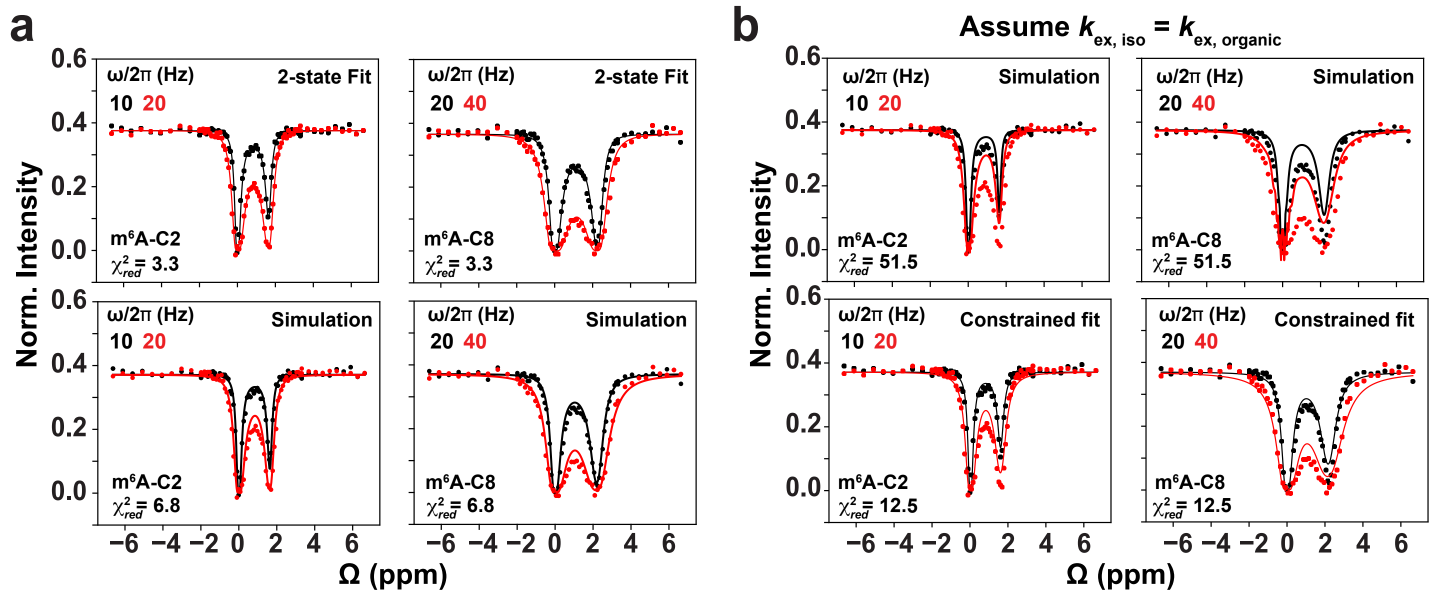


**Extended Data Fig. 3. Kinetic modelling of CS pathway at T = 65°C.** **a**, Comparison of experimental m$^6$A6 C2/C8 CEST data points measured for dsGGACU$^{m6A}$ with a line computed from a 2-state fit to the RD data (2-state Fit) and from 3-state simulations of the CS pathway (Simulation) using the Bloch-McConnell equations, without any adjustable parameters when assuming the methylamino isomerization kinetics measured for ssGGACU$^{m6A}$. **b**, Results for the 3-state CS simulation and constrained 3-state fit to the CS pathway when decreasing the rate of exchange for methylamino isomerization ($k_{ex,iso}$) in ssRNA by 20-fold to mimic results from prior measurements[1] on the nucleobase in organic solvent ($k_{ex,organic}$ = 500 s$^{-1}$). Error bars for CEST profiles (smaller than data points) were obtained using standard deviation of 3 measurements of peak intensity with zero relaxation delay as described in Methods. RF field powers used for CEST are color-coded.

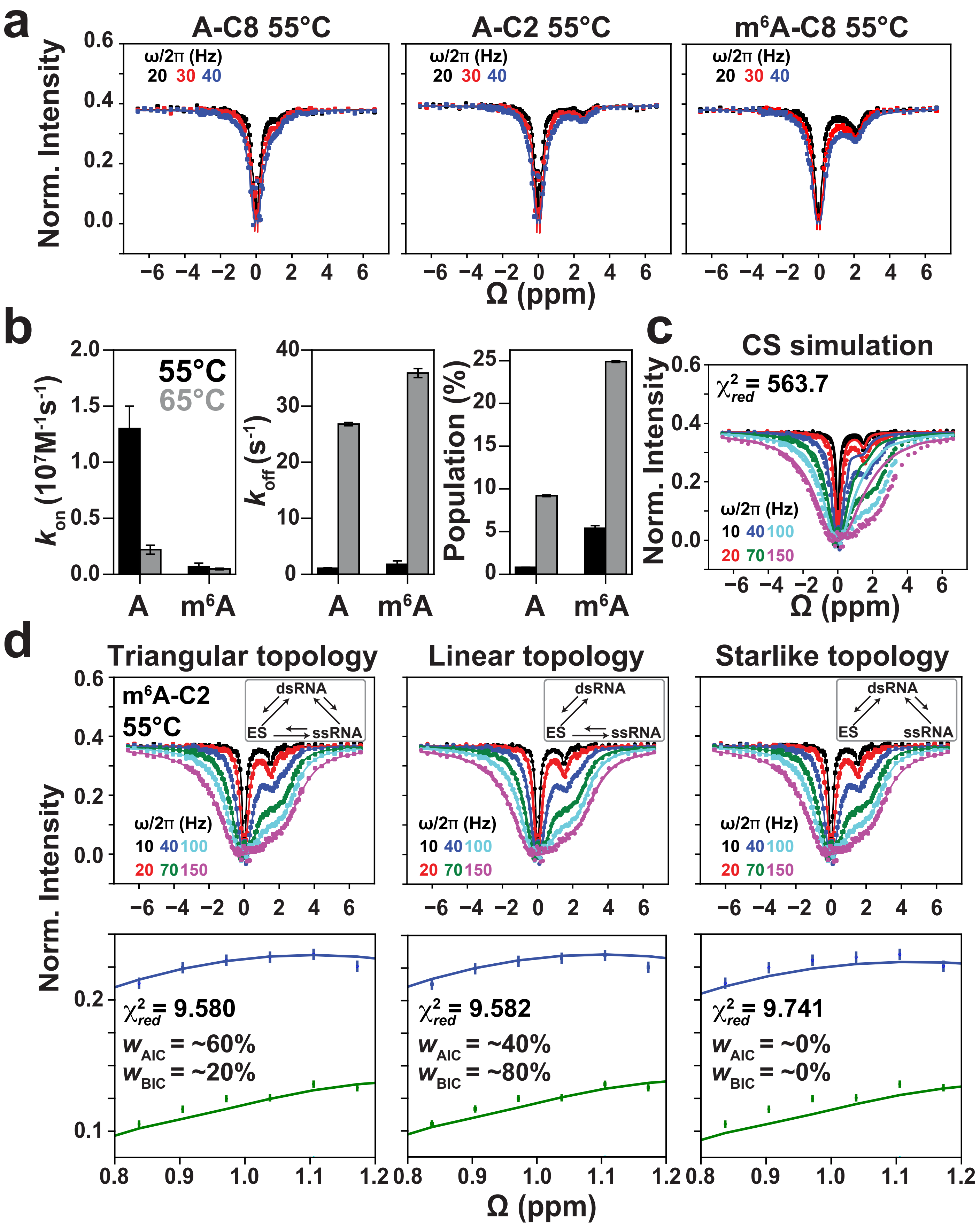
a
A-C8 55°C
A-C2 55°C
m6A-C8 55°C
ω/2π (Hz)
20 30 40
Norm. Intensity
Ω (ppm)
b
55°C
65°C
$k_{on}$ ($10^7 M^{-1}s^{-1}$)
$k_{off}$ ($s^{-1}$)
Population (%)
A
m6A
c
CS simulation
$\chi^2_{red}$ = 563.7
ω/2π (Hz)
10 40 100
20 70 150
Ω (ppm)
d
Triangular topology
Linear topology
Starlike topology
m6A-C2
55°C
dsRNA
ES
ssRNA
$\chi^2_{red}$ = 9.580
$w_{AIC}$ = ~60%
$w_{BIC}$ = ~20%
$\chi^2_{red}$ = 9.582
$w_{AIC}$ = ~40%
$w_{BIC}$ = ~80%
$\chi^2_{red}$ = 9.741
$w_{AIC}$ = ~0%
$w_{BIC}$ = ~0%
Ω (ppm)

**Extended Data Fig. 4. CEST measurements for dsGGACU with and without m$^6$A at T = 55°C. a**, $^{13}$C CEST profiles for A6 C2/C8 (left and middle) and m$^6$A6 C8 (right) in dsGGACU and dsGGACU$^{m6A6}$ respectively at T = 55°C. **b**, Fitted $k_{on}$, $k_{off}$ for dsGGACU, apparent $k_{on}$ ($k_{on,m6A}^{app}$), $k_{off}$ ($k_{off,m6A}^{app}$) for dsGGACU$^{m6A6}$, and ssRNA populations from 2-state fits of dsGGACU and dsGGACU$^{m6A6}$ CEST data at T = 55°C and 65°C (the 65°C data was obtained from a prior study[2]). **c**, 3-state CS simulation (solid lines) versus experimental CEST data points for dsGGACU$^{m6A6}$ m$^6$A-C2 at T = 55°C. **d**, Unconstrained 3-state fit using triangular topology (left), linear topology (middle) or starlike topology (right) for the m$^6$A6 C2 $^{13}$C CEST profile of dsGGACU$^{m6A6}$ at T = 55°C. $\chi^2_{red}$ and AIC/BIC weights were used to select the best model. The AIC and BIC analysis rejects the starlike topology. Both the full (top) and zoomed-in (bottom) profiles are shown. Solid lines denote fits or simulations performed using the Bloch-McConnell equations as described in Methods. RF field powers for CEST profiles are color coded, while the error bars for CEST profiles (smaller than data points) were obtained using standard deviation of 3 measurements of peak intensity with zero relaxation delay as described in Methods. Buffer used for RD measurements was composed of 15 mM sodium phosphate, 25 mM sodium chloride, 0.1 mM EDTA in a 90% $H_2O$:10% $D_2O$ mixture at pH 6.8.

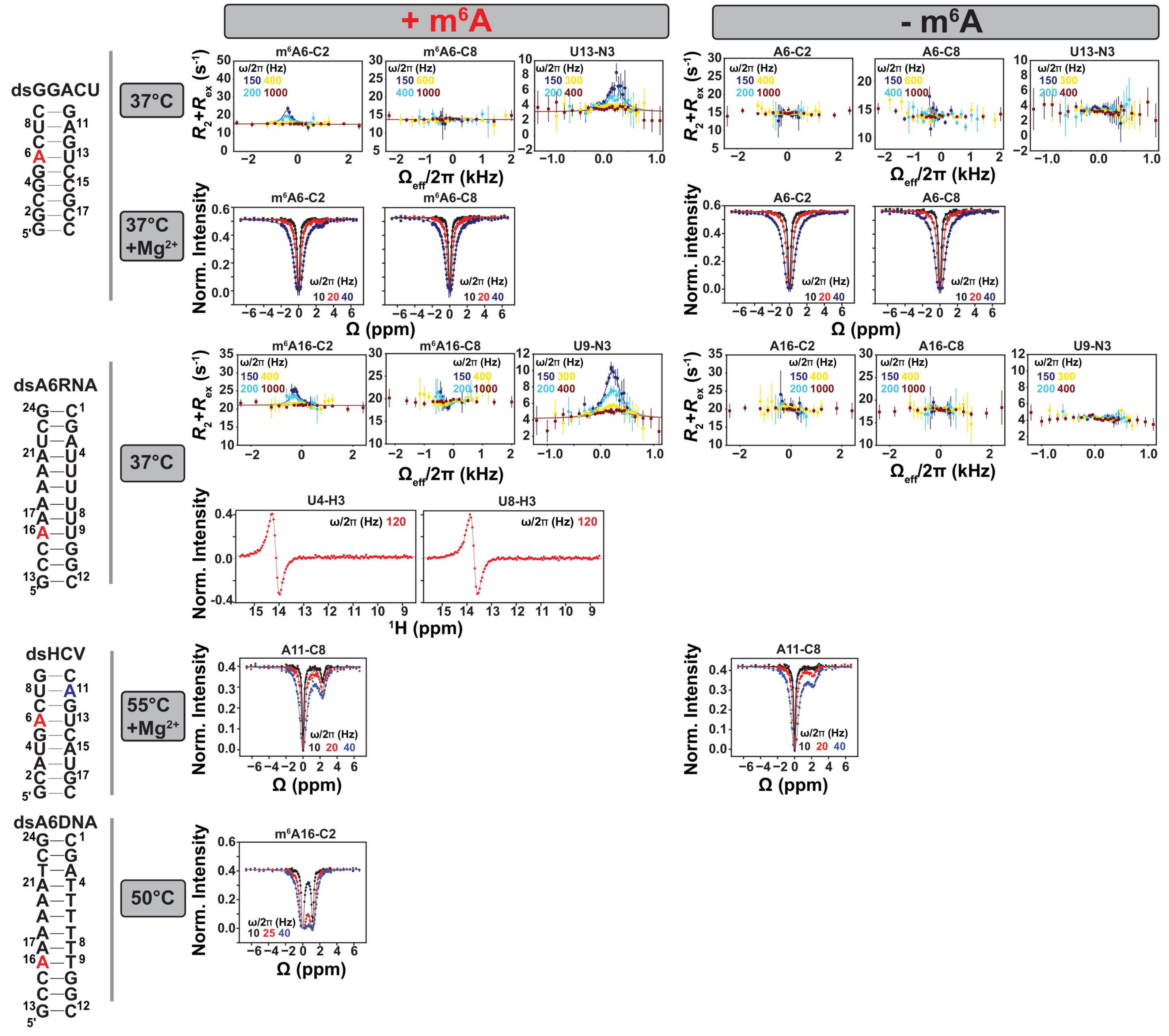


**Extended Data Fig. 5. $R_{1\rho}$ and CEST profiles for four RNA duplexes under different conditions.** These RD data probe the new ES (m$^6$(*syn*)A···U) in dsGGACU$^{m6A}$ and dsA6RNA$^{m6A}$, and hybridization kinetics in dsHCV/dsHCV$^{m6A}$ and dsDNA$^{m6A}$. Shown are various duplex constructs with m$^6$A sites highlighted in red, experimental conditions, and the corresponding RD data. The labeling scheme of constructs were described in Extended Data Fig. 1. Solid lines denote a fit to the Bloch-McConnell equations as described in Methods. Spin-lock powers for $R_{1\rho}$ and RF field powers for CEST profiles are color coded. Error bars for CEST profiles (smaller than data points) were obtained using standard deviation of 3 measurements of peak intensity with zero relaxation delay as described in Methods. Error bars for $R_{1\rho}$ profiles were obtained using Monte-Carlo simulations as described in Methods. Buffer used for RD measurements was composed of 15 mM sodium phosphate, 25 mM sodium chloride, 0.1 mM EDTA in a 90% $H_2O$:10% $D_2O$ mixture at pH 6.8. +$Mg^{2+}$ corresponds to 3 mM $Mg^{2+}$ in the buffer.

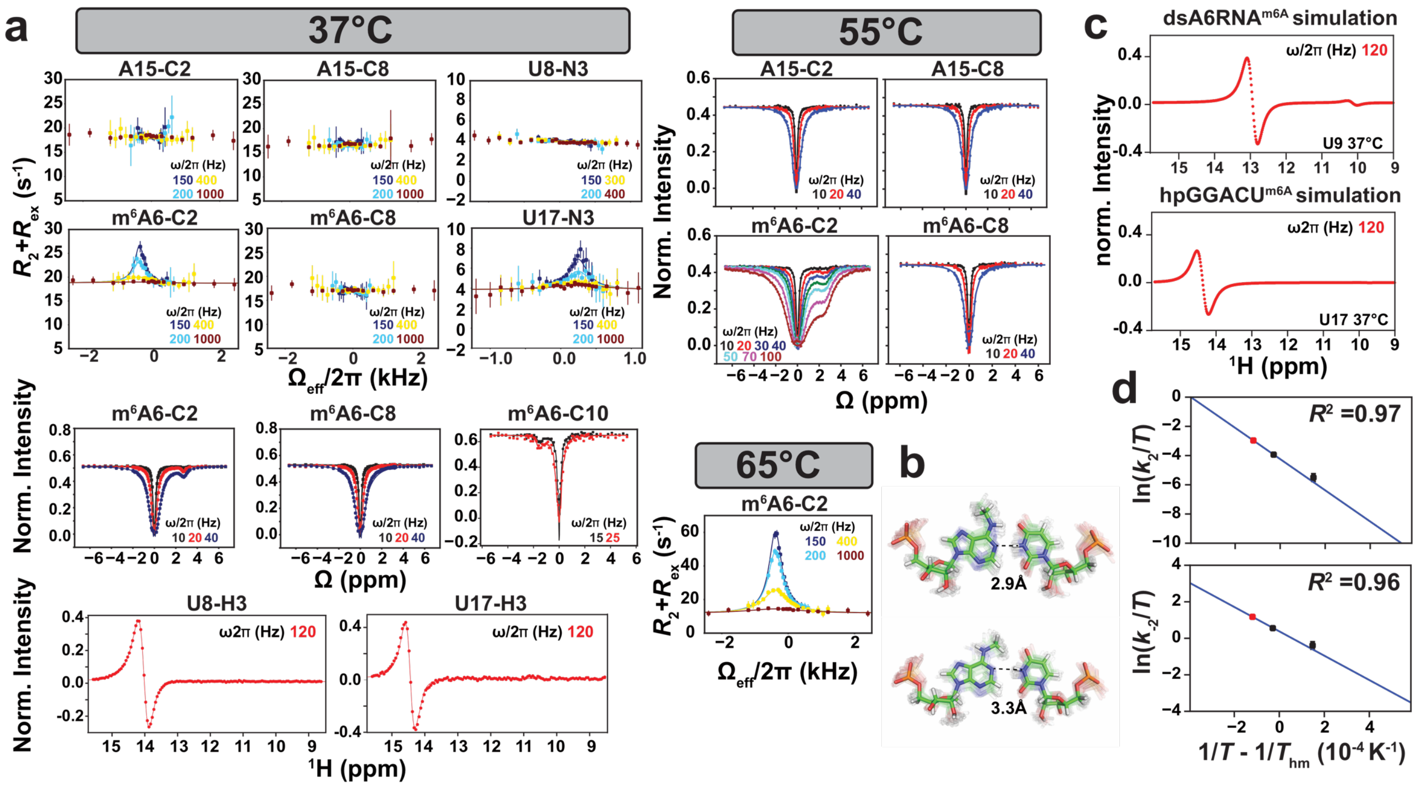


**Extended Data Fig. 6. Characterizing dsRNA$^{syn}$ ES using RD and MD simulations.** **a**, $R_{1\rho}$ and CEST profiles measured for hpGGACU$^{m6A}$ at T = 37°C-65°C. Spin-lock powers for $R_{1\rho}$ and RF field powers for CEST profiles are color coded. Solid lines denote fits or simulations performed using the Bloch-McConnell equations as described in Methods. Error bars for CEST profiles (smaller than data points) were obtained using standard deviation of 3 measurements of peak intensity with zero relaxation delay as described in Methods. Error bars for $R_{1\rho}$ profiles were obtained using Monte-Carlo simulations as described in Methods. **b**, Model structures for m$^6$(*anti*)A-U and m$^6$(*syn*)A···U base pairs generated using MD simulations, highlighting the average imino hydrogen bond lengths in both conformations. **c**, $^1$H CEST simulations showing the absence of the minor peak in hpGGACU$^{m6A}$ likely due to the small ES2 population (~0.6%) but a detectable minor peak in dsA6RNA$^{m6A}$ which has a higher population (~1.2%). **d**, van't Hoff plots showing the temperature dependence of the forward ($k_2$) and reverse ($k_{-2}$) rate constants for methylamino isomerization in hpGGACU$^{m6A}$. Data points in black and red were measured using C10 CEST and C2 $R_{1\rho}$ respectively. $R^2$ denotes coefficient of determination (Methods). Error bars were obtained by propagating the errors in $k_2$ and $k_{-2}$ obtained from $R_{1\rho}$ or CEST measurements. Buffer used for RD measurements was composed of 15 mM sodium phosphate, 25 mM sodium chloride, 0.1 mM EDTA in a 90% $H_2O$:10% $D_2O$ mixture at pH 6.8.

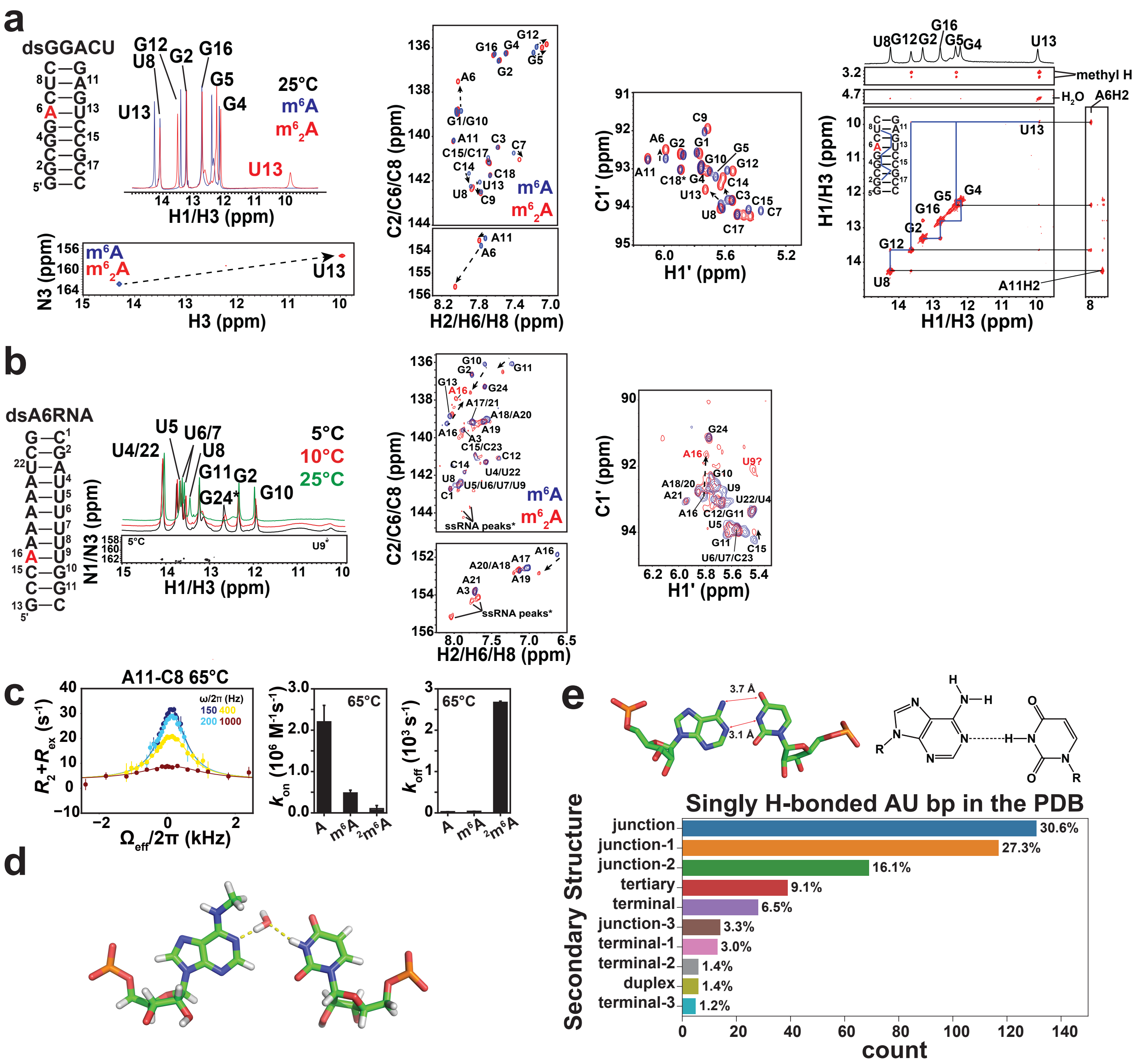

**Extended Data Fig. 7. Conformational characterization of m$^6$(*syn*)A···U bp using the m$^6_2$A···U ES mimic and a structure-based survey.** (**a**, **b**) dsGGACU and dsA6RNA constructs showing methylation sites (in red) along with 2D NMR spectra. **a**, 1D $^1$H (T = 25°C), 2D [$^{15}$N, $^1$H] HSQC (T = 25°C), 2D [$^{13}$C, $^1$H] HSQC (T = 37°C) spectra for m$^6$A (blue) and m$^6_2$A (red) modified dsGGACU along with 2D NOESY spectra (T = 25°C) showing sequential connectivity between imino protons for m$^6_2$A dsGGACU. U13 N3 is $^{15}$N site-labeled in both modified RNAs. **b**, 1D $^1$H (T = 5°C,10°C and 25°C) spectra for m$^6_2$A A6RNA along with 2D [$^{15}$N, $^1$H] HSQC (T = 5°C) and 2D [$^{13}$C, $^1$H] HSQC (T = 10°C) spectra for m$^6$A (blue) and m$^6_2$A (red) modified dsA6RNA. **c**, $^{13}$C $R_{1\rho}$ profile for A11-C8 in dsGGACU$^{m62A}$ at T = 65°C, and comparison of apparent $k_{on}$ and $k_{off}$ in unmethylated, m$^6$A modified and

$m^6{}_2A$ modified dsGGACU at T = 65°C.  Solid lines denote a fit of the data to the Bloch-McConnell equations (Methods).  Spin-lock powers for $R_{1\rho}$ RD profiles are color coded.  Error bars for the $R_{1\rho}$ profiles and for the rate constants were obtained using a Monte-Carlo based approach as described in Methods.  **d**, A snapshot of the $m^6$(*syn*)A···U bp in dsGGACU$^{m6A}$ from MD simulations where the methylamino group of $m^6A$ was enforced into a *syn* conformation, showing a water mediated hydrogen bond.  **e,** Survey of the PDB[3] reveals 428 singly H-bonded A-U bps with structures similar to the $m^6$(*syn*)A ···U bp.  Shown are the different motifs (for definition see Methods) in which the singly H-bonded A-U bps were identified.  Also shown is a representative example (PDBID: 1LNG) of a singly H-bonded A-U bp from the PDB along with its chemical structure.  Buffer used for NMR experiments was composed of 15 mM sodium phosphate, 25 mM sodium chloride, 0.1 mM EDTA in a 90% $H_2O$:10% $D_2O$ mixture at pH 6.8.

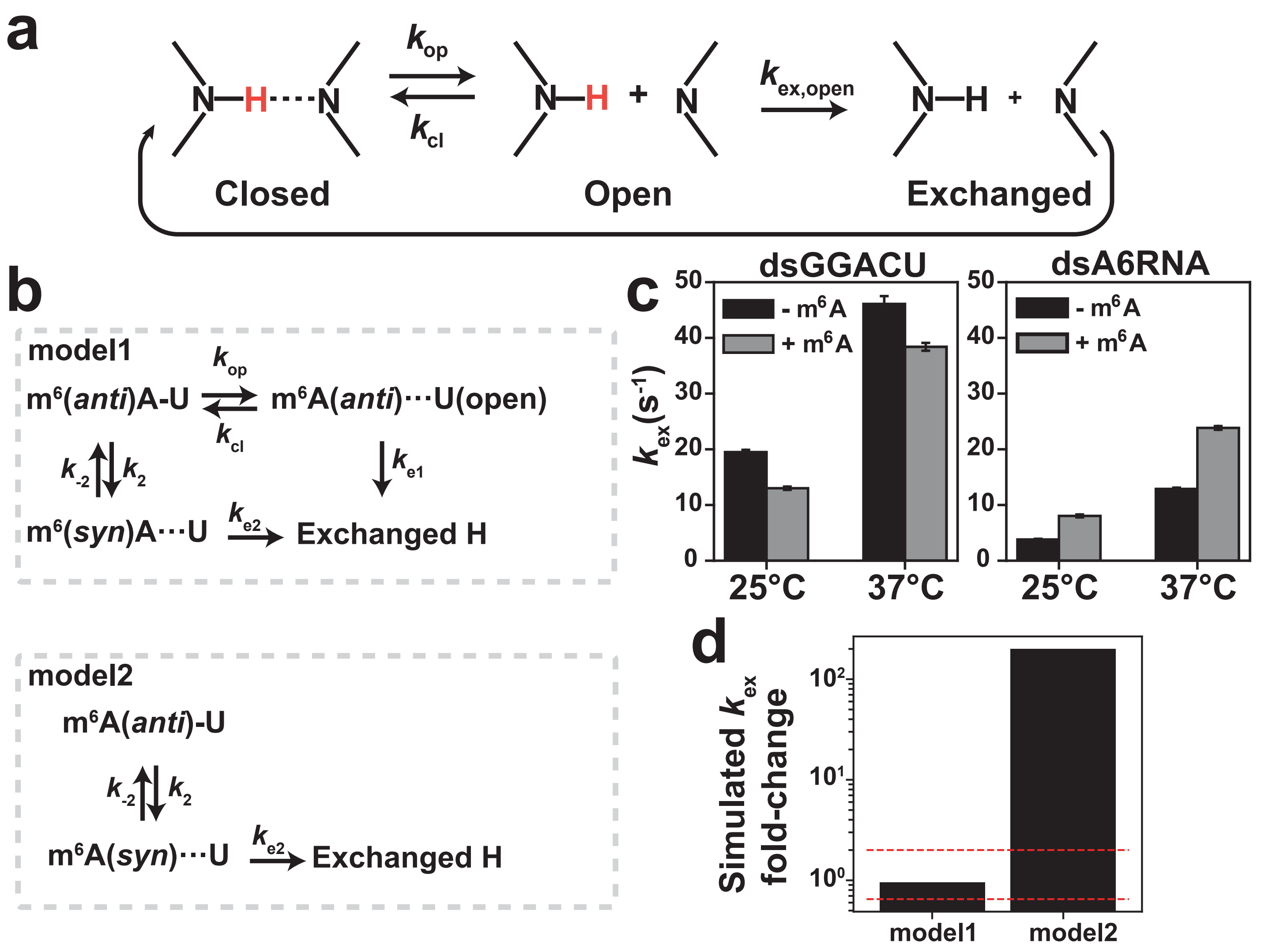


**Extended Data Fig. 8. Imino proton exchange measurements.** **a**, Schematic of a two-step imino proton exchange for an RNA base pair. **b**, Models for m$^6$A-U base opening. In model 1, m$^6$(*syn*)A···U bp can also contribute to solvent exchange in addition to the canonical base opening state. In model 2 m$^6$(*syn*)A···U replaces the canonical base opening state and is the dominant contributior to solvent exchange (Supplementary Note 2). **c**, The apparent exchange rate constant of imino and water proton for unmethylated ($k_{ex}$) and methylated ($k_{ex,m6A}^{app}$) dsGGACU and dsA6RNA measured using imino proton exchange experiments. Error bars in $k_{ex}$ were the standard fitting errors as described in Methods. **d**, Comparison of simulated fold-change ($k_{ex}/k_{ex,m6A}^{app}$) of $k_{ex}$ in unmethylated and methylated RNA based on model 1 or 2. The two red dashed lines indicate the range of fold-change measured experimentally in panel **c**. Buffer used for imino exchange experiments was composed of 15 mM sodium phosphate, 25 mM sodium chloride, 0.1 mM EDTA in a 90% $H_2O$:10% $D_2O$ mixture at pH 6.8.

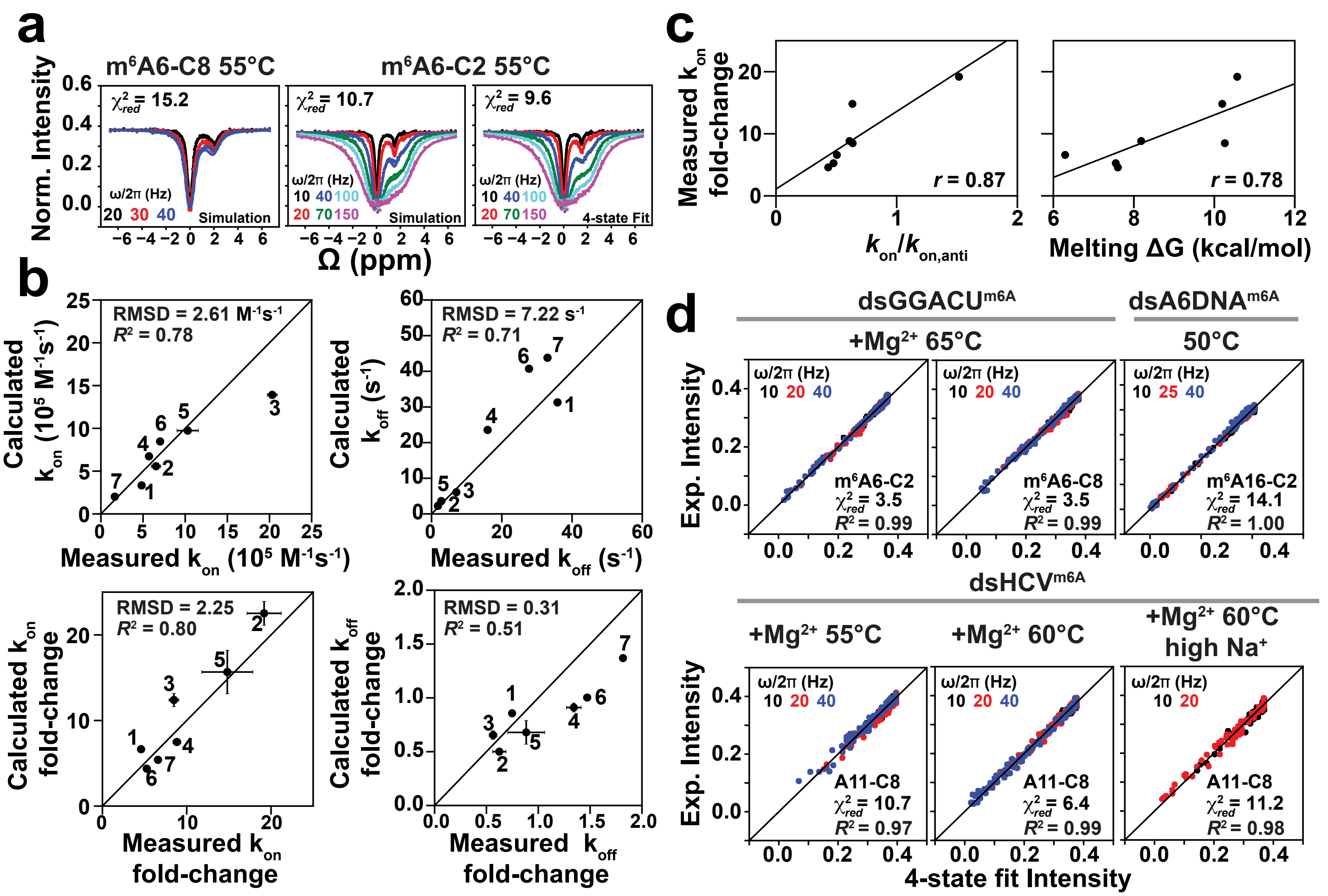


**Extended Data Fig. 9. Predicting the impact of m⁶A on hybridization kinetics in dsRNA/DNA. a**, Comparison of experimental m⁶A6 C2/C8 CEST data points measured for dsGGACU$^{m6A}$ at T = 55°C with a line computed from 4-state simulations (Methods) of the CS+IF model without any adjustable parameters (Simulation, left and middle) or from constrained 4-state CS+IF fit (4-state Fit, right). **b**, Comparison of the experimentally measured and predicted (using the CS+IF 4-state CEST simulation) apparent $k_{on}$ , $k_{off}$ and the fold-change relative to unmethylated duplex ($k_{on}$ fold-change = $k_{on}$(unmethylated)/ $k_{on,m6A}^{app}$ and $k_{off}$ fold-change = $k_{off}$(unmethylated)/ $k_{off,m6A}^{app}$ ) for RNA and DNA duplexes. Each point corresponds to a different duplex and/or experimental condition. All buffers contained 40 mM $Na^+$, unless stated otherwise: (1) dsGGACU$^{m6A}$ at T = 65° C, (2) at T = 55° C, (3) with 3 mM $Mg^{2+}$ at T = 65°C; (4) dsHCV$^{m6A}$ with 3 mM $Mg^{2+}$ at T = 60° C, (5) with 3 mM $Mg^{2+}$ at T = 55° C, (6) with 3 mM $Mg^{2+}$ and 100 mM $Na^+$ at T = 60° C; (7) dsA6DNA$^{m6A}$ at T = 50° C. **c**, Correlations between experimentally measured apparent $k_{on}$ fold-change and $k_{on}$(unmethylated)/ $k_{on,anti}$ (left) or melting free energy of the unmethylated dsRNA (right). $r$ is the Pearson's correlation coefficient. **d**, Comparison of measured CEST intensities[2] for m⁶A6 C2/C8 in dsGGACU$^{m6A}$, A11 C8 in dsHCV$^{m6A}$ and m⁶A16 C2 in dsA6DNA$^{m6A}$ at different conditions (+$Mg^{2+}$

corresponds to with 3 mM $Mg^{2+}$ and high $Na^+$ corresponds to with 100 mM $Na^+$) and values obtained from constrained 4-state fit using the CS+IF model. $R^2$ denotes coefficient of determination. Error bars for CEST profiles (smaller than data points) were obtained from standard deviations. Error bars for $k_{on}$ and $k_{off}$ were determined using a Monte-Carlo approach as described in Methods.

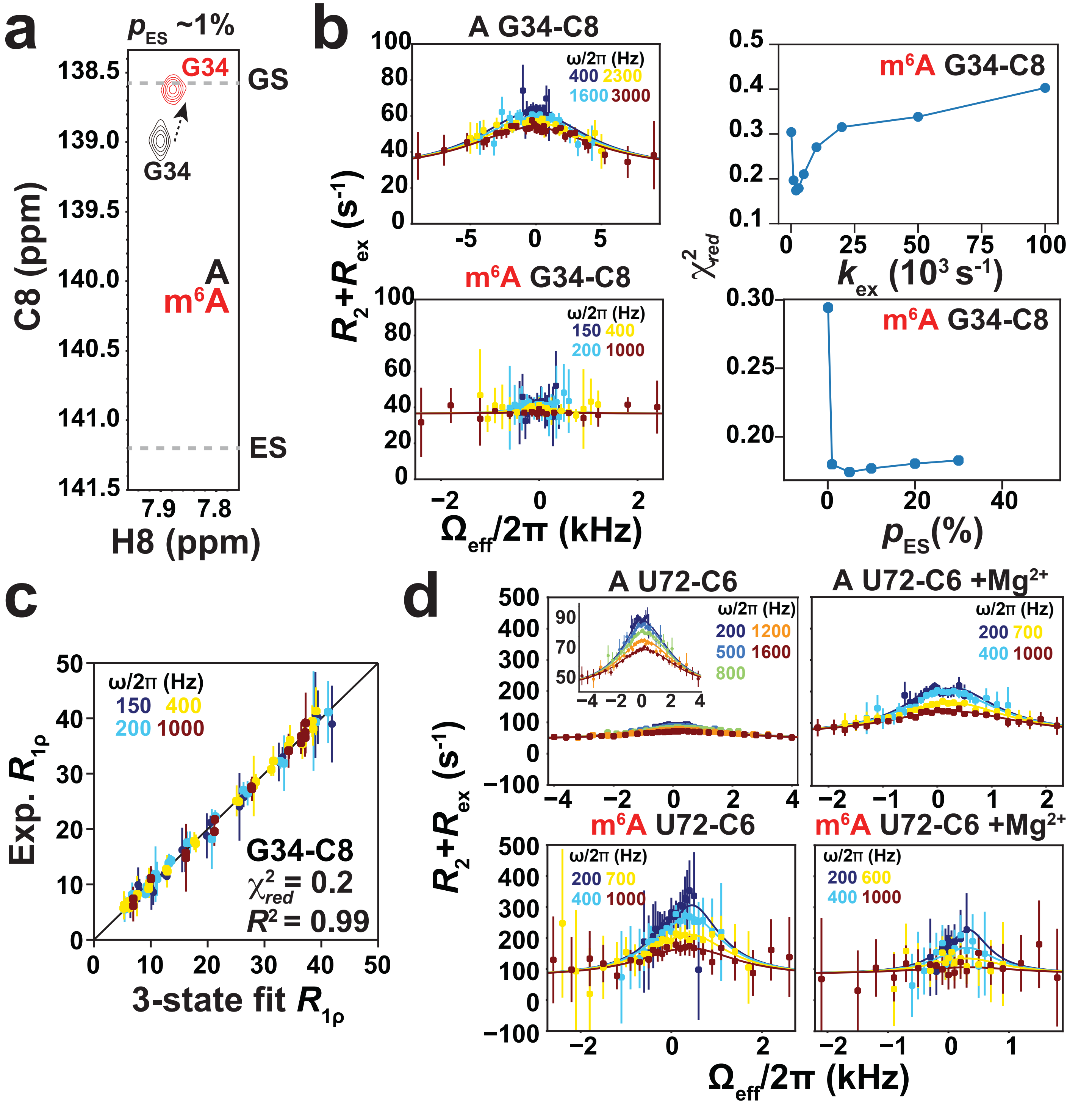


**Extended Data Fig. 10. m⁶A slows select RNA conformational transitions.** **a,** 2D [¹³C, ¹H] HSQC spectra of unmethylated and methylated TAR showing the G34-C8 resonance at T = 25°C. Also shown are the ¹³C chemical shifts for G34 C8 in unmethylated and methylated TAR, and GS and ES peak positions of unmethylated/methylated RNA (indicated with gray dashed lines)[4]. **b,** Comparison of G34 C8 ¹³C $R_{1\rho}$ profiles measured for unmethylated TAR in a prior study[4] with m⁶A35 methylated TAR at T = 25°C. Also shown is $\chi^2_{red}$ from 2-state fit of the RD data measured for methylated TAR G34-C8 as a function of varying $k_{ex}$ (top) or $p_{ES}$ (bottom). **c,** Comparison of measured and constrained 3-state (CS model) fit $R_{1\rho}$ value. A 3-state CS fit was performed because CS flux is >99% based on CS+IF modeling (Methods). Errors in $R_{1\rho}$ values were estimated using Monte-Carlo simulations as described in Methods. **d,** $^{13}C$ $R_{1\rho}$ RD profiles of unmethylated and methylated RREIIB for U72 C6 with and without 3 mM $Mg^{2+}$ at T = 25°C. Solid lines in the $R_{1\rho}$ RD profiles denote fits of the data to the Bloch-McConnnell equations (Methods). Spin-lock powers for $R_{1\rho}$ RD profiles are color coded. Error bars for the $R_{1\rho}$ profiles were obtained using Monte-Carlo simulations as described in Methods. Buffer used for RD measurements was composed of 15 mM sodium phosphate, 25 mM sodium chloride, 0.1 mM EDTA in a 90% $H_2O$:10% $D_2O$ mixture at pH 6.8.

### Supplementary Tables

**Supplementary Table 1. Exchange parameters obtained from unconstrained 2-state fitting of $^{13}$C and $^{15}$N $R_{1\rho}$, and $^{13}$C and $^{15}$N CEST data.** Shown are the chemical shift difference between GS and ES (Δω), ES population ($p_{ES}$), exchange rate constant ($k_{ex}$), longitudinal relaxation rate constant ($R_1$), transverse relaxation rate constant for the GS ($R_2$) and ES ($R_{2,ES}$ is assumed to be equal to $R_2$ except for CEST experiments measuring duplex hybridization[2]) and the $\chi^2_{red}$ obtained from fitting $R_{1\rho}$ and CEST data. Errors for all exchange parameters were determined using a Monte-Carlo approach as described in Methods.

| $R_{1\rho}$ | | | | | | | | |
|---|---|---|---|---|---|---|---|---|
| Sample | Resonance | Δω (ppm) | $p_{ES}$ (%) | $k_{ex}$ ($s^{-1}$) | $R_1$ ($s^{-1}$) | $R_2$ ($s^{-1}$) | $R_{2,ES}$ ($s^{-1}$) | $\chi^2_{red}$ |
| ssGGACU$^{m6A}$ 20°C | m$^6$A6-C2 | -0.53±0.02 | 9.2±0.5 | 506±14 | 4.59±0.07 | 9.76±0.08 | - | 0.3 |
| ssGGACU$^{m6A}$ 25°C | m$^6$A6-C2 | -0.55±0.05 | 8.4±1.3 | 607±38 | 4.53±0.19 | 8.26±0.22 | - | 2.1 |
| ssGGACU$^{m6A}$ 37°C | m$^6$A6-C2 | -0.52±0.08 | 11.8±4.1 | 2333±95 | 4.00±0.08 | 5.41±0.15 | - | 0.3 |
| ssA6RNA$^{m6A}$ 20°C | m$^6$A6-C2 | -0.72±0.08 | 4.3±1.0 | 901±137 | 4.65±0.12 | 10.32±0.15 | - | 0.2 |
| m$^6$AMP 20°C | C2 | -0.76±0.01 | 6.6±0.2 | 1400±20 | 2.62±0.03 | 3.14±0.04 | - | 0.5 |
| m$^6$AMP 25°C | C2 | -0.83±0.05 | 5.8±0.7 | 2241±62 | 2.21±0.05 | 3.00±0.10 | - | 0.9 |
| m$^6$AMP 37°C | C2 | -0.89±0.22 | 4.6±2.3 | 6051±266 | 1.67±0.03 | 2.25±0.15 | - | 1.2 |
| m$^6$AMP $Mg^{2+}$, 25°C | C2 | -0.63±0.03 | 9.4±0.8 | 2357±46 | 2.62±0.03 | 3.46±0.07 | - | 0.8 |
| hpGGACU$^{m6A}$ 37°C[a] | m$^6$A6-C2 | 2.76±0.08 | 0.6 | 176±5 | 3.44±0.02 | 18.74±0.03 | - | 0.3 |
| | U17-N3 | -4.31±0.14 | | | 2.17±0.02 | 3.90±0.03 | - | |
| hpGGACU$^{m6A}$ 65°C | m$^6$A6-C2 | 2.60±0.02 | 1.6±0.0 | 1113±20 | 4.84±0.04 | 11.87±0.06 | - | 0.4 |
| dsGGACU$^{m6A}$ 37°C | m$^6$A6-C2 | 2.86±0.10 | 0.8±0.6 | 118±95 | 4.03±0.03 | 15.03±0.03 | - | 0.4 |
| | U13-N3 | -4.74±0.22 | | | 2.14±0.04 | 3.30±0.04 | - | |
| dsGGACU$^{m62A}$ 65°C | A11-C8 | -1.09±0.05 | 9.2±0.9 | 2913±35 | 2.36±0.05 | 3.69±1.04 | 0.00±10.57 | 0.5 |
| dsA6RNA$^{m6A}$ 37°C | m$^6$A16-C2 | 1.84±0.08 | 1.2±0.5 | 103±60 | 4.54±0.03 | 21.02±0.03 | - | 0.4 |
| TAR 25°C[b] | G34-C8 | 2.64±0.48 | 21.9±10.3 | 32251±4213 | 3.43±0.26 | 28.94±5.64 | - | 0.4 |
| TAR$^{m6A}$ 25°C[c] | G34-C8 | 1.31±0.10 | 1 | 2397±468 | 1.71±0.13 | 36.50±0.31 | - | 0.2 |
| RRE 25°C (ES1)[d] | U72-C6 | -3.41±0.34 | 4.8±1.2 | 10727±573 | 0.85±0.13 | 47.41±1.36 | - | 0.5 |
| RRE 25°C (ES2)[d] | U72-C6 | -0.53±0.08 | 36.6±4.5 | 2529±1557 | | | - | |
| RRE$^{m6A}$ 25°C | U72-C6 | -3.30±0.16 | 18.9±1.5 | 6105±253 | 2.52±1.47 | 49.96±5.15 | - | 0.1 |
| RRE $Mg^{2+}$, 25°C (ES1)[d] | U72-C6 | -2.30±0.10 | 17.5±2.1 | 6195±423 | 0.00±0.68 | 60.4±10.25 | - | 0.3 |
| RRE $Mg^{2+}$, 25°C (ES2)[d] | U72-C6 | -1.20±0.50 | 3.9±0.1 | 2364±53 | | | | |
| RRE$^{m6A}$ $Mg^{2+}$, 25°C (ES1) | U72-C6 | -2.2±0.07 | 20.7±1.7 | 6004±212 | 0.00±0.56 | 61.85±3.05 | - | 0.2 |
| **CEST** | | | | | | | | |

| | | | | | | | | |
|---|---|---|---|---|---|---|---|---|
| ssGGACU$^{m6A}$ 25°C | m$^6$A6-C10 | 2.96±0.01 | 9.2±0.1 | 640.1±14.1 | 0.94±0.01 | 4.92±1.38 | - | 1.2 |
| ssGGACU$^{m6A}$ 37°C | m$^6$A6-C10 | 2.95±0.02 | 9.0±0.0 | 1957.1±60.0 | 0.71±0.02 | 20.0±2.34 | - | 6.0 |
| ssGGACU$^{m6A}$ 45°C | m$^6$A6-C10 | 3.46±0.10 | 7.0±0.0 | 4413.5±238.0 | 1.08±0.05 | 10.85±2.98 | - | 11.0 |
| dsGGACU 55°C | A6-C8 | 2.55±0.01 | 0.8±0.0 | 136.5±8.4 | 4.61±0.00 | 11.04±0.06 | 128.90±11.17 | 7.5 |
| | A6-C2 | 1.23±0.00 | | | 4.80±0.00 | 11.29±0.06 | 97.70±12.25 | |
| dsGGACU$^{m6A}$ 55°C | m$^6$A6-C8 | 2.08±0.00 | 5.4±0.3 | 33.6±2.1 | 4.55±0.01 | 11.66±0.04 | 154.61±4.21 | 7.4 |
| dsHCV 55°C | A11-C8 | 2.18±0.02 | 1.5±0.7 | 166.5±22.8 | 4.27±0.01 | 13.13±0.03 | 162.44±29.77 | 13.0 |
| dsHCV$^{m6A}$ 55°C | A11-C8 | 2.28±0.00 | 6.0±0.4 | 46.0±4.0 | 4.31±0.03 | 11.96±0.08 | 87.22±4.36 | 10.7 |
| hpGGACU$^{m6A}$ 37°C | m$^6$A6-C2 | 2.78±0.00 | 0.6±0.0 | 196.2±9.6 | 3.30±0.00 | 18.36±0.05 | - | 21.0 |
| | m$^6$A6-C10 | -1.58±0.03 | | | 1.43±0.00 | 4.08±0.07 | - | |
| | U17-N3 | -4.83±0.01 | | | 3.41±0.00 | 9.45±0.07 | - | |
| hpGGACU$^{m6A}$ 55°C | m$^6$A6-C2 | 2.61±0.00 | 1.1±0.0 | 546.8±3.2 | 4.17±0.00 | 12.89±0.04 | - | 21.1 |
| | m$^6$A6-C10 | -1.67±0.01 | | | 2.02±0.01 | 4.99±0.09 | - | |
| dsGGACU$^{m6A}$ $Mg^{2+}$,37°C | m$^6$A6-C2 | 2.68±0.02 | 0.4±0.1 | 98.1±23.7 | 3.30±0.01 | 22.78±0.08 | - | 2.4 |
| dsA6DNA$^{m6A}$ 50°C | m$^6$A16-C2 | 1.17±0.00 | 32.4±0.1 | 102.2±0.8 | 2.50±0.01 | 10.79±0.18 | 10.57±0.38 | 14.9 |

[(a)] $p_{ES}$ is ill-defined due to slow exchange. To fit the $R_{1\rho}$ data, $p_{ES}$ was assumed to be equal to the more reliable value obtained from CEST measurement. (b) data obtained from a prior study[4]. (c) Δω and $p_{ES}$ are not well determined. To fit the $R_{1\rho}$ data, $p_{ES}$ was assumed to be equal to the more reliable value obtained from chemical shift analysis (Extended Data Fig. 10a, Methods). (d) data obtained from a prior study[6].

**Supplementary Table 2. Exchange parameters from constrained 3-state CS (Fig. 2a) fits at T = 65°C and constrained 4-state CS+IF (Fig. 5a) fits at T = 55°C and 65°C for dsGGACU$^{m6A}$ m$^6$A6 C2 and C8 $^{13}$C CEST data.**

| Parameters | 65°C (CS) | | 65°C (CS+IF) | | 55°C (CS+IF) | |
|---|---|---|---|---|---|---|
| | m$^6$A6-C2 | m$^6$A6-C8 | m$^6$A6-C2 | m$^6$A6-C8 | m$^6$A6-C2 | m$^6$A6-C8 |
| $\Delta\omega_{dsRNA,syn}$ (ppm) | - | | 2.20±0.00 | 0* | 2.42±0.0 | 0* |
| $\Delta\omega_{ssRNA,syn}$ (ppm) | 1.70±0.00 | 2.45±0.01 | 1.69±0.00 | 2.4±0.0 | 1.56±0.0 | 2.16±0.0 |
| $\Delta\omega_{ssRNA,anti}$ (ppm) | 1.10±0.07 | 0* | 1.08±0.00 | 0* | 0.70±0.0 | 0* |
| $p_{dsRNA,syn}$ (%) | - | | 1.0±0.0 | | 1.2±0.1 | |
| $p_{ssRNA,syn}$ (%) | 22.9±0.1 | | 22.1±0.1 | | 4.5±0.1 | |
| $p_{ssRNA,anti}$ (%) | 2.2±0.1 | | 2.1±0.1 | | 0.2±0.0 | |
| $k_1$ (s$^{-1}$) | 2129.1±126.1 | | 1888.1±29.8 | | 272.5±1.3 | |
| $k_{-1}$ (s$^{-1}$) | 19476.8±131.4 | | 20111.9±186.8 | | 6127.5±1.3 | |
| $k_{on,anti}$ (M$^{-1}$s$^{-1}$) | 5.1±0.4×10$^6$ | | 4.9±0.2 ×10$^6$ | | 13.4±0.0×10$^6$ | |
| $k_{off,anti}$ (s$^{-1}$) | 37.7±3.4 | | 29.4±0.4 | | 1.2±0.0 | |
| $k_2$ (s$^{-1}$) | - | | 14.4±0.0 | | 6.5±0.0 | |
| $k_{-2}$ (s$^{-1}$) | - | | 1075.6±0.0 | | 512.2±1.2 | |
| $k_{on,syn}$ (M$^{-1}$s$^{-1}$) | - | | 1.0±0.0×10$^5$ | | 5.7±0.0×10$^5$ | |
| $k_{off,syn}$ (s$^{-1}$) | - | | 487.9±0.1 | | 90.9±0.2 | |
| $R_{1,GS}$ (s$^{-1}$) | 3.45±0.01 | 3.47±0.00 | 3.44±0.00 | 3.47±0.01 | 4.69±0.00 | 4.48±0.00 |
| $R_{2,GS}$ (s$^{-1}$) | 7.60±0.30 | 4.01±0.54 | 6.93±0.00 | 5.20±0.00 | 10.66±0.02 | 12.70±0.00 |
| $R_{2,ssRNA,syn}$ (s$^{-1}$) | 15.26±1.91 | 60.90±11.96 | 17.10±0.00 | 87.90±3.00 | 35.1±0.33 | 150.2±0.00 |
| $R_{2,ssRNA,anti}$ (s$^{-1}$) | 13.16±1.52 | 63.83±4.22 | 17.10±1.29 | 82.01±3.71 | 35.1±0.94 | 150.2±0.00 |
| $\chi^2_{red}$ | 3.0 | | 3.0 | | 9.1 | |

GS is dsRNA$^{anti}$ in both 3-state and 4-state fits. The two ESs are ssRNA$^{syn}$ and ssRNA$^{anti}$ in 3-state fits (Fig. 2a) while the three ESs are dsRNA$^{syn}$, ssRNA$^{syn}$ and ssRNA$^{anti}$ in 4-state fits (Fig. 5a). $k_1$ and $k_{-1}$ are the forward and backward rate constants, respectively for methylamino isomerization in ssRNA. $k_{on,anti}$ and $k_{off,anti}$ are the annealing and melting rate constants, respectively for hybridization when m$^6$A is *anti*. $k_2$ and $k_{-2}$ are the forward and backward rate constants, respectively for methylamino isomerization in dsRNA. $k_{on,syn}$ and $k_{off,syn}$ are the annealing and melting rate constants, respectively for hybridization when the methylamino is *syn*. Errors for all exchange parameters were determined using a Monte-Carlo approach as described in Methods. * these values were fixed to be 0.

**Supplementary Table 3. Exchange parameters from unconstrained fitting dsGGACU$^{m6A}$ m$^6$A6 C2 CEST data measured at T = 55°C assuming the triangular 3-state model (Fig. 3b).**

| | m$^6$A6-C2 |
|---|---|
| $\Delta\omega_{ss}$ (ppm) | 1.50±0.02 |
| $\Delta\omega_{ES}$ (ppm) | 2.39±0.02 |
| $p_{ss}$ (%) | 4.6±1.1 |
| $p_{ES}$ (%) | 1.4±0.1 |
| $k_{ex,ss}$ (s$^{-1}$) | 9.5±7.9 |
| $k_{ex,2}$ (s$^{-1}$) | 511.5±16.6 |
| $k_{ex,ss\text{-}ES}$ (s$^{-1}$) | 155.0±39.7 |
| $R_1$ (s$^{-1}$) | 4.71±0.06 |
| $R_{2,GS}$ (s$^{-1}$) | 10.63±0.16 |
| $R_{2,ss}$ (s$^{-1}$) | 36.51±8.52 |
| $\chi^2_{red}$ | 9.6 |

In the 3-state fit, GS is dsRNA$^{anti}$, the two minor states are ss, which is the ssRNA (ssRNA$^{syn}$⇌ssRNA$^{anti}$) state, and ES, which is dsRNA$^{syn}$. $k_{ex,ss}$ is the exchange rate constant between GS and ssRNA, $k_{ex,2}$ is the exchange rate constant between GS and ES. $k_{ex,ss\text{-}ES}$ is the exchange rate constant between ss and ES. Errors for all exchange parameters were determined using a Monte-Carlo approach as described in Methods.

**Supplementary Table 4. Thermodynamic parameters from UV melting experiments.** Shown are number of replicates (n), oligo concentration ($C_t$), melting temperature ($T_m$), standard enthalpy difference (ΔH°), entropy difference (ΔS°) and free energy difference ($ΔG°_{37°C}$) at T = 37°C.

| Construct | n | $C_t$ (μM) | $T_m$(°C) | ΔH° (kcal/mol) | ΔS° (e.u.) | $ΔG°_{37°C}$ (kcal/mol) |
|---|---|---|---|---|---|---|
| dsA6RNA | 3 | 3 | 40.7±0.2 | -95.0±1.5 | -275.6±5.0 | -9.4±0.1 |
| $dsA6RNA^{m6A}$ | 3 | 3 | 37.0±0.2 | -93.6±2.6 | -275.3±7.9 | -8.3±0.2 |
| dsA6DNA[(a)] | 3 | 3 | 39.2±0.2 | -98.6±2.9 | -288.8±9.1 | -9.1±0.1 |
| $dsA6DNA^{m6A}$ | 3 | 3 | 36.2±0.1 | -93.6±1.2 | -276.0±3.8 | -8.0±0.1 |

Errors represent one standard deviation (n=3 independent measurements). Uncertainty in the calculated thermodynamic parameters were determined by error propagation as described in Methods.
[(a)] Data obtained from[2].

**Supplementary Table 5. Spin lock powers and offsets used in the $R_{1\rho}$ experiments.**

| Nuclei | [spin lock power] {offset frequencies} |
|---|---|
| | $[\omega/2\pi\ (s^{-1})]\ \{\Omega_{eff}/2\pi\ (s^{-1})\}$ |
| **ssGGACU$^{m6A}$, 20°C** | |
| m$^6$A6-C2 | [150, 200, 250, 300, 400, 500, 600, 700, 900, 1000, 1200, 1400, 1600, 2000, 2500] {0}<br>[50] {-180, -140, -100, -60, -20, 20, 30, 40, 50, 60, 70, 80, 90, 100, 120, 140, 180}<br>[200] {-640, -440, -340, -280, -220, -180, -140, -100, -60, -20, 20, 30, 40, 50, 60, 70, 80, 90, 100, 120, 140, 180, 220, 260, 300, 340, 400, 460, 560, 760}<br>[400] {-1540, -1340, -1140, -990, -840, -690, -540, -390, -240, -190, -140, -90, -40, 10, 60, 110, 160, 210, 260, 310, 360, 510, 660, 810, 960, 1110, 1260, 1460}<br>[1000] {-2340, -1740, -1340, -940, -740, -540, -340, -240, -90, -15, 60, 135, 210, 360, 460, 660, 860, 1060, 1460, 1860, 2460} |
| **ssGGACU$^{m6A}$, 25°C** | |
| m$^6$A6-C2 | [150, 200, 250, 300, 400, 500, 600, 700, 900, 1000, 1200, 1400, 1600, 2000, 2500] {0}<br>[50] {-180, -140, -100, -60, -20, 20, 30, 40, 50, 60, 60, 70, 80, 90, 100, 120, 140, 180}<br>[200] {-640, -440, -340, -280, -220, -180, -140, -100, -60, -20, 20, 30, 40, 50, 60, 60, 70, 80, 90, 100, 120, 140, 180, 220, 260, 300, 340, 400, 460, 560}<br>[400] {-1540, -1340, -1140, -990, -840, -690, -540, -390, -240, -190, -140, -90, -40, 10, 60, 60, 110, 160, 210, 260, 310, 360, 510, 660, 810, 960, 1110, 1260, 1460}<br>[1000] {-2340, -1740, -1340, -940, -740, -540, -340, -240, -90, -15, 60, 60, 135, 210, 360, 460, 660, 860, 1060, 1460, 1860, 2460} |
| **ssGGACU$^{m6A}$, 37°C** | |
| m$^6$A6-C2 | [150, 200, 250, 300, 400, 500, 600, 700, 900, 1000, 1200, 1400, 1600, 2000, 2500] {0}<br>[50] {-180, -140, -100, -60, -20, 20, 30, 40, 50, 60, 60, 70, 80, 90, 100, 120, 140, 180}<br>[200] {-640, -440, -340, -280, -220, -180, -140, -100, -60, -20, 20, 30, 40, 50, 60, 60, 70, 80, 90, 100, 120, 140, 180, 220, 260, 300, 340, 400, 460, 560}<br>[400] {-1540, -1340, -1140, -990, -840, -690, -540, -390, -240, -190, -140, -90, -40, 10, 60, 60, 110, 160, 210, 260, 310, 360, 510, 660, 810, 960, 1110, 1260, 1460}<br>[1000] {-2340, -1740, -1340, -940, -740, -540, -340, -240, -90, -15, 60, 60, 135, 210, 360, 460, 660, 860, 1060, 1460, 1860, 2460} |
| **ssA6$^{m6A}$, 20°C** | |
| m$^6$A6-C2 | [150, 200, 250, 300, 400, 500, 600, 700, 900, 1000, 1200, 1400, 1600, 2000, 2500] {0}<br>[150] {-400, -340, -280, -240, -200, -160, -120, -80, -40, 40, 80, 120, 160, 200, 240, 280, 340, 400}<br>[200] {-600, -500, -400, -350, -300, -250, -200, -150, -100, -50, 50, 100, 150, 200, 250, 300, 350, 400, 500, 600}<br>[400] {-1200, -1050, -900, -750, -600, -450, -300, -250, -200, -150, -100, -50, 50, 100, 150, 200, 250, 300, 450, 600, 750, 900, 1050, 1200}<br>[1000] {-2400, -1800, -1200, -900, -600, -300, -150, -50, 50, 150, 300, 600, 900, 1200, 1800, 2400} |
| **m$^6$AMP, 20°C** | |
| C2 | [150, 200, 250, 300, 400, 500, 600, 700, 900, 1000, 1200, 1400, 1600, 2000, 2500] {0}<br>[150] {-400, -340, -280, -240, -200, -160, -120, -80, -40, 40, 80, 120, 160, 200, 240, 280, 340, 400}<br>[200] {-600, -500, -400, -350, -300, -250, -200, -150, -100, -50, 50, 100, 150, 200, 250, 300, 350, 400, 500, 600}<br>[400] {-1200, -1050, -900, -750, -600, -450, -300, -250, -200, -150, -100, -50, 50, 100, 150, 200, 250, 300, 450, 600, 750, 900, 1050, 1200}<br>[1000] {-2400, -1800, -1200, -900, -600, -300, -150, -50, 50, 150, 300, 600, 900, 1200, 1800, 2400} |
| **AMP, 25°C** | |
| C2 | [150, 200, 250, 300, 400, 500, 600, 700, 900, 1000, 1200, 1400, 1600, 2000, 2500] {0}<br>[150] {-400, -340, -280, -240, -200, -160, -120, -80, -40, 40, 80, 120, 160, 200, 240, 280, 340, 400}<br>[200] {-600, -500, -400, -350, -300, -250, -200, -150, -100, -50, 50, 100, 150, 200, 250, 300, 350, 400, 500, 600}<br>[400] {-1200, -1050, -900, -750, -600, -450, -300, -250, -200, -150, -100, -50, 50, 100, 150, 200, 250, 300, 450, 600, 750, 900, 1050, 1200}<br>[1000] {-2400, -1800, -1200, -900, -600, -300, -150, 50, 150, 300, 600, 900, 1200, 1800, 2400} |
| C8 | [150, 200, 250, 300, 400, 500, 600, 700, 900, 1000, 1200, 1400, 1600, 2000, 2500] {0}<br>[150] {-400, -340, -280, -240, -200, -160, -120, -80, -40, 40, 80, 120, 160, 200, 240, 280, 340, 400} |

| | |
|---|---|
| | [200] {-600, -500, -400, -350, -300, -250, -200, -150, -100, -50, 50, 100, 150, 200, 250, 300, 350, 400, 500, 600}<br>[400] {-1200, -1050, -900, -750, -600, -450, -300, -250, -200, -150, -100, -50, 50, 100, 150, 200, 250, 300, 450, 600, 750, 900, 1050, 1200}<br>[1000] {-2400, -1800, -1200, -900, -600, -300, -150, 50, 150, 300, 600, 900, 1200, 1800, 2400} |
| **m$^6$AMP, 25°C** | |
| C2 | [150, 200, 250, 300, 400, 500, 600, 700, 900, 1000, 1200, 1400, 1600, 2000, 2500] {0}<br>[150] {-400, -340, -280, -240, -200, -160, -120, -80, -40, 40, 80, 120, 160, 200, 240, 280, 340, 400}<br>[200] {-600, -500, -400, -350, -300, -250, -200, -150, -100, -50, 50, 100, 150, 200, 250, 300, 350, 400, 500, 600}<br>[400] {-1200, -1050, -900, -750, -600, -450, -300, -250, -200, -150, -100, -50, 50, 100, 150, 200, 250, 300, 450, 600, 750, 900, 1050, 1200}<br>[1000] {-2400, -1800, -1200, -900, -600, -300, -150, 50, 150, 300, 600, 900, 1200, 1800, 2400} |
| C8 | [150, 200, 250, 300, 400, 500, 600, 700, 900, 1000, 1200, 1400, 1600, 2000, 2500] {0}<br>[150] {-400, -340, -280, -240, -200, -160, -120, -80, -40, 40, 80, 120, 160, 200, 240, 280, 340, 400}<br>[200] {-600, -500, -400, -350, -300, -250, -200, -150, -100, -50, 50, 100, 150, 200, 250, 300, 350, 400, 500, 600}<br>[400] {-1200, -1050, -900, -750, -600, -450, -300, -250, -200, -150, -100, -50, 50, 100, 150, 200, 250, 300, 450, 600, 750, 900, 1050, 1200}<br>[1000] {-2400, -1800, -1200, -900, -600, -300, -150, 50, 150, 300, 600, 900, 1200, 1800, 2400} |
| **m$^6$AMP, 37°C** | |
| C2 | [150, 200, 250, 300, 400, 500, 600, 700, 900, 1000, 1200, 1400, 1600, 2000, 2500] {0}<br>[150] {-400, -340, -280, -240, -200, -160, -120, -80, -40, 40, 80, 120, 160, 200, 240, 280, 340, 400}<br>[200] {-600, -500, -400, -350, -300, -250, -200, -150, -100, -50, 50, 100, 150, 200, 250, 300, 350, 400, 500, 600}<br>[400] {-1200, -1050, -900, -750, -600, -450, -300, -250, -200, -150, -100, -50, 50, 100, 150, 200, 250, 300, 450, 600, 750, 900, 1050, 1200}<br>[1000] {-2400, -1800, -1200, -900, -600, -300, -150, -50, 50, 150, 300, 600, 900, 1200, 1800, 2400} |
| **AMP 3mM $Mg^{2+}$, 25°C** | |
| C2 | [150, 200, 250, 300, 400, 500, 600, 700, 900, 1000, 1200, 1400, 1600, 2000, 2500] {0}<br>[150] {-400, -340, -280, -240, -200, -160, -120, -80, -40, 40, 80, 120, 160, 200, 280, 340, 400}<br>[200] {-600, -500, -400, -350, -300, -250, -200, -100, -50, 50, 100, 150, 200, 250, 300, 350, 400, 500, 600}<br>[400] {-1200, -1050, -900, -750, -600, -450, -300, -250, -200, -150, -100, -50, 50, 100, 150, 200, 250, 300, 450, 600, 750, 900, 1050, 1200}<br>[1000] {-2400, -1800, -1200, -900, -600, -300, -150, -50, 50, 150, 300, 600, 900, 1200, 1800, 2400} |
| C8 | [150, 200, 250, 300, 400, 500, 600, 700, 900, 1000, 1200, 1400, 1600, 2000, 2500] {0}<br>[150] {-400, -340, -280, -240, -200, -160, -120, -80, -40, 40, 80, 120, 160, 200, 280, 340, 400}<br>[200] {-600, -500, -400, -350, -300, -250, -200, -100, -50, 50, 100, 150, 200, 250, 300, 350, 400, 500, 600}<br>[400] {-1200, -1050, -900, -750, -600, -450, -300, -250, -200, -150, -100, -50, 50, 100, 150, 200, 250, 300, 450, 600, 750, 900, 1050, 1200}<br>[1000] {-2400, -1800, -1200, -900, -600, -300, -150, -50, 50, 150, 300, 600, 900, 1200, 1800, 2400} |
| **m$^6$AMP 3mM $Mg^{2+}$, 25°C** | |
| C2 | [150, 200, 250, 300, 400, 500, 600, 700, 900, 1000, 1200, 1400, 1600, 2000, 2500] {0}<br>[150] {-400, -340, -280, -240, -200, -160, -120, -80, -40, 40, 80, 120, 160, 200, 280, 340, 400}<br>[200] {-600, -500, -400, -350, -300, -250, -200, -100, -50, 50, 100, 150, 200, 250, 300, 350, 400, 500, 600}<br>[400] {-1200, -1050, -900, -750, -600, -450, -300, -250, -200, -150, -100, -50, 50, 100, 150, 200, 250, 300, 450, 600, 750, 900, 1050, 1200}<br>[1000] {-2400, -1800, -1200, -900, -600, -300, -150, -50, 50, 150, 300, 600, 900, 1200, 1800, 2400} |
| C8 | [150, 200, 250, 300, 400, 500, 600, 700, 900, 1000, 1200, 1400, 1600, 2000, 2500] {0}<br>[150] {-400, -340, -280, -240, -200, -160, -120, -80, -40, 40, 80, 120, 160, 200, 280, 340, 400}<br>[200] {-600, -500, -400, -350, -300, -250, -200, -100, -50, 50, 100, 150, 200, 250, 300, 350, 400, 500, 600}<br>[400] {-1200, -1050, -900, -750, -600, -450, -300, -250, -200, -150, -100, -50, 50, 100, 150, 200, 250, 300, 450, 600, 750, 900, 1050, 1200}<br>[1000] {-2400, -1800, -1200, -900, -600, -300, -150, -50, 50, 150, 300, 600, 900, 1200, 1800, 2400} |
| **hpGGACU$^{m6A}$, 37°C** | |

| | |
|---|---|
| m$^{6}$A6-C2 | [150, 200, 250, 300, 400, 500, 600, 700, 900, 1000, 1200, 1400, 1600, 2000, 2500] {0}<br>[150] {-400, -340, -280, -240, -200, -160, -120, -80, -40, 40, 80, 120, 160, 200, 240, 280, 340, 400}<br>[200] {-600, -500, -400, -350, -300, -250, -200, -150, -100, -50, 50, 100, 150, 200, 250, 300, 350, 400, 500, 600}<br>[400] {-1200, -1050, -900, -750, -600, -450, -300, -250, -200, -150, -100, -50, 50, 100, 150, 200, 250, 300, 450, 600, 750, 900, 1050, 1200}<br>[1000] {-2400, -1800, -1200, -900, -600, -300, -150, -50, 50, 150, 300, 600, 900, 1200, 1800, 2400} |
| **hpGGACU$^{m6A}$, 55°C** | |
| m$^{6}$A6-C2 | [150, 200, 250, 300, 400, 500, 600, 700, 900, 1000, 1200, 1400, 1600, 2000, 2500] {0}<br>[150] {-400, -340, -280, -240, -200, -160, -120, -80, -40, 40, 80, 120, 160, 200, 240, 280, 340, 400}<br>[200] {-600, -500, -400, -350, -300, -250, -200, -150, -100, -50, 50, 100, 150, 200, 250, 300, 350, 400, 500, 600}<br>[400] {-1200, -1050, -900, -750, -600, -450, -300, -250, -200, -150, -100, -50, 50, 100, 150, 200, 250, 300, 450, 600, 750, 900, 1050, 1200}<br>[1000] {-2400, -1800, -1200, -900, -600, -300, -150, -50, 50, 150, 300, 600, 900, 1200, 1800, 2400} |
| m$^{6}$A6-C8 | [150, 200, 250, 300, 400, 500, 600, 700, 900, 1000, 1200, 1400, 1600, 2000, 2500] {0}<br>[150] {-400, -340, -280, -240, -200, -160, -120, -80, -40, 40, 80, 120, 160, 200, 240, 280, 340, 400}<br>[200] {-600, -500, -400, -350, -300, -250, -200, -150, -100, -50, 50, 100, 150, 200, 250, 300, 350, 400, 500, 600}<br>[400] {-1200, -1050, -900, -750, -600, -450, -300, -250, -200, -150, -100, -50, 50, 100, 150, 200, 250, 300, 450, 600, 750, 900, 1050, 1200}<br>[1000] {-2400, -1800, -1200, -900, -600, -300, -150, -50, 50, 150, 300, 600, 900, 1200, 1800, 2400} |
| **hpGGACU$^{m6A}$, 65°C** | |
| m$^{6}$A6-C2 | [150, 200, 250, 300, 400, 500, 600, 700, 900, 1000, 1200, 1400, 1600, 2000, 2500] {0}<br>[150] {-400, -340, -280, -240, -200, -160, -120, -80, -40, 40, 80, 120, 160, 200, 240, 280, 340, 400}<br>[200] {-600, -500, -400, -350, -300, -250, -200, -150, -100, -50, 50, 100, 150, 200, 250, 300, 350, 400, 500, 600}<br>[400] {-1200, -1050, -900, -750, -600, -450, -300, -250, -200, -150, -100, -50, 50, 100, 150, 200, 250, 300, 450, 600, 750, 900, 1050, 1200}<br>[1000] {-2400, -1800, -1200, -900, -600, -300, -150, -50, 50, 150, 300, 600, 900, 1200, 1800, 2400} |
| **dsGGACU, 37°C** | |
| A6-C2 | [150, 200, 250, 300, 400, 500, 600, 700, 900, 1000, 1200, 1400, 1600, 2000, 2500] {0}<br>[150] {-400, -340, -280, -240, -200, -160, -120, -80, -40, 40, 80, 120, 160, 200, 240, 280, 340, 400}<br>[200] {-600, -500, -400, -350, -300, -250, -200, -150, -100, -50, 50, 100, 150, 200, 250, 300, 350, 400, 500, 600}<br>[400] {-1200, -1050, -900, -750, -600, -450, -300, -250, -200, -150, -100, -50, 50, 100, 150, 200, 250, 300, 450, 600, 750, 900, 1050, 1200}<br>[1000] {-2400, -1800, -1200, -900, -600, -300, -150, -50, 50, 150, 300, 600, 900, 1200, 1800, 2400} |
| A6-C8 | [150, 200, 250, 300, 400, 500, 600, 700, 900, 1000, 1200, 1400, 1600, 2000, 2500] {0}<br>[150] {-400, -340, -280, -240, -200, -160, -120, -80, -40, 40, 80, 120, 160, 200, 240, 280, 340, 400}<br>[200] {-600, -500, -400, -350, -300, -250, -200, -150, -100, -50, 50, 100, 150, 200, 250, 300, 350, 400, 500, 600}<br>[400] {-1200, -1050, -900, -750, -600, -450, -300, -250, -200, -150, -100, -50, 50, 100, 150, 200, 250, 300, 450, 600, 750, 900, 1050, 1200}<br>[1000] {-2400, -1800, -1200, -900, -600, -300, -150, -50, 50, 150, 300, 600, 900, 1200, 1800, 2400} |
| U13-N3 | [150, 200, 250, 300, 400, 500, 600, 700, 900, 1000, 1200, 1400, 1600, 2000, 2500] {0}<br>[150] {-300, -200, -140, -80, -40, 40, 80, 120, 160, 200, 200, 240, 280, 320, 360, 400, 440}<br>[200] {-600, -400, -300, -200, -150, -100, -50, 50, 100, 150, 200, 200, 250, 300, 350, 400, 450, 500, 550, 600}<br>[300] {-850, -700, -550, -400, -300, -250, -200, -100, -50, 50, 100, 150, 200, 200, 250, 300, 350, 400, 450, 500, 600, 650, 700, 800}<br>[400] {-1200, -1000, -850, -700, -550, -400, -250, -100, -50, 50, 100, 150, 200, 200, 250, 300, 350, 400, 450, 500, 650, 800, 950, 1100} |
| **dsGGACU$^{m6A}$, 37°C** | |
| m$^{6}$A6-C2 | [150, 200, 250, 300, 400, 500, 600, 700, 900, 1000, 1200, 1400, 1600, 2000, 2500] {0}<br>[150] {-400, -340, -280, -240, -200, -160, -120, -80, -40, 40, 80, 120, 160, 200, 240, 280, 340, 400}<br>[200] {-600, -500, -400, -350, -300, -250, -200, -150, -100, -50, 50, 100, 150, 200, 250, 300, 350, 400, 500, 600} |

| | |
|---|---|
| | [400] {-1200, -1050, -900, -750, -600, -450, -300, -250, -200, -150, -100, -50, 50, 100, 150, 200, 250, 300, 450, 600, 750, 900, 1050, 1200}<br>[1000] {-2400, -1800, -1200, -900, -600, -300, -150, -50, 50, 150, 300, 600, 900, 1200, 1800, 2400} |
| $m^6$A6-C8 | [150, 200, 250, 300, 400, 500, 600, 700, 900, 1000, 1200, 1400, 1600, 2000, 2500] {0}<br>[150] {-400, -340, -280, -240, -200, -160, -120, -80, -40, 40, 80, 120, 160, 200, 240, 280, 340, 400}<br>[200] {-600, -500, -400, -350, -300, -250, -200, -150, -100, -50, 50, 100, 150, 200, 250, 300, 350, 400, 500, 600}<br>[400] {-1200, -1050, -900, -750, -600, -450, -300, -250, -200, -150, -100, -50, 50, 100, 150, 200, 250, 300, 450, 600, 750, 900, 1050, 1200}<br>[1000] {-2400, -1800, -1200, -900, -600, -300, -150, -50, 50, 150, 300, 600, 900, 1200, 1800, 2400} |
| U13-N3 | [150, 200, 250, 300, 400, 500, 600, 700, 900, 1000, 1200, 1400, 1600, 2000, 2500] {0}<br>[150] {-300, -200, -140, -80, -40, 40, 80, 120, 160, 200, 200, 240, 280, 320, 360, 400, 440}<br>[200] {-600, -400, -300, -200, -150, -100, -50, 50, 100, 150, 200, 200, 250, 300, 350, 400, 450, 500, 550, 600}<br>[300] {-850, -700, -550, -400, -300, -250, -200, -100, -50, 50, 100, 150, 200, 200, 250, 300, 350, 400, 450, 500, 600, 650, 700, 800}<br>[400] {-1200, -1000, -850, -700, -550, -400, -250, -100, -50, 50, 100, 150, 200, 200, 250, 300, 350, 400, 450, 500, 650, 800, 950, 1100} |
| **dsGGACU$^{m62A}$, 65°C** | |
| A11-C8 | [150, 200, 250, 300, 400, 500, 600, 700, 900, 1000, 1200, 1400, 1600, 2000, 2500] {0}<br>[150] {-400, -340, -280, -240, -200, -160, -120, -80, -40, 40, 80, 120, 160, 200, 240, 280, 340, 400}<br>[200] {-600, -500, -400, -350, -300, -250, -200, -150, -100, -50, 50, 100, 150, 200, 250, 300, 350, 400, 500, 600}<br>[400] {-1200, -1050, -900, -750, -600, -450, -300, -250, -200, -150, -100, -50, 50, 100, 150, 200, 250, 300, 450, 600, 750, 900, 1050, 1200}<br>[1000] {-2400, -1800, -1200, -900, -600, -300, -150, -50, 50, 150, 300, 600, 900, 1200, 1800, 2400} |
| **dsA6RNA, 37°C** | |
| A16-C2 | [150, 200, 250, 300, 400, 500, 600, 700, 900, 1000, 1200, 1400, 1600, 2000, 2500] {0}<br>[150] {-400, -340, -280, -240, -200, -160, -120, -80, -40, 40, 80, 120, 160, 200, 240, 280, 340, 400}<br>[200] {-600, -500, -400, -350, -300, -250, -200, -150, -100, -50, 50, 100, 150, 200, 250, 300, 350, 400, 500, 600}<br>[400] {-1200, -1050, -900, -750, -600, -450, -300, -250, -200, -150, -100, -50, 50, 100, 150, 200, 250, 300, 450, 600, 750, 900, 1050, 1200}<br>[1000] {-2400, -1800, -1200, -900, -600, -300, -150, -50, 50, 150, 300, 600, 900, 1200, 1800, 2400} |
| A16-C8 | [150, 200, 250, 300, 400, 500, 600, 700, 900, 1000, 1200, 1400, 1600, 2000, 2500] {0}<br>[150] {-400, -340, -280, -240, -200, -160, -120, -80, -40, 40, 80, 120, 160, 200, 240, 280, 340, 400}<br>[200] {-600, -500, -400, -350, -300, -250, -200, -150, -100, -50, 50, 100, 150, 200, 250, 300, 350, 400, 500, 600}<br>[400] {-1200, -1050, -900, -750, -600, -450, -300, -250, -200, -150, -100, -50, 50, 100, 150, 200, 250, 300, 450, 600, 750, 900, 1050, 1200}<br>[1000] {-2400, -1800, -1200, -900, -600, -300, -150, -50, 50, 150, 300, 600, 900, 1200, 1800, 2400} |
| U9-N3 | [150, 200, 250, 300, 400, 500, 600, 700, 900, 1000, 1200, 1400, 1600, 2000, 2500] {0}<br>[150] {-300, -200, -140, -80, -40, 40, 80, 120, 160, 200, 200, 240, 280, 320, 360, 400, 440}<br>[200] {-600, -400, -300, -200, -150, -100, -50, 50, 100, 150, 200, 200, 250, 300, 350, 400, 450, 500, 550, 600}<br>[300] {-850, -700, -550, -400, -300, -250, -200, -100, -50, 50, 100, 150, 200, 200, 250, 300, 350, 400, 450, 500, 600, 650, 700, 800}<br>[400] {-1200, -1000, -850, -700, -550, -400, -250, -100, -50, 50, 100, 150, 200, 200, 250, 300, 350, 400, 450, 500, 650, 800, 950, 1100} |
| **dsA6RNA$^{m6A}$, 37°C** | |
| $m^6$A16-C2 | [150, 200, 250, 300, 400, 500, 600, 700, 900, 1000, 1200, 1400, 1600, 2000, 2500] {0}<br>[150] {-400, -340, -280, -240, -200, -160, -120, -80, -40, 40, 80, 120, 160, 200, 240, 280, 340, 400}<br>[200] {-600, -500, -400, -350, -300, -250, -200, -150, -100, -50, 50, 100, 150, 200, 250, 300, 350, 400, 500, 600}<br>[400] {-1200, -1050, -900, -750, -600, -450, -300, -250, -200, -150, -100, -50, 50, 100, 150, 200, 250, 300, 450, 600, 750, 900, 1050, 1200}<br>[1000] {-2400, -1800, -1200, -900, -600, -300, -150, -50, 50, 150, 300, 600, 900, 1200, 1800, 2400} |

| | |
|---|---|
| $m^6$A16-C8 | [150, 200, 250, 300, 400, 500, 600, 700, 900, 1000, 1200, 1400, 1600, 2000, 2500] {0}<br>[150] {-400, -340, -280, -240, -200, -160, -120, -80, -40, 40, 80, 120, 160, 200, 240, 280, 340, 400}<br>[200] {-600, -500, -400, -350, -300, -250, -200, -150, -100, -50, 50, 100, 150, 200, 250, 300, 350, 400, 500, 600}<br>[400] {-1200, -1050, -900, -750, -600, -450, -300, -250, -200, -150, -100, -50, 50, 100, 150, 200, 250, 300, 450, 600, 750, 900, 1050, 1200}<br>[1000] {-2400, -1800, -1200, -900, -600, -300, -150, -50, 50, 150, 300, 600, 900, 1200, 1800, 2400} |
| U9-N3 | [150, 200, 250, 300, 400, 500, 600, 700, 900, 1000, 1200, 1400, 1600, 2000, 2500] {0}<br>[150] {-300, -200, -140, -80, -40, 40, 80, 120, 160, 200, 200, 240, 280, 320, 360, 400, 440}<br>[200] {-600, -400, -300, -200, -150, -100, -50, 50, 100, 150, 200, 200, 250, 300, 350, 400, 450, 500, 550, 600}<br>[300] {-850, -700, -550, -400, -300, -250, -200, -100, -50, 50, 100, 150, 200, 200, 250, 300, 350, 400, 450, 500, 600, 650, 700, 800}<br>[400] {-1200, -1000, -850, -700, -550, -400, -250, -100, -50, 50, 100, 150, 200, 200, 250, 300, 350, 400, 450, 500, 650, 800, 950, 1100} |
| **TAR$^{m6A35}$, 25°C** | |
| G34-C8 | [150, 200, 250, 300, 400, 500, 600, 700, 900, 1000, 1200, 1400, 1600, 2000, 2500] {0}<br>[150] {-400, -340, -280, -240, -200, -160, -120, -80, -40, 40, 80, 120, 160, 200, 240, 280, 340, 400}<br>[200] {-600, -500, -400, -350, -300, -250, -200, -150, -100, -50, 50, 100, 150, 200, 250, 300, 350, 400, 500, 600}<br>[400] {-1200, -1050, -900, -750, -600, -450, -300, -250, -200, -150, -100, -50, 50, 100, 150, 200, 250, 300, 450, 600, 750, 900, 1050, 1200}<br>[1000] {-2400, -1800, -1200, -900, -600, -300, -150, -50, 50, 150, 300, 600, 900, 1200, 1800, 2400} |
| **RRE$^{m6A68}$, 25°C** | |
| U72-C6 | [100, 150, 200, 250, 300, 400, 500, 700, 900, 1200, 1600, 2000, 2400, 2800] {0}<br>[200] {-600, -500, -400, -350, -300, -250, -200, -150, -100, -50, 50, 100, 150, 200, 250, 300, 350, 400, 500, 600}<br>[400] {-1100, -900, -700, -600, -500, -450, -400, -350, -300, -200, -100, 100, 200, 300, 400, 500, 600, 700, 900, 1100}<br>[700] {-2400, -2100, -1800, -1500, -1200, -1000, -800, -600, -500, -400, -300, -200, -100, 100, 200, 300, 400, 500, 700, 900, 1100, 1400, 1700}<br>[1000] {-2600, -2200, -1800, -1500, -1200, -900, -700, -500, -400, -300, -200, -100, 100, 200, 300, 400, 500, 700, 900, 1100, 1400, 1800, 2200, 2600} |
| **RRE$^{m6A68}$ 3mM $Mg^{2+}$, 25°C** | |
| U72-C6 | [150, 200, 250, 300, 400, 500, 600, 800, 1000, 1600, 2000, 2500, 3500] {0}<br>[200] {-300, -200, -100, -50, -25, 25, 50, 100, 200, 300}<br>[400] {-600, -500, -400, -300, -200, -100, -50, 50, 100, 200, 300, 400, 500, 600}<br>[600] {-900, -700, -500, -300, -200, -100, -50, 50, 100, 200, 300, 500, 700, 900}<br>[1000] {-2100, -1500, -1200, -900, -700, -500, -300, -200, -100, 100, 200, 300, 500, 700, 900, 1200, 1500, 1800} |

**Supplementary Table 6. RF field powers and offsets used in CEST experiments.**

| Nuclei | [RF field power] {offset frequencies}<br>[ω/2π ($s^{-1}$)] {Ω/2π ($s^{-1}$)} |
|---|---|
| **ssGGACU$^{m6A}$, 25°C** | |
| m$^6$A6-C10 | [30] {-801, -751, -701, -651, -601, -585, -570, -554, -538, -522, -507, -491, -475, -459, -443, -428, -412, -396, -380, -364, -349, -333, -317, -301, -281, -260, -239, -219, -198, -177, -156, -136, -115, -94, -74, -53, -32, -12, 8, 29, 49, 70, 91, 112, 132, 153, 174, 194, 215, 236, 256, 277, 298, 314, 329, 345, 361, 377, 392, 408, 424, 440, 456, 471, 487, 503, 519, 535, 550, 566, 582, 598, 648, 698, 748, 798}<br>[50] {-803, -753, -703, -653, -603, -587, -571, -556, -540, -524, -508, -492, -477, -461, -445, -429, -413, -398, -382, -366, -350, -334, -319, -303, -282, -262, -241, -220, -199, -179, -158, -137, -117, -96, -75, -55, -34, -13, 6, 27, 48, 69, 89, 110, 131, 151, 172, 193, 213, 234, 255, 275, 296, 312, 328, 343, 359, 375, 391, 407, 422, 438, 454, 470, 486, 501, 517, 533, 549, 565, 580, 596, 646, 696, 746, 796} |
| **ssGGACU$^{m6A}$, 37°C** | |
| m$^6$A6-C10 | [50] {-796, -746, -696, -646, -596, -580, -565, -549, -533, -517, -501, -486, -470, -454, -438, -422, -407, -391, -375, -359, -344, -328, -312, -296, -275, -255, -234, -213, -193, -172, -151, -131, -110, -89, -69, -48, -27, -6, 13, 34, 55, 75, 96, 117, 137, 158, 179, 199, 220, 241, 261, 282, 303, 319, 334, 350, 366, 382, 398, 413, 429, 445, 461, 477, 492, 508, 524, 540, 555, 571, 587, 603, 653, 703, 753, 803}<br>[80] {-803, -753, -703, -653, -603, -587, -571, -556, -540, -524, -508, -492, -477, -461, -445, -429, -414, -398, -382, -366, -350, -335, -319, -303, -282, -262, -241, -220, -200, -179, -158, -137, -117, -96, -75, -55, -34, -13, 6, 27, 48, 68, 89, 110, 130, 151, 172, 193, 213, 234, 255, 275, 296, 312, 328, 343, 359, 375, 391, 407, 422, 438, 454, 470, 485, 501, 517, 533, 549, 564, 580, 596, 646, 696, 746, 796} |
| **ssGGACU$^{m6A}$, 45°C** | |
| m$^6$A6-C10 | [150] {-805, -785, -765, -745, -724, -704, -684, -664, -643, -623, -603, -583, -562, -542, -522, -502, -481, -461, -441, -421, -400, -380, -360, -340, -319, -299, -279, -259, -238, -218, -198, -178, -157, -137, -117, -96, -76, -56, -36, -15, 4, 24, 44, 65, 85, 105, 125, 146, 166, 186, 206, 227, 247, 267, 287, 308, 328, 348, 368, 389, 409, 429, 449, 470, 490, 510, 530, 551, 571, 591, 611, 632, 652, 672, 692, 713, 733, 753, 773, 794}<br>[300] {-806, -785, -765, -745, -725, -704, -684, -664, -644, -623, -603, -583, -563, -542, -522, -502, -481, -461, -441, -421, -400, -380, -360, -340, -319, -299, -279, -259, -238, -218, -198, -178, -157, -137, -117, -97, -76, -56, -36, -16, 4, 24, 44, 64, 85, 105, 125, 145, 166, 186, 206, 226, 247, 267, 287, 307, 328, 348, 368, 388, 409, 429, 449, 469, 490, 510, 530, 550, 571, 591, 611, 631, 652, 672, 692, 712, 733, 753, 773, 793} |
| **dsGGACU, 55°C** | |
| A6-C2 | [20] {-1000, -944, -888, -833, -777, -722, -666, -611, -555, -500, -489, -479, -469, -459, -449, -439, -429, -419, -409, -398, -388, -378, -368, -358, -348, -338, -328, -318, -308, -297, -287, -277, -267, -257, -247, -237, -227, -217, -207, -196, -186, -176, -166, -156, -146, -136, -126, -116, -106, -95, -85, -75, -65, -55, -45, -35, -25, -15, -5, 5, 15, 25, 35, 45, 55, 65, 75, 85, 95, 106, 116, 126, 136, 146, 156, 166, 176, 186, 196, 207, 217, 227, 237, 247, 257, 267, 277, 287, 297, 308, 318, 328, 338, 348, 358, 368, 378, 388, 398, 409, 419, 429, 439, 449, 459, 469, 479, 489, 500, 555, 611, 666, 722, 777, 833, 888, 944, 1000}<br>[30] {-1000, -944, -888, -833, -777, -722, -666, -611, -555, -500, -489, -479, -469, -459, -449, -439, -429, -419, -409, -398, -388, -378, -368, -358, -348, -338, -328, -318, -308, -297, -287, -277, -267, -257, -247, -237, -227, -217, -207, -196, -186, -176, -166, -156, -146, -136, -126, -116, -106, -95, -85, -75, -65, -55, -45, -35, -25, -15, -5, 5, 15, 25, 35, 45, 55, 65, 75, 85, 95, 106, 116, 126, 136, 146, 156, 166, 176, 186, 196, 207, 217, 227, 237, 247, 257, 267, 277, 287, 297, 308, 318, 328, 338, 348, 358, 368, 378, 388, 398, 409, 419, 429, 439, 449, 459, 469, 479, 489, 500, 555, 611, 666, 722, 777, 833, 888, 944, 1000}<br>[40] {-1000, -944, -888, -833, -777, -722, -666, -611, -555, -500, -489, -479, -469, -459, -449, -439, -429, -419, -409, -398, -388, -378, -368, -358, -348, -338, -328, -318, -308, -297, -287, -277, -267, -257, -247, -237, -227, -217, -207, -196, -186, -176, -166, -156, -146, -136, -126, -116, -106, -95, -85, -75, -65, -55, -45, -35, -25, -15, -5, 5, 15, 25, 35, 45, 55, 65, 75, 85, 95, 106, 116, 126, 136, 146, 156, 166, 176, 186, 196, 207, 217, 227, 237, 247, 257, 267, 277, 287, 297, 308, 318, 328, 338, 348, 358, 368, 378, 388, 398, 409, 419, 429, 439, 449, 459, 469, 479, 489, 500, 555, 611, 666, 722, 777, 833, 888, 944, 1000} |

| | |
|---|---|
| A6-C8 | [20] {-1000.0, -944.4, -888.9, -833.3, -777.8, -722.2, -666.7, -611.1, -555.6, -500.0, -500.0, -487.3, -474.7, -462.0, -449.4, -436.7, -424.1, -411.4, -398.7, -386.1, -373.4, -360.8, -348.1, -335.4, -322.8, -310.1, -297.5, -284.8, -272.2, -259.5, -246.8, -234.2, -221.5, -208.9, -196.2, -183.5, -170.9, -158.2, -145.6, -132.9, -120.3, -107.6, -94.9, -82.3, -69.6, -57.0, -44.3, -31.6, -19.0, -6.3, 6.3, 19.0, 31.6, 44.3, 57.0, 69.6, 82.3, 94.9, 107.6, 120.3, 132.9, 145.6, 158.2, 170.9, 183.5, 196.2, 208.9, 221.5, 234.2, 246.8, 259.5, 272.2, 284.8, 297.5, 310.1, 322.8, 335.4, 348.1, 360.8, 373.4, 386.1, 398.7, 411.4, 424.1, 436.7, 449.4, 462.0, 474.7, 487.3, 500.0, 500.0, 555.6, 611.1, 666.7, 722.2, 777.8, 833.3, 888.9, 944.4, 1000.0}<br>[40] {-1000.0, -944.4, -888.9, -833.3, -777.8, -722.2, -666.7, -611.1, -555.6, -500.0, -500.0, -487.3, -474.7, -462.0, -449.4, -436.7, -424.1, -411.4, -398.7, -386.1, -373.4, -360.8, -348.1, -335.4, -322.8, -310.1, -297.5, -284.8, -272.2, -259.5, -246.8, -234.2, -221.5, -208.9, -196.2, -183.5, -170.9, -158.2, -145.6, -132.9, -120.3, -107.6, -94.9, -82.3, -69.6, -57.0, -44.3, -31.6, -19.0, -6.3, 6.3, 19.0, 31.6, 44.3, 57.0, 69.6, 82.3, 94.9, 107.6, 120.3, 132.9, 145.6, 158.2, 170.9, 183.5, 196.2, 208.9, 221.5, 234.2, 246.8, 259.5, 272.2, 284.8, 297.5, 310.1, 322.8, 335.4, 348.1, 360.8, 373.4, 386.1, 398.7, 411.4, 424.1, 436.7, 449.4, 462.0, 474.7, 487.3, 500.0, 500.0, 555.6, 611.1, 666.7, 722.2, 777.8, 833.3, 888.9, 944.4, 1000.0} |
| **dsGGACU$^{m6A6}$, 55°C** | |
| m$^6$A6-C2 | [10] {-1000, -944, -888, -833, -777, -722, -666, -611, -555, -500, -489, -479, -469, -459, -449, -439, -429, -419, -409, -398, -388, -378, -368, -358, -348, -338, -328, -318, -308, -297, -287, -277, -267, -257, -247, -237, -227, -217, -207, -196, -186, -176, -166, -156, -146, -136, -126, -116, -106, -95, -85, -75, -65, -55, -45, -35, -25, -15, -5, 5, 15, 25, 35, 45, 55, 65, 75, 85, 95, 106, 116, 126, 136, 146, 156, 166, 176, 186, 196, 207, 217, 227, 237, 247, 257, 267, 277, 287, 297, 308, 318, 328, 338, 348, 358, 368, 378, 388, 398, 409, 419, 429, 439, 449, 459, 469, 479, 489, 500, 555, 611, 666, 722, 777, 833, 888, 944, 1000}<br>[20] {-1000, -944, -888, -833, -777, -722, -666, -611, -555, -500, -489, -479, -469, -459, -449, -439, -429, -419, -409, -398, -388, -378, -368, -358, -348, -338, -328, -318, -308, -297, -287, -277, -267, -257, -247, -237, -227, -217, -207, -196, -186, -176, -166, -156, -146, -136, -126, -116, -106, -95, -85, -75, -65, -55, -45, -35, -25, -15, -5, 5, 15, 25, 35, 45, 55, 65, 75, 85, 95, 106, 116, 126, 136, 146, 156, 166, 176, 186, 196, 207, 217, 227, 237, 247, 257, 267, 277, 287, 297, 308, 318, 328, 338, 348, 358, 368, 378, 388, 398, 409, 419, 429, 439, 449, 459, 469, 479, 489, 500, 555, 611, 666, 722, 777, 833, 888, 944, 1000}<br>[40] {-1000, -944, -888, -833, -777, -722, -666, -611, -555, -500, -489, -479, -469, -459, -449, -439, -429, -419, -409, -398, -388, -378, -368, -358, -348, -338, -328, -318, -308, -297, -287, -277, -267, -257, -247, -237, -227, -217, -207, -196, -186, -176, -166, -156, -146, -136, -126, -116, -106, -95, -85, -75, -65, -55, -45, -35, -25, -15, -5, 5, 15, 25, 35, 45, 55, 65, 75, 85, 95, 106, 116, 126, 136, 146, 156, 166, 176, 186, 196, 207, 217, 227, 237, 247, 257, 267, 277, 287, 297, 308, 318, 328, 338, 348, 358, 368, 378, 388, 398, 409, 419, 429, 439, 449, 459, 469, 479, 489, 500, 555, 611, 666, 722, 777, 833, 888, 944, 1000}<br>[70] {-1000, -944, -888, -833, -777, -722, -666, -611, -555, -500, -489, -479, -469, -459, -449, -439, -429, -419, -409, -398, -388, -378, -368, -358, -348, -338, -328, -318, -308, -297, -287, -277, -267, -257, -247, -237, -227, -217, -207, -196, -186, -176, -166, -156, -146, -136, -126, -116, -106, -95, -85, -75, -65, -55, -45, -35, -25, -15, -5, 5, 15, 25, 35, 45, 55, 65, 75, 85, 95, 106, 116, 126, 136, 146, 156, 166, 176, 186, 196, 207, 217, 227, 237, 247, 257, 267, 277, 287, 297, 308, 318, 328, 338, 348, 358, 368, 378, 388, 398, 409, 419, 429, 439, 449, 459, 469, 479, 489, 500, 555, 611, 666, 722, 777, 833, 888, 944, 1000}<br>[100] {-1000, -944, -888, -833, -777, -722, -666, -611, -555, -500, -489, -479, -469, -459, -449, -439, -429, -419, -409, -398, -388, -378, -368, -358, -348, -338, -328, -318, -308, -297, -287, -277, -267, -257, -247, -237, -227, -217, -207, -196, -186, -176, -166, -156, -146, -136, -126, -116, -106, -95, -85, -75, -65, -55, -45, -35, -25, -15, -5, 5, 15, 25, 35, 45, 55, 65, 75, 85, 95, 106, 116, 126, 136, 146, 156, 166, 176, 186, 196, 207, 217, 227, 237, 247, 257, 267, 277, 287, 297, 308, 318, 328, 338, 348, 358, 368, 378, 388, 398, 409, 419, 429, 439, 449, 459, 469, 479, 489, 500, 555, 611, 666, 722, 777, 833, 888, 944, 1000}<br>[150] {-1000, -944, -888, -833, -777, -722, -666, -611, -555, -500, -489, -479, -469, -459, -449, -439, -429, -419, -409, -398, -388, -378, -368, -358, -348, -338, -328, -318, -308, -297, -287, -277, -267, -257, -247, -237, -227, -217, -207, -196, -186, -176, -166, -156, -146, -136, -126, -116, -106, -95, -85, -75, -65, -55, -45, -35, -25, -15, -5, 5, 15, 25, 35, 45, 55, 65, 75, 85, 95, 106, 116, 126, 136, 146, 156, 166, 176, 186, 196, 207, 217, 227, 237, 247, 257, 267, 277, 287, 297, 308, 318, 328, 338, 348, 358, 368, 378, 388, 398, 409, 419, 429, 439, 449, 459, 469, 479, 489, 500, 555, 611, 666, 722, 777, 833, 888, 944, 1000} |
| m$^6$A6-C8 | [20] {-1000, -944, -888, -833, -777, -722, -666, -611, -555, -500, -489, -479, -469, -459, -449, -439, -429, -419, -409, -398, -388, -378, -368, -358, -348, -338, -318, -308, -297, -287, -277, -267, -257, -247, -237, -227, -217, -207, -196, -186, -176, -166, -156, -146, -136, -126, -116, -106, -95, -85, -75, -65, -55, -45, -35, -25, -5, 5, 15, 25, 35, 45, 55, 65, 75, 85, 95, 106, 116, 126, 136, 146, 156, 166, 176, 186, 196, 207, 217, 227, 237, 247, 257, 267, 277, 287, 297, 308, 318, 328, 338, 348, 358, 368, 378, 388, 398, 409, 419, 429, 439, 449, 459, 469, 479, 489, 500, 555, 611, 666, 722, 777, 833, 888, 944, 1000}<br>[30] {-1000, -944, -888, -833, -777, -722, -666, -611, -555, -500, -489, -479, -469, -459, -449, -439, -429, -419, -409, -398, -388, -378, -368, -358, -348, -338, -328, -318, -308, -297, -287, -277, -267, -257, -247, |

| | |
|---|---|
| | -237, -227, -217, -207, -196, -186, -176, -166, -156, -146, -136, -126, -116, -106, -95, -85, -75, -65, -55, -45, -35, -5, 5, 15, 25, 35, 45, 55, 65, 75, 85, 95, 106, 116, 126, 136, 146, 156, 166, 176, 186, 196, 207, 217, 227, 237, 247, 257, 267, 277, 287, 297, 308, 318, 328, 338, 348, 358, 368, 378, 388, 398, 409, 419, 429, 439, 449, 459, 469, 479, 489, 500, 555, 611, 666, 722, 777, 833, 888, 944, 1000}<br>[40] {-1000, -944, -888, -833, -777, -722, -666, -611, -555, -500, -489, -479, -469, -459, -449, -439, -429, -419, -409, -398, -388, -378, -368, -358, -348, -338, -328, -318, -308, -297, -287, -277, -267, -257, -247, -237, -227, -217, -207, -196, -186, -176, -166, -156, -146, -136, -126, -116, -106, -95, -85, -75, -65, -55, -45, -5, 5, 35, 45, 55, 65, 75, 85, 95, 106, 116, 126, 136, 146, 156, 166, 176, 186, 196, 207, 217, 227, 237, 247, 257, 267, 277, 287, 297, 308, 318, 328, 338, 348, 358, 368, 378, 388, 398, 409, 419, 429, 439, 449, 459, 469, 479, 489, 500, 555, 611, 666, 722, 777, 833, 888, 944, 1000} |
| **dsA6RNA$^{m6A}$, 37°C** | |
| U4-H3 | [120] { 367, 372, 377, 382, 386, 391, 396, 401, 406, 410, 415, 420, 425, 429, 434, 439, 444, 448, 453, 458, 463, 468, 472, 477, 482, 487, 491, 496, 501, 506, 511, 515, 520, 525, 530, 534, 539, 544, 549, 554, 558, 563, 568, 573, 577, 582, 587, 592, 597, 601, 606, 611, 616, 620, 625, 630, 635, 639, 644, 649, 654, 659, 663, 668, 673, 678, 682, 687, 692, 697, 702, 706, 711, 716, 721, 725, 730, 735, 740, 745, 749, 754, 759, 764, 768, 773, 778, 783, 787, 792, 797, 802, 807, 811, 816, 821, 826, 830, 835, 840, 845, 850, 854, 859, 864, 869, 873, 878, 883, 888, 893, 897, 902, 907, 912, 916, 921, 926, 931, 936, 940, 945, 950, 955, 959, 964, 969, 974, 978, 983, 988, 993, 998, 1002, 1007, 1012, 1017, 1021, 1026, 1031} |
| U8-H3 | [120] { 367, 372, 377, 382, 386, 391, 396, 401, 406, 410, 415, 420, 425, 429, 434, 439, 444, 448, 453, 458, 463, 468, 472, 477, 482, 487, 491, 496, 501, 506, 511, 515, 520, 525, 530, 534, 539, 544, 549, 554, 558, 563, 568, 573, 577, 582, 587, 592, 597, 601, 606, 611, 616, 620, 625, 630, 635, 639, 644, 649, 654, 659, 663, 668, 673, 678, 682, 687, 692, 697, 702, 706, 711, 716, 721, 725, 730, 735, 740, 745, 749, 754, 759, 764, 768, 773, 778, 783, 787, 792, 797, 802, 807, 811, 816, 821, 826, 830, 835, 840, 845, 850, 854, 859, 864, 869, 873, 878, 883, 888, 893, 897, 902, 907, 912, 916, 921, 926, 931, 936, 940, 945, 950, 955, 959, 964, 969, 974, 978, 983, 988, 993, 998, 1002, 1007, 1012, 1017, 1021, 1026, 1031} |
| U9-H3 | [120] { 367, 372, 377, 382, 386, 391, 396, 401, 406, 410, 415, 420, 425, 429, 434, 439, 444, 448, 453, 458, 463, 468, 472, 477, 482, 487, 491, 496, 501, 506, 511, 515, 520, 525, 530, 534, 539, 544, 549, 554, 558, 563, 568, 573, 577, 582, 587, 592, 597, 601, 606, 611, 616, 620, 625, 630, 635, 639, 644, 649, 654, 659, 663, 668, 673, 678, 682, 687, 692, 697, 702, 706, 711, 716, 721, 725, 730, 735, 740, 745, 749, 754, 759, 764, 768, 773, 778, 783, 787, 792, 797, 802, 807, 811, 816, 821, 826, 830, 835, 840, 845, 850, 854, 859, 864, 869, 873, 878, 883, 888, 893, 897, 902, 907, 912, 916, 921, 926, 931, 936, 940, 945, 950, 955, 959, 964, 969, 974, 978, 983, 988, 993, 998, 1002, 1007, 1012, 1017, 1021, 1026, 1031} |
| **hpGGACU$^{m6A}$, 37°C** | |
| $m^6$A6-C2 | [10] {-1000, -922, -844, -766, -688, -611, -533, -455, -377, -300, -290, -281, -272, -262, -253, -244, -234, -225, -216, -206, -197, -188, -178, -169, -160, -151, -141, -132, -123, -113, -104, -95, -85, -76, -67, -57, -48, -39, -30, 30, 39, 49, 58, 68, 77, 87, 97, 106, 116, 125, 135, 145, 154, 164, 173, 183, 193, 202, 212, 221, 231, 241, 250, 260, 269, 279, 288, 298, 308, 317, 327, 336, 346, 356, 365, 375, 384, 394, 404, 413, 423, 432, 442, 452, 461, 471, 480, 490, 500, 555, 611, 666, 722, 777, 833, 888, 944, 1000}<br>[20] {-1000, -922, -844, -766, -688, -611, -533, -455, -377, -300, -290, -281, -272, -262, -253, -244, -234, -225, -216, -206, -197, -188, -178, -169, -160, -151, -141, -132, -123, -113, -104, -95, -85, -76, -67, -57, -48, -39, -30, 30, 39, 49, 58, 68, 77, 87, 97, 106, 116, 125, 135, 145, 154, 164, 173, 183, 193, 202, 212, 221, 231, 241, 250, 260, 269, 279, 288, 298, 308, 317, 327, 336, 346, 356, 365, 375, 384, 394, 404, 413, 423, 432, 442, 452, 461, 471, 480, 490, 500, 555, 611, 666, 722, 777, 833, 888, 944, 1000}<br>[40] {-1000, -922, -844, -766, -688, -611, -533, -455, -377, -300, -290, -281, -272, -262, -253, -244, -234, -225, -216, -206, -197, -188, -178, -169, -160, -151, -141, -132, -123, -113, -104, -95, -85, -76, -67, -57, -48, -39, -30, 30, 39, 49, 58, 68, 77, 87, 97, 106, 116, 125, 135, 145, 154, 164, 173, 183, 193, 202, 212, 221, 231, 241, 250, 260, 269, 279, 288, 298, 308, 317, 327, 336, 346, 356, 365, 375, 384, 394, 404, 413, 423, 432, 442, 452, 461, 471, 480, 490, 500, 555, 611, 666, 722, 777, 833, 888, 944, 1000} |
| $m^6$A6-C8 | [10] {-1000, -922, -844, -766, -688, -611, -533, -455, -377, -300, -290, -281, -272, -262, -253, -244, -234, -225, -216, -206, -197, -188, -178, -169, -160, -151, -141, -132, -123, -113, -104, -95, -85, -76, -67, -57, -48, -39, -30, 30, 39, 49, 58, 68, 77, 87, 97, 106, 116, 125, 135, 145, 154, 164, 173, 183, 193, 202, 212, 221, 231, 241, 250, 260, 269, 279, 288, 298, 308, 317, 327, 336, 346, 356, 365, 375, 384, 394, 404, 413, 423, 432, 442, 452, 461, 471, 480, 490, 500, 555, 611, 666, 722, 777, 833, 888, 944, 1000}<br>[20] {-1000, -922, -844, -766, -688, -611, -533, -455, -377, -300, -290, -281, -272, -262, -253, -244, -234, -225, -216, -206, -197, -188, -178, -169, -160, -151, -141, -132, -123, -113, -104, -95, -85, -76, -67, -57, -48, -39, -30, 30, 39, 49, 58, 68, 77, 87, 97, 106, 116, 125, 135, 145, 154, 164, 173, |

| | |
|---|---|
| | 183, 193, 202, 212, 221, 231, 241, 250, 260, 269, 279, 288, 298, 308, 317, 327, 336, 346, 356, 365, 375, 384, 394, 404, 413, 423, 432, 442, 452, 461, 471, 480, 490, 500, 555, 611, 666, 722, 777, 833, 888, 944, 1000}<br>[40] {-1000, -922, -844, -766, -688, -611, -533, -455, -377, -300, -290, -281, -272, -262, -253, -244, -234, -225, -216, -206, -197, -188, -178, -169, -160, -151, -141, -132, -123, -113, -104, -95, -85, -76, -67, -57, -48, -39, -30, 30, 39, 49, 58, 68, 77, 87, 97, 106, 116, 125, 135, 145, 154, 164, 173, 183, 193, 202, 212, 221, 231, 241, 250, 260, 269, 279, 288, 298, 308, 317, 327, 336, 346, 356, 365, 375, 384, 394, 404, 413, 423, 432, 442, 452, 461, 471, 480, 490, 500, 555, 611, 666, 722, 777, 833, 888, 944, 1000} |
| $m^6$A6-C10 | [15] {-802, -758, -713, -669, -625, -580, -536, -491, -447, -402, -389, -375, -362, -348, -335, -321, -307, -294, -280, -267, -253, -240, -226, -213, -199, -185, -172, -158, -145, -131, -118, -104, -90, -77, -63, -50, -36, -23, -9, 17, 31, 44, 58, 71, 85, 98, 112, 125, 139, 153, 166, 180, 193, 207, 220, 234, 248, 261, 275, 288, 302, 315, 329, 342, 356, 370, 383, 397, 441, 486, 530, 574, 619, 663, 708, 752, 797}<br>[25] {-802, -758, -713, -669, -624, -580, -536, -491, -447, -402, -389, -375, -361, -348, -334, -321, -307, -294, -280, -267, -253, -239, -226, -212, -199, -185, -172, -158, -145, -131, -117, -104, -90, -77, -63, -50, -36, -23, -9, 17, 31, 44, 58, 71, 85, 99, 112, 126, 139, 153, 166, 180, 193, 207, 221, 234, 248, 261, 275, 288, 302, 315, 329, 343, 356, 370, 383, 397, 441, 486, 530, 575, 619, 663, 708, 752, 797} |
| U17-N3 | [10] {-1000.0, -922.2, -844.4, -766.7, -688.9, -611.1, -533.3, -455.6, -377.8, -300.0, -300.0, -290.7, -281.4, -272.1, -262.8, -253.4, -244.1, -234.8, -225.5, -216.2, -206.9, -197.6, -188.3, -179.0, -169.7, -160.3, -151.0, -141.7, -132.4, -123.1, -113.8, -104.5, -95.2, -85.9, -76.6, -67.2, -57.9, -48.6, -39.3, -30.0, 30.0, 39.6, 49.2, 58.8, 68.4, 78.0, 87.6, 97.1, 106.7, 116.3, 125.9, 135.5, 145.1, 154.7, 164.3, 173.9, 183.5, 193.1, 202.7, 212.2, 221.8, 231.4, 241.0, 250.6, 260.2, 269.8, 279.4, 289.0, 298.6, 308.2, 317.8, 327.3, 336.9, 346.5, 356.1, 365.7, 375.3, 384.9, 394.5, 404.1, 413.7, 423.3, 432.9, 442.4, 452.0, 461.6, 471.2, 480.8, 490.4, 500.0, 500.0, 555.6, 611.1, 666.7, 722.2, 777.8, 833.3, 888.9, 944.4, 1000.0} |
| U17-H3 | [120] { 367, 372, 377, 382, 386, 391, 396, 401, 406, 410, 415, 420, 425, 429, 434, 439, 444, 448, 453, 458, 463, 468, 472, 477, 482, 487, 491, 496, 501, 506, 511, 515, 520, 525, 530, 534, 539, 544, 549, 554, 558, 563, 568, 573, 577, 582, 587, 592, 597, 601, 606, 611, 616, 620, 625, 630, 635, 639, 644, 649, 654, 659, 663, 668, 673, 678, 682, 687, 692, 697, 702, 706, 711, 716, 721, 725, 730, 735, 740, 745, 749, 754, 759, 764, 768, 773, 778, 783, 787, 792, 797, 802, 807, 811, 816, 821, 826, 830, 835, 840, 845, 850, 854, 859, 864, 869, 873, 878, 883, 888, 893, 897, 902, 907, 912, 916, 921, 926, 931, 936, 940, 945, 950, 955, 959, 964, 969, 974, 978, 983, 988, 993, 998, 1002, 1007, 1012, 1017, 1021, 1026, 1031} |
| U8-H3 | [120] { 367, 372, 377, 382, 386, 391, 396, 401, 406, 410, 415, 420, 425, 429, 434, 439, 444, 448, 453, 458, 463, 468, 472, 477, 482, 487, 491, 496, 501, 506, 511, 515, 520, 525, 530, 534, 539, 544, 549, 554, 558, 563, 568, 573, 577, 582, 587, 592, 597, 601, 606, 611, 616, 620, 625, 630, 635, 639, 644, 649, 654, 659, 663, 668, 673, 678, 682, 687, 692, 697, 702, 706, 711, 716, 721, 725, 730, 735, 740, 745, 749, 754, 759, 764, 768, 773, 778, 783, 787, 792, 797, 802, 807, 811, 816, 821, 826, 830, 835, 840, 845, 850, 854, 859, 864, 869, 873, 878, 883, 888, 893, 897, 902, 907, 912, 916, 921, 926, 931, 936, 940, 945, 950, 955, 959, 964, 969, 974, 978, 983, 988, 993, 998, 1002, 1007, 1012, 1017, 1021, 1026, 1031} |
| **hpGGACU$^{m6A}$, 55°C** | |
| A15-C2 | [10] {-1000, -944, -888, -833, -777, -722, -666, -611, -555, -500, -490, -480, -471, -461, -452, -442, -432, -423, -413, -404, -394, -384, -375, -365, -356, -346, -336, -327, -317, -308, -298, -288, -279, -269, -260, -250, -241, -231, -221, -212, -202, -193, -183, -173, -164, -154, -145, -135, -125, -116, -106, -97, -87, -77, -68, -58, -49, -39, -30, 30, 39, 49, 58, 68, 77, 87, 97, 106, 116, 125, 135, 145, 154, 164, 173, 183, 193, 202, 212, 221, 231, 241, 250, 260, 269, 279, 288, 298, 308, 317, 327, 336, 346, 356, 365, 375, 384, 394, 404, 413, 423, 432, 442, 452, 461, 471, 480, 490, 500, 555, 611, 666, 722, 777, 833, 888, 944, 1000}<br>[20] {-1000, -944, -888, -833, -777, -722, -666, -611, -555, -500, -490, -480, -471, -461, -452, -442, -432, -423, -413, -404, -394, -384, -375, -365, -356, -346, -336, -327, -317, -308, -298, -288, -279, -269, -260, -250, -241, -231, -221, -212, -202, -193, -183, -173, -164, -154, -145, -135, -125, -116, -106, -97, -87, -77, -68, -58, -49, -39, -30, 30, 39, 49, 58, 68, 77, 87, 97, 106, 116, 125, 135, 145, 154, 164, 173, 183, 193, 202, 212, 221, 231, 241, 250, 260, 269, 279, 288, 298, 308, 317, 327, 336, 346, 356, 365, 375, 384, 394, 404, 413, 423, 432, 442, 452, 461, 471, 480, 490, 500, 555, 611, 666, 722, 777, 833, 888, 944, 1000}<br>[40] {-1000, -944, -888, -833, -777, -722, -666, -611, -555, -500, -490, -480, -471, -461, -452, -442, -432, -423, -413, -404, -394, -384, -375, -365, -356, -346, -336, -327, -317, -308, -298, -288, -279, -269, -260, -250, -241, -231, -221, -212, -202, -193, -183, -173, -164, -154, -145, -135, -125, -116, -106, -97, -87, -77, -68, -58, -49, -39, -30, 30, 39, 49, 58, 68, 77, 87, 97, 106, 116, 125, 135, 145, 154, 164, 173, 183, 193, 202, 212, 221, 231, 241, 250, 260, 269, 279, 288, 298, 308, 317, 327, 336, 346, 356, 365, 375, 384, 394, 404, 413, 423, 432, 442, 452, 461, 471, 480, 490, 500, 555, 611, 666, 722, 777, 833, 888, 944, 1000} |

| | |
|---|---|
| A15-C8 | [10] {-1000, -944, -888, -833, -777, -722, -666, -611, -555, -500, -490, -480, -471, -461, -452, -442, -432, -423, -413, -404, -394, -384, -375, -365, -356, -346, -336, -327, -317, -308, -298, -288, -279, -269, -260, -250, -241, -231, -221, -212, -202, -193, -183, -173, -164, -154, -145, -135, -125, -116, -106, -97, -87, -77, -68, -58, -49, -39, -30, 30, 39, 49, 58, 68, 77, 87, 97, 106, 116, 125, 135, 145, 154, 164, 173, 183, 193, 202, 212, 221, 231, 241, 250, 260, 269, 279, 288, 298, 308, 317, 327, 336, 346, 356, 365, 375, 384, 394, 404, 413, 423, 432, 442, 452, 461, 471, 480, 490, 500, 555, 611, 666, 722, 777, 833, 888, 944, 1000}<br>[20] {-1000, -944, -888, -833, -777, -722, -666, -611, -555, -500, -490, -480, -471, -461, -452, -442, -432, -423, -413, -404, -394, -384, -375, -365, -356, -346, -336, -327, -317, -308, -298, -288, -279, -269, -260, -250, -241, -231, -221, -212, -202, -193, -183, -173, -164, -154, -145, -135, -125, -116, -106, -97, -87, -77, -68, -58, -49, -39, -30, 30, 39, 49, 58, 68, 77, 87, 97, 106, 116, 125, 135, 145, 154, 164, 173, 183, 193, 202, 212, 221, 231, 241, 250, 260, 269, 279, 288, 298, 308, 317, 327, 336, 346, 356, 365, 375, 384, 394, 404, 413, 423, 432, 442, 452, 461, 471, 480, 490, 500, 555, 611, 666, 722, 777, 833, 888, 944, 1000}<br>[40] {-1000, -944, -888, -833, -777, -722, -666, -611, -555, -500, -490, -480, -471, -461, -452, -442, -432, -423, -413, -404, -394, -384, -375, -365, -356, -346, -336, -327, -317, -308, -298, -288, -279, -269, -260, -250, -241, -231, -221, -212, -202, -193, -183, -173, -164, -154, -145, -135, -125, -116, -106, -97, -87, -77, -68, -58, -49, -39, -30, 30, 39, 49, 58, 68, 77, 87, 97, 106, 116, 125, 135, 145, 154, 164, 173, 183, 193, 202, 212, 221, 231, 241, 250, 260, 269, 279, 288, 298, 308, 317, 327, 336, 346, 356, 365, 375, 384, 394, 404, 413, 423, 432, 442, 452, 461, 471, 480, 490, 500, 555, 611, 666, 722, 777, 833, 888, 944, 1000} |
| $m^6$A6-C2 | [10] {-1000, -979, -959, -939, -919, -898, -878, -858, -838, -818, -797, -777, -757, -737, -717, -696, -676, -656, -636, -616, -595, -575, -555, -535, -515, -494, -474, -454, -434, -414, -393, -373, -353, -333, -313, -292, -272, -252, -232, -212, -191, -171, -151, -131, -111, -90, -70, -50, -30, -10, 10, 30, 50, 70, 90, 111, 131, 151, 171, 191, 212, 232, 252, 272, 292, 313, 333, 353, 373, 393, 414, 434, 454, 474, 494, 515, 535, 555, 575, 595, 616, 636, 656, 676, 696, 717, 737, 757, 777, 797, 818, 838, 858, 878, 898, 919, 939, 959, 979, 1000}<br>[20] {-1000, -979, -959, -939, -919, -898, -878, -858, -838, -818, -797, -777, -757, -737, -717, -696, -676, -656, -636, -616, -595, -575, -555, -535, -515, -494, -474, -454, -434, -414, -393, -373, -353, -333, -313, -292, -272, -252, -232, -212, -191, -171, -151, -131, -111, -90, -70, -50, -30, -10, 10, 30, 50, 70, 90, 111, 131, 151, 171, 191, 212, 232, 252, 272, 292, 313, 333, 353, 373, 393, 414, 434, 454, 474, 494, 515, 535, 555, 575, 595, 616, 636, 656, 676, 696, 717, 737, 757, 777, 797, 818, 838, 858, 878, 898, 919, 939, 959, 979, 1000}<br>[30] {-1000, -979, -959, -939, -919, -898, -878, -858, -838, -818, -797, -777, -757, -737, -717, -696, -676, -656, -636, -616, -595, -575, -555, -535, -515, -494, -474, -454, -434, -414, -393, -373, -353, -333, -313, -292, -272, -252, -232, -212, -191, -171, -151, -131, -111, -90, -70, -50, -30, -10, 10, 30, 50, 70, 90, 111, 131, 151, 171, 191, 212, 232, 252, 272, 292, 313, 333, 353, 373, 393, 414, 434, 454, 474, 494, 515, 535, 555, 575, 595, 616, 636, 656, 676, 696, 717, 737, 757, 777, 797, 818, 838, 858, 878, 898, 919, 939, 959, 979, 1000}<br>[40] {-1000, -979, -959, -939, -919, -898, -878, -858, -838, -818, -797, -777, -757, -737, -717, -696, -676, -656, -636, -616, -595, -575, -555, -535, -515, -494, -474, -454, -434, -414, -393, -373, -353, -333, -313, -292, -272, -252, -232, -212, -191, -171, -151, -131, -111, -90, -70, -50, -30, -10, 10, 30, 50, 70, 90, 111, 131, 151, 171, 191, 212, 232, 252, 272, 292, 313, 333, 353, 373, 393, 414, 434, 454, 474, 494, 515, 535, 555, 575, 595, 616, 636, 656, 676, 696, 717, 737, 757, 777, 797, 818, 838, 858, 878, 898, 919, 939, 959, 979, 1000}<br>[50] {-1000, -979, -959, -939, -919, -898, -878, -858, -838, -818, -797, -777, -757, -737, -717, -696, -676, -656, -636, -616, -595, -575, -555, -535, -515, -494, -474, -454, -434, -414, -393, -373, -353, -333, -313, -292, -272, -252, -232, -212, -191, -171, -151, -131, -111, -90, -70, -50, -30, -10, 10, 30, 50, 70, 90, 111, 131, 151, 171, 191, 212, 232, 252, 272, 292, 313, 333, 353, 373, 393, 414, 434, 454, 474, 494, 515, 535, 555, 575, 595, 616, 636, 656, 676, 696, 717, 737, 757, 777, 797, 818, 838, 858, 878, 898, 919, 939, 959, 979, 1000}<br>[70] {-1000, -979, -959, -939, -919, -898, -878, -858, -838, -818, -797, -777, -757, -737, -717, -696, -676, -656, -636, -616, -595, -575, -555, -535, -515, -494, -474, -454, -434, -414, -393, -373, -353, -333, -313, -292, -272, -252, -232, -212, -191, -171, -151, -131, -111, -90, -70, -50, -30, -10, 10, 30, 50, 70, 90, 111, 131, 151, 171, 191, 212, 232, 252, 272, 292, 313, 333, 353, 373, 393, 414, 434, 454, 474, 494, 515, 535, 555, 575, 595, 616, 636, 656, 676, 696, 717, 737, 757, 777, 797, 818, 838, 858, 878, 898, 919, 939, 959, 979, 1000}<br>[100] {-1000, -979, -959, -939, -919, -898, -878, -858, -838, -818, -797, -777, -757, -737, -717, -696, -676, -656, -636, -616, -595, -575, -555, -535, -515, -494, -474, -454, -434, -414, -393, -373, -353, -333, -313, -292, -272, -252, -232, -212, -191, -171, -151, -131, -111, -90, -70, -50, -30, -10, 10, 30, 50, 70, 90, 111, 131, 151, 171, 191, 212, 232, 252, 272, 292, 313, 333, 353, 373, 393, 414, |

| | |
|---|---|
| | 434, 454, 474, 494, 515, 535, 555, 575, 595, 616, 636, 656, 676, 696, 717, 737, 757, 777, 797, 818, 838, 858, 878, 898, 919, 939, 959, 979, 1000} |
| $m^6$A6-C8 | [10] {-1000, -944, -888, -833, -777, -722, -666, -611, -555, -500, -490, -480, -471, -461, -452, -442, -432, -423, -413, -404, -394, -384, -375, -365, -356, -346, -336, -327, -317, -308, -298, -288, -279, -269, -260, -250, -241, -231, -221, -212, -202, -193, -183, -173, -164, -154, -145, -135, -125, -116, -106, -97, -87, -77, -68, -58, -49, -39, -30, 30, 39, 49, 58, 68, 77, 87, 97, 106, 116, 125, 135, 145, 154, 164, 173, 183, 193, 202, 212, 221, 231, 241, 250, 260, 269, 279, 288, 298, 308, 317, 327, 336, 346, 356, 365, 375, 384, 394, 404, 413, 423, 432, 442, 452, 461, 471, 480, 490, 500, 555, 611, 666, 722, 777, 833, 888, 944, 1000}<br>[20] {-1000, -944, -888, -833, -777, -722, -666, -611, -555, -500, -490, -480, -471, -461, -452, -442, -432, -423, -413, -404, -394, -384, -375, -365, -356, -346, -336, -327, -317, -308, -298, -288, -279, -269, -260, -250, -241, -231, -221, -212, -202, -193, -183, -173, -164, -154, -145, -135, -125, -116, -106, -97, -87, -77, -68, -58, -49, -39, -30, 30, 39, 49, 58, 68, 77, 87, 97, 106, 116, 125, 135, 145, 154, 164, 173, 183, 193, 202, 212, 221, 231, 241, 250, 260, 269, 279, 288, 298, 308, 317, 327, 336, 346, 356, 365, 375, 384, 394, 404, 413, 423, 432, 442, 452, 461, 471, 480, 490, 500, 555, 611, 666, 722, 777, 833, 888, 944, 1000}<br>[40] {-1000, -944, -888, -833, -777, -722, -666, -611, -555, -500, -490, -480, -471, -461, -452, -442, -432, -423, -413, -404, -394, -384, -375, -365, -356, -346, -336, -327, -317, -308, -298, -288, -279, -269, -260, -250, -241, -231, -221, -212, -202, -193, -183, -173, -164, -154, -145, -135, -125, -116, -106, -97, -87, -77, -68, -58, -49, -39, -30, 30, 39, 49, 58, 68, 77, 87, 97, 106, 116, 125, 135, 145, 154, 164, 173, 183, 193, 202, 212, 221, 231, 241, 250, 260, 269, 279, 288, 298, 308, 317, 327, 336, 346, 356, 365, 375, 384, 394, 404, 413, 423, 432, 442, 452, 461, 471, 480, 490, 500, 555, 611, 666, 722, 777, 833, 888, 944, 1000} |
| $m^6$A6-C10 | [25] {-805, -785, -765, -745, -724, -704, -684, -664, -643, -623, -603, -583, -562, -542, -522, -502, -481, -461, -441, -421, -400, -380, -360, -340, -319, -299, -279, -259, -238, -218, -198, -178, -157, -137, -117, -96, -76, -56, -36, -15, 4, 24, 44, 65, 85, 105, 125, 146, 166, 186, 206, 227, 247, 267, 287, 308, 328, 348, 368, 389, 409, 429, 449, 470, 490, 510, 530, 551, 571, 591, 611, 632, 652, 672, 692, 713, 733, 753, 773, 794}<br>[50] {-806, -785, -765, -745, -725, -704, -684, -664, -644, -623, -603, -583, -563, -542, -522, -502, -481, -461, -441, -421, -400, -380, -360, -340, -319, -299, -279, -259, -238, -218, -198, -178, -157, -137, -117, -97, -76, -56, -36, 24, 44, 64, 85, 105, 125, 145, 166, 186, 206, 226, 247, 267, 287, 307, 328, 348, 368, 388, 409, 429, 449, 469, 490, 510, 530, 550, 571, 591, 611, 631, 652, 672, 692, 712, 733, 753, 773, 793} |
| **dsGGACU, 37°C, $Mg^{2+}$** | |
| A6-C2 | [10] {-1000, -922, -844, -766, -688, -611, -533, -455, -377, -300, -290, -281, -272, -262, -253, -244, -234, -225, -216, -206, -197, -188, -178, -169, -160, -151, -141, -132, -123, -113, -104, -95, -85, -76, -67, -57, -48, -39, -30, 30, 39, 49, 58, 68, 77, 87, 97, 106, 116, 125, 135, 145, 154, 164, 173, 183, 193, 202, 212, 221, 231, 241, 250, 260, 269, 279, 288, 298, 308, 317, 327, 336, 346, 356, 365, 375, 384, 394, 404, 413, 423, 432, 442, 452, 461, 471, 480, 490, 500, 555, 611, 666, 722, 777, 833, 888, 944, 1000}<br>[20] {-1000, -922, -844, -766, -688, -611, -533, -455, -377, -300, -290, -281, -272, -262, -253, -244, -234, -225, -216, -206, -197, -188, -178, -169, -160, -151, -141, -132, -123, -113, -104, -95, -85, -76, -67, -57, -48, -39, -30, 30, 39, 49, 58, 68, 77, 87, 97, 106, 116, 125, 135, 145, 154, 164, 173, 183, 193, 202, 212, 221, 231, 241, 250, 260, 269, 279, 288, 298, 308, 317, 327, 336, 346, 356, 365, 375, 384, 394, 404, 413, 423, 432, 442, 452, 461, 471, 480, 490, 500, 555, 611, 666, 722, 777, 833, 888, 944, 1000}<br>[40] {-1000, -922, -844, -766, -688, -611, -533, -455, -377, -300, -290, -281, -272, -262, -253, -244, -234, -225, -216, -206, -197, -188, -178, -169, -160, -151, -141, -132, -123, -113, -104, -95, -85, -76, -67, -57, -48, -39, -30, 30, 39, 49, 58, 68, 77, 87, 97, 106, 116, 125, 135, 145, 154, 164, 173, 183, 193, 202, 212, 221, 231, 241, 250, 260, 269, 279, 288, 298, 308, 317, 327, 336, 346, 356, 365, 375, 384, 394, 404, 413, 423, 432, 442, 452, 461, 471, 480, 490, 500, 555, 611, 666, 722, 777, 833, 888, 944, 1000} |
| A6-C8 | [10] {-1000, -922, -844, -766, -688, -611, -533, -455, -377, -300, -290, -281, -272, -262, -253, -244, -234, -225, -216, -206, -197, -188, -178, -169, -160, -151, -141, -132, -123, -113, -104, -95, -85, -76, -67, -57, -48, -39, -30, 30, 39, 49, 58, 68, 77, 87, 97, 106, 116, 125, 135, 145, 154, 164, 173, 183, 193, 202, 212, 221, 231, 241, 250, 260, 269, 279, 288, 298, 308, 317, 327, 336, 346, 356, 365, 375, 384, 394, 404, 413, 423, 432, 442, 452, 461, 471, 480, 490, 500, 555, 611, 666, 722, 777, 833, 888, 944, 1000}<br>[20] {-1000, -922, -844, -766, -688, -611, -533, -455, -377, -300, -290, -281, -272, -262, -253, -244, -234, -225, -216, -206, -197, -188, -178, -169, -160, -151, -141, -132, -123, -113, -104, -95, -85, -76, -67, -57, -48, -39, -30, 30, 39, 49, 58, 68, 77, 87, 97, 106, 116, 125, 135, 145, 154, 164, 173, 183, 193, 202, 212, 221, 231, 241, 250, 260, 269, 279, 288, 298, 308, 317, 327, 336, 346, 356, 365, 375, 384, 394, 404, 413, 423, 432, 442, 452, 461, 471, 480, 490, 500, 555, 611, 666, 722, 777, 833, 888, 944, 1000}<br>[40] {-1000, -922, -844, -766, -688, -611, -533, -455, -377, -300, -290, -281, -272, -262, -253, -244, -234, -225, -216, -206, -197, -188, -178, -169, -160, -151, -141, -132, -123, -113, -104, -95, -85, -76, -67, -57, |

| | |
|---|---|
| | -48, -39, -30, 30, 39, 49, 58, 68, 77, 87, 97, 106, 116, 125, 135, 145, 154, 164, 173, 183, 193, 202, 212, 221, 231, 241, 250, 260, 269, 279, 288, 298, 308, 317, 327, 336, 346, 356, 365, 375, 384, 394, 404, 413, 423, 432, 442, 452, 461, 471, 480, 490, 500, 555, 611, 666, 722, 777, 833, 888, 944, 1000} |
| **dsGGACU$^{m6A}$, 37°C, Mg$^{2+}$** | |
| m$^6$A6-C2 | [10] {-1000, -922, -844, -766, -688, -611, -533, -455, -377, -300, -290, -281, -272, -262, -253, -244, -234, -225, -216, -206, -197, -188, -178, -169, -160, -151, -141, -132, -123, -113, -104, -95, -85, -76, -67, -57, -48, -39, -30, 30, 39, 49, 58, 68, 77, 87, 97, 106, 116, 125, 135, 145, 154, 164, 173, 183, 193, 202, 212, 221, 231, 241, 250, 260, 269, 279, 288, 298, 308, 317, 327, 336, 346, 356, 365, 375, 384, 394, 404, 413, 423, 432, 442, 452, 461, 471, 480, 490, 500, 555, 611, 666, 722, 777, 833, 888, 944, 1000}<br>[20] {-1000, -922, -844, -766, -688, -611, -533, -455, -377, -300, -290, -281, -272, -262, -253, -244, -234, -225, -216, -206, -197, -188, -178, -169, -160, -151, -141, -132, -123, -113, -104, -95, -85, -76, -67, -57, -48, -39, -30, 30, 39, 49, 58, 68, 77, 87, 97, 106, 116, 125, 135, 145, 154, 164, 173, 183, 193, 202, 212, 221, 231, 241, 250, 260, 269, 279, 288, 298, 308, 317, 327, 336, 346, 356, 365, 375, 384, 394, 404, 413, 423, 432, 442, 452, 461, 471, 480, 490, 500, 555, 611, 666, 722, 777, 833, 888, 944, 1000}<br>[40] {-1000, -922, -844, -766, -688, -611, -533, -455, -377, -300, -290, -281, -272, -262, -253, -244, -234, -225, -216, -206, -197, -188, -178, -169, -160, -151, -141, -132, -123, -113, -104, -95, -85, -76, -67, -57, -48, -39, -30, 30, 39, 49, 58, 68, 77, 87, 97, 106, 116, 125, 135, 145, 154, 164, 173, 183, 193, 202, 212, 221, 231, 241, 250, 260, 269, 279, 288, 298, 308, 317, 327, 336, 346, 356, 365, 375, 384, 394, 404, 413, 423, 432, 442, 452, 461, 471, 480, 490, 500, 555, 611, 666, 722, 777, 833, 888, 944, 1000} |
| m$^6$A6-C8 | [10] {-1000, -922, -844, -766, -688, -611, -533, -455, -377, -300, -290, -281, -272, -262, -253, -244, -234, -225, -216, -206, -197, -188, -178, -169, -160, -151, -141, -132, -123, -113, -104, -95, -85, -76, -67, -57, -48, -39, -30, 30, 39, 49, 58, 68, 77, 87, 97, 106, 116, 125, 135, 145, 154, 164, 173, 183, 193, 202, 212, 221, 231, 241, 250, 260, 269, 279, 288, 298, 308, 317, 327, 336, 346, 356, 365, 375, 384, 394, 404, 413, 423, 432, 442, 452, 461, 471, 480, 490, 500, 555, 611, 666, 722, 777, 833, 888, 944, 1000}<br>[20] {-1000, -922, -844, -766, -688, -611, -533, -455, -377, -300, -290, -281, -272, -262, -253, -244, -234, -225, -216, -206, -197, -188, -178, -169, -160, -151, -141, -132, -123, -113, -104, -95, -85, -76, -67, -57, -48, -39, -30, 30, 39, 49, 58, 68, 77, 87, 97, 106, 116, 125, 135, 145, 154, 164, 173, 183, 193, 202, 212, 221, 231, 241, 250, 260, 269, 279, 288, 298, 308, 317, 327, 336, 346, 356, 365, 375, 384, 394, 404, 413, 423, 432, 442, 452, 461, 471, 480, 490, 500, 555, 611, 666, 722, 777, 833, 888, 944, 1000}<br>[40] {-1000, -922, -844, -766, -688, -611, -533, -455, -377, -300, -290, -281, -272, -262, -253, -244, -234, -225, -216, -206, -197, -188, -178, -169, -160, -151, -141, -132, -123, -113, -104, -95, -85, -76, -67, -57, -48, -39, -30, 30, 39, 49, 58, 68, 77, 87, 97, 106, 116, 125, 135, 145, 154, 164, 173, 183, 193, 202, 212, 221, 231, 241, 250, 260, 269, 279, 288, 298, 308, 317, 327, 336, 346, 356, 365, 375, 384, 394, 404, 413, 423, 432, 442, 452, 461, 471, 480, 490, 500, 555, 611, 666, 722, 777, 833, 888, 944, 1000} |
| **dsHCV, 55°C, Mg$^{2+}$** | |
| A11-C8 | [10] {-1000, -922, -844, -766, -688, -611, -533, -455, -377, -300, -290, -281, -272, -262, -253, -244, -234, -225, -216, -206, -197, -188, -178, -169, -160, -151, -141, -132, -123, -113, -104, -95, -85, -76, -67, -57, -48, -39, -30, 30, 39, 49, 58, 68, 77, 87, 97, 106, 116, 125, 135, 145, 154, 164, 173, 183, 193, 202, 212, 221, 231, 241, 250, 260, 269, 279, 288, 298, 308, 317, 327, 336, 346, 356, 365, 375, 384, 394, 404, 413, 423, 432, 442, 452, 461, 471, 480, 490, 500, 555, 611, 666, 722, 777, 833, 888, 944, 1000}<br>[20] {-1000, -922, -844, -766, -688, -611, -533, -455, -377, -300, -290, -281, -272, -262, -253, -244, -234, -225, -216, -206, -197, -188, -178, -169, -160, -151, -141, -132, -123, -113, -104, -95, -85, -76, -67, -57, -48, -39, -30, 30, 39, 49, 58, 68, 77, 87, 97, 106, 116, 125, 135, 145, 154, 164, 173, 183, 193, 202, 212, 221, 231, 241, 250, 260, 269, 279, 288, 298, 308, 317, 327, 336, 346, 356, 365, 375, 384, 394, 404, 413, 423, 432, 442, 452, 461, 471, 480, 490, 500, 555, 611, 666, 722, 777, 833, 888, 944, 1000}<br>[40] {-1000, -922, -844, -766, -688, -611, -533, -455, -377, -300, -290, -281, -272, -262, -253, -244, -234, -225, -216, -206, -197, -188, -178, -169, -160, -151, -141, -132, -123, -113, -104, -95, -85, -76, -67, -57, -48, -39, -30, 30, 39, 49, 58, 68, 77, 87, 97, 106, 116, 125, 135, 145, 154, 164, 173, 183, 193, 202, 212, 221, 231, 241, 250, 260, 269, 279, 288, 298, 308, 317, 327, 336, 346, 356, 365, 375, 384, 394, 404, 413, 423, 432, 442, 452, 461, 471, 480, 490, 500, 555, 611, 666, 722, 777, 833, 888, 944, 1000} |
| **dsHCV$^{m6A}$, 55°C, Mg$^{2+}$** | |
| A11-C8 | [10] {-1000, -922, -844, -766, -688, -611, -533, -455, -377, -300, -290, -281, -272, -262, -253, -244, -234, -225, -216, -206, -197, -188, -178, -169, -160, -151, -141, -132, -123, -113, -104, -95, -85, -76, -67, -57, -48, -39, -30, 30, 39, 49, 58, 68, 77, 87, 97, 106, 116, 125, 135, 145, 154, 164, 173, 183, 193, 202, 212, 221, 231, 241, 250, 260, 269, 279, 288, 298, 308, 317, 327, 336, 346, 356, 365, 375, 384, 394, 404, 413, 423, 432, 442, 452, 461, 471, 480, 490, 500, 555, 611, 666, 722, 777, 833, 888, 944, 1000}<br>[20] {-1000, -922, -844, -766, -688, -611, -533, -455, -377, -300, -290, -281, -272, -262, -253, -244, -234, -225, -216, -206, -197, -188, -178, -169, -160, -151, -141, -132, -123, -113, -104, -95, -85, -76, -67, -57, -48, -39, -30, 30, 39, 49, 58, 68, 77, 87, 97, 106, 116, 125, 135, 145, 154, 164, 173, 183, 193, 202, 212, |

| | |
|---|---|
| | 221, 231, 241, 250, 260, 269, 279, 288, 298, 308, 317, 327, 336, 346, 356, 365, 375, 384, 394, 404, 413, 423, 432, 442, 452, 461, 471, 480, 490, 500, 555, 611, 666, 722, 777, 833, 888, 944, 1000}<br>[40] {-1000, -922, -844, -766, -688, -611, -533, -455, -377, -300, -290, -281, -272, -262, -253, -244, -234, -225, -216, -206, -197, -188, -178, -169, -160, -151, -141, -132, -123, -113, -104, -95, -85, -76, -67, -57, -48, -39, -30, 30, 39, 49, 58, 68, 77, 87, 97, 106, 116, 125, 135, 145, 154, 164, 173, 183, 193, 202, 212, 221, 231, 241, 250, 260, 269, 279, 288, 298, 308, 317, 327, 336, 346, 356, 365, 375, 384, 394, 404, 413, 423, 432, 442, 452, 461, 471, 480, 490, 500, 555, 611, 666, 722, 777, 833, 888, 944, 1000} |
| **dsA6DNA$^{m6A}$, 50°C** | |
| m$^6$A6-C2 | [10] {-1000, -922, -844, -766, -688, -611, -533, -455, -377, -300, -290, -281, -272, -262, -253, -244, -234, -225, -216, -206, -197, -188, -178, -169, -160, -151, -141, -132, -123, -113, -104, -95, -85, -76, -67, -57, -48, -39, -30, 30, 39, 49, 58, 68, 77, 87, 97, 106, 116, 125, 135, 145, 154, 164, 173, 183, 193, 202, 212, 221, 231, 241, 250, 260, 269, 279, 288, 298, 308, 317, 327, 336, 346, 356, 365, 375, 384, 394, 404, 413, 423, 432, 442, 452, 461, 471, 480, 490, 500, 555, 611, 666, 722, 777, 833, 888, 944, 1000}<br>[25] {-1000, -922, -844, -766, -688, -611, -533, -455, -377, -300, -290, -281, -272, -262, -253, -244, -234, -225, -216, -206, -197, -188, -178, -169, -160, -151, -141, -132, -123, -113, -104, -95, -85, -76, -67, -57, -48, -39, -30, 30, 39, 49, 58, 68, 77, 87, 97, 106, 116, 125, 135, 145, 154, 164, 173, 183, 193, 202, 212, 221, 231, 241, 250, 260, 269, 279, 288, 298, 308, 317, 327, 336, 346, 356, 365, 375, 384, 394, 404, 413, 423, 432, 442, 452, 461, 471, 480, 490, 500, 555, 611, 666, 722, 777, 833, 888, 944, 1000}<br>[40] {-1000, -922, -844, -766, -688, -611, -533, -455, -377, -300, -290, -281, -272, -262, -253, -244, -234, -225, -216, -206, -197, -188, -178, -169, -160, -151, -141, -132, -123, -113, -104, -95, -85, -76, -67, -57, -48, -39, -30, 30, 39, 49, 58, 68, 77, 87, 97, 106, 116, 125, 135, 145, 154, 164, 173, 183, 193, 202, 212, 221, 231, 241, 250, 260, 269, 279, 288, 298, 308, 317, 327, 336, 346, 356, 365, 375, 384, 394, 404, 413, 423, 432, 442, 452, 461, 471, 480, 490, 500, 555, 611, 666, 722, 777, 833, 888, 944, 1000} |

**Supplementary Table 7. Relaxation delay times used for measuring the exchange rate of water and imino protons.**

| Construct | Temperature (°C) | Delay time (sec) |
|---|---|---|
| dsGGACU | 25 | 0.0, 0.05, 0.1, 0.15, 0.2, 0.25, 0.3, 0.4, 0.5, 0.6, 0.7, 0.9, 1.2, 1.5, 2.0, 3.0, 4.0 |
| dsGGACU | 37 | 0.0, 0.003, 0.006, 0.01, 0.02, 0.03, 0.04, 0.05, 0.1, 0.2, 0.3, 0.5, 0.7, 1.0, 1.5, 2.0, 3.0 |
| dsGGACU$^{m6A}$ | 25 | 0.0, 0.01, 0.02, 0.05, 0.08, 0.1, 0.12, 0.15, 0.18, 0.2, 0.25, 0.3, 0.5, 0.7, 1.0, 1.5, 2.0, 3.0, 4.0 |
| dsGGACU$^{m6A}$ | 37 | 0.0, 0.003, 0.006, 0.01, 0.02, 0.03, 0.04, 0.05, 0.1, 0.2, 0.3, 0.5, 0.7, 1.0, 1.5, 2.0 |
| dsA6RNA | 25 | 0.0, 0.005, 0.01, 0.02, 0.03, 0.04, 0.05, 0.1, 0.2, 0.3, 0.5, 0.7, 1.0, 1.3, 1.6, 2.0, 2.5, 3.0, 3.5 |
| dsA6RNA | 37 | 0.0, 0.005, 0.01, 0.02, 0.03, 0.04, 0.05, 0.1, 0.2, 0.3, 0.5, 0.7, 1.0, 1.3, 1.6, 2.0, 2.5, 3.0, 3.5 |
| dsA6RNA$^{m6A}$ | 25 | 0.0, 0.005, 0.01, 0.02, 0.03, 0.04, 0.05, 0.1, 0.2, 0.3, 0.5, 0.7, 1.0, 1.3, 1.6, 2.0, 2.5, 3.0, 3.5 |
| dsA6RNA$^{m6A}$ | 37 | 0.0, 0.005, 0.01, 0.02, 0.03, 0.04, 0.05, 0.1, 0.2, 0.3, 0.5, 0.7, 1.0, 1.3, 1.6, 2.0, 2.5, 3.0, 3.5 |

### Supplementary Notes

#### Supplementary Note 1. Additional support for the methylamino group being *syn* in the ES detected in hpGGACU$^{m6A}$

The rate constants for *syn* → *anti* interconversion in dsRNA ($k_{-2}$ ~500 $s^{-1}$ at T = 55°C) and ssRNA ($k_1$ ~300 $s^{-1}$ at T = 55°C) are similar to each other (Supplementary Table 2). This is expected given that in both cases, the *syn* methylamino group is not H-bonded. Conversely, the much slower *anti* → *syn* interconversion in dsRNA ($k_2$ ~10 $s^{-1}$ at T = 55°C) relative to ssRNA ($k_{-1}$ ~6,000 $s^{-1}$ at T = 55°C) is also expected given that the *anti* methylamino group is H-bonded in dsRNA but not in ssRNA. In addition, $k_{on,syn}$ (ssRNA$^{syn}$ → dsRNA$^{syn}$) and $k_{off,syn}$ (dsRNA$^{syn}$ → ssRNA$^{syn}$) values are ~20-fold slower and ~80-fold faster than their unmethylated RNA counterparts respectively at T = 55°C (Supplementary Table 2). These large changes in hybridization kinetics relative to the unmodified duplex are in line with those reported previously for mismatches[7]. This is reasonable because one would expect the hybridization kinetics of dsRNA$^{syn}$ (ssRNA$^{syn}$⇌dsRNA$^{syn}$) to be similar to those of dsRNA containing mismatches[7], given that $m^6$A in the *syn* conformation is expected to lose at least one Watson-Crick H-bond in dsRNA.

#### Supplementary Note 2. Base opening kinetic modeling

We tested two models to examine the impact of $m^6$(*syn*)A···U on base opening kinetics,

1. $m^6$(*syn*)A···U can also contribute to solvent exchange in addition to the canonical base opening state (Extended Data Fig. 8b, model 1).
2. $m^6$(*syn*)A···U replaces the canonical base opening state and is the dominant contributor to solvent exchange (Extended Data Fig. 8b, model 2).

Kinetic simulations of these two models were performed using differential equations below to simulate the apparent solvent exchange rate constant ($k_{ex}$) of methylated RNA assuming model 1 or model 2.

Model 1:

$$\frac{\mathrm{d}[m6(anti)A-U]}{\mathrm{dt}} = -k_{op}[m6(anti)A-U] + k_{cl}[m6(anti)A-U(open)] + k_{-2}[m6(syn)A-U] - k_2[m6(anti)A-U]$$

$$\frac{\mathrm{d}[m6(anti)A-U(open)]}{\mathrm{dt}} = k_{op}[m6(anti)A-U] - k_{cl}[m6(anti)A-U(open)] - k_{e1}[m6(anti)A-U(open)]$$

$$\frac{\mathrm{d}[m6(syn)A-U]}{\mathrm{dt}} = k_2[m6(anti)A-U] - k_{-2}[m6(syn)A-U] - k_{e2}[m6(syn)A-U]$$

$$\frac{\mathrm{d}[Exchanged\ H]}{\mathrm{dt}} = k_{e1}[m6(anti)A-U(open)] + k_{e2}[m6(syn)A-U]$$

Model 2:

$$\frac{\mathrm{d}[m6(anti)A-U]}{\mathrm{dt}} = k_{-2}[m6(syn)A-U] - k_2[m6(anti)A-U]$$

$$\frac{\mathrm{d}[m6(syn)A-U]}{\mathrm{dt}} = -k_{-2}[m6(syn)A-U] + k_2[m6(anti)A-U] - k_{e2}[m6(syn)A-U]$$

$$\frac{\mathrm{d}[Exchanged\ H]}{\mathrm{dt}} = k_{e2}[m6(syn)A-U]$$

$k_{op}$ and $k_{cl}$ are the opening and closing rate constants of $m^6$(*anti*)A-U bp, respectively and were assumed to be equal to those of canonical base opening in unmethylated A-U bp (700 $s^{-1}$ and $1.7 \times 10^7$ $s^{-1}$ respectively[8]). $k_2$, $k_{-2}$ were methylamino isomerization rate constants in dsRNA measured using RD. $k_{e1}$ and $k_{e2}$ are the intrinsic imino proton exchange rate constants for $m^6$(*anti*)A···U(open) and $m^6$(*syn*)A···U, respectively (Extended Data Fig. 8b) and were both assumed to be equal to the corresponding rate of unmethylated A-U base-open state ($k_{ex,open}$, Extended Data Fig. 8a), which is $10^6$ $s^{-1}$ based on a prior study[9]. $[Exchanged\ H]$ was simulated at 100 evenly distributed time points from 0 s to 0.5 s for model 1, 0 s to 5 s for model 2. The simulated $[Exchanged\ H]$ at multiple time points was then fit to

$$[Exchanged\ H] = A\left(1 - e^{-k_{ex,m6A}^{app}t}\right)$$

Where $t$ is time, $k_{ex,m6A}^{app}$ is the apparent base opening exchange rate constant for methylated RNA. A is a pre-exponential factor. The fitted $k_{ex,m6A}^{app}$ was then compared with that of unmethylated RNA ($k_{\mathrm{ex}} = \frac{k_{op}k_{ex,open}}{k_{cl}+k_{ex,open}}$) to calculate the $k_{\mathrm{ex}}$ fold-change ($k_{ex,m6A}^{app}/k_{\mathrm{ex}}$) shown in Extended Data Fig. 8d. The simulation results suggest that $m^6A$ has little effect on $k_{\mathrm{ex}}$ in model 1 while it slows $k_{\mathrm{ex}}$ by ~200-fold in model 2. We did not observe a significant effect of $m^6A$ on $k_{\mathrm{ex}}$ (Extended Data Fig. 8c), consistent with model 1. Therefore, the $m^6$(*syn*)A···U does not replace the canonical base-open state. However, we cannot rule out that $m^6A$ can be an alternative base-open state.

**Supplementary Note 3. RREIIB RD analysis**

We previously showed that methylating A68 has minor effects on the ES1 + ES2 populations in a different three-way junction context in the absence and presence of $Mg^{2+}$ based on analysis of $^{15}N$ site-labelled U72 imino resonance[10]. The RD data (Extended Data Fig. 10d) measured here for the RREIIB stem-loop in the absence of $Mg^{2+}$ indicates that $m^6A$ increases $p_{\mathrm{ES1}}$ by ~3-fold (Supplementary Table 1). This could be due to improved stacking interactions when $m^6A$ is near flexible sites. The data also show that $m^6A$ diminishes the RD contribution from ES2, possibly due to destabilization and/or reduction of $k_{\mathrm{ex}}$ when forming the ES2 $m^6$A68(*anti*)-G50(*anti*) mismatch[10]. The impact of $m^6A$ on the measured RREIIB RD was weaker in the presence of 3 mM $Mg^{2+}$ (Extended Data Fig. 10d) although it was not possible to reliably determine the ES1 exchange parameters due to weak RD. NMR RD studies are needed to examine whether $m^6A$ impacts the kinetics of ES1 and/or ES2 exchange in the three-way junction context.

**Supplementary Note 4. Synthesis of ($^{13}$C10)-m$^6$A RNA phosphoramidite.**

Detailed description of synthetic protocols is shown in Supplementary Fig. 1a.

*3',5'-O-bis(t-butylsilyl)-2'-O-(t-butyldimethylsilyl)-N-6-$^{13}$C-methyladenosine (2)*

Compound **1** (1.00 g, 1.85 mmol) was dissolved in anhydrous N,N-dimethylformamide (10 ml) under argon atmosphere. Then $^{13}$C-methylamine hydrochloride (2.0 eq, 253 mg, 3.70 mmol) and triethylamine (2.0 eq, 515 µl, 3.70 mmol) were added and the mixture was stirred at room temperature for 3 days. After complete conversion of the starting material was indicated by TLC analysis (EtOAc/n-hexane = 6/4), the solvent was evaporated under high vacuum. The residue was dissolved in ethyl acetate, washed once with saturated sodium bicarbonate solution and once with brine. The organic layer was collected, dried over anhydrous sodium sulfate, filtered and evaporated. Final purification via silica flash column chromatography using a gradient of 20% to 70% ethyl acetate in n-hexane to give pure compound **2** as white foam. The product was dired under high vacuum.

Yield: 782 mg (1.46 mmol, 79 %)

TLC: EtOAc/n-hexane = 6/4; $R_f$ = 0.4

$^1$H-NMR (300 MHz, $CDCl_3$, 25°C): δ 8.36 (s, 1H, C(2)*H*); 7.75 (s, 1H, C(8)*H*); 5.90 (s, 1H, C(1')*H*); 5.87 (s, 1H, N*H*); 4.62 (d, 1H, C(2')*H*); 4.55 (dd, 1H, C(3')*H*); 4.47 (dd, 1H, C(5')*H*); 4.20 (m, 1H, C(4')*H*); 4.02 (dd, C(5'')*H*); 3.2 (dd, 3H, $^1J_{CH}$ = 138.8 Hz, NH$^{13}$C*H$_3$*); 1.08 (s, 9H, tBu); 1.04 (s, 9H, tBu); 0.92 (s, 9H, tBu); 0.15 (s, 3H, $SiCH_3$); 0.14 (s, 3H, $SiCH_3$) ppm.

$^{13}$C-NMR (75 MHz, $CDCl_3$, 25°C): δ 153.5 (C(2)); 138.1 (C(8)); 92.6 (C(1')); 76.0 (C(3')); 75.6 (C(2')); 74.8 (C(4')); 68.0 (C(5')); 27.8 ($^{13}CH_3$); 27.6 ($CH_3$, tBu); 27.2 ($CH_3$, tBu); 26.0 ($CH_3$, tBu); -4.2 ($SiCH_3$); -4.8 ($SiCH_3$) ppm.

*2'-O-(t-butyldimethylsilyl)-N-6-$^{13}$C-methyladenosine (3)*

Compound **2** (782 mg, 1.46 mmol) was dissolved in anhydrous dichloromethane (10 ml) under argon atmosphere and cooled to 0°C using an ice bath. After hydrogen fluoride pyridine (70 % HF in pyridine, 3.2 eq, 122 µl, 4.70 mmol) was diluted with anhydrous pyridine (700 µl) in an ice cooled Eppendorf tube, the mixture was added to the stirred solution of **2**. After stirring under argon at 0 °C for 3 hours completion of the reaction was indicated via TLC analysis (EtOAc/n-hexane = 7/3). The mixture was diluted with dichloromethane, washed once with water and once with saturated sodium bicarbonate solution. The organic layer was collected, dried over anhydrous sodium sulfate, filtered and evaporated. The crude yellowish foam of **3** was used for the next synthesis step without further purification.

Yield: quantitative

TLC: EtOAc/n-hexane = 7/3; $R_f$ = 0.2

*5'-O-(4,4'-dimethoxytrityl)-2'-O-(t-butyldimethylsilyl)-N-6-$^{13}$C-methyladenosine (4)*

Crude Compound **3** (577 mg, 1.46 mmol) together with a spatula tip 4-(dimethyl- amino)pyridine was coevaporated trice with anhydrous pyridine and subsequently dissolved in anhydrous pyridine (20 ml) under argon atmosphere. Then 4,4'-dimethoxytrityl chloride (1.1 eq, 542 mg, 1.60 mmol) was added to the solution in three portions within 1 hour and the mixture was stirred for 3 hours at room temperature. After TLC analysis ($CH_2Cl_2$/MeOH = 9/1) showed full conversion of the starting material the reaction was quenched with methanol (1 ml) and evaporated to an oily residue which was coevaporated twice with toluene. The residue was dissolved in dichloromethane and washed twice with 5 % citric acid and twice with saturated sodium bicarbonate solution. The organic layer was collected, dried over sodium sulfate, filtered and evaporated. The crude product was applied to a silica gel column with methylene chloride and eluted using a gradient from 0 to 3% of methanol in dichloromethane to give compound **4** as slightly yellow foam. The product was dried under high vacuum.

Yield: 920 mg (1.32 mmol, 90 % referred to **2**)

TLC: $CH_2Cl_2$/MeOH = 9/1; $R_f$ = 0.8

$^1$H-NMR (300 MHz, DMSO-$d_6$, 25°C): δ 8.25 (s, 1H, C(8)*H*); 8.17 (s, 1H, C(2)*H*); 7.75 (s, 1H, N*H*); 7.44-6.80 (m, 13H, C*H* arom); 5.95 (d, 1H, C(1')*H*); 5.10 (d, 1H, C(3')O*H*); 4.85 (m, 1H, C(2')H); 4.26 (m, 1H, C(3')H); 4.10 (m, 1H, C(4')H); 3.73 (s, 6H, OC$H_3$); 3.27 (m, 2H, C(5')*H*; C(5'')*H*); 2.97 (d, 3H, $^1J_{CH}$ = 138.4 Hz, $^{13}CH_3$); 0.75 (s, 9H, tBu); -0.04 (s, 3H, SiC$H_3$); -0.14 (s, 3H, SiC$H_3$) ppm.

$^{13}$C-NMR (75 MHz, DMSO-$d_6$, 25°C): δ 151.6 (C(2)); 138.2 (C(8)); 130.8-110.5 (C arom); 87.4 (C(1')); 82.9 (C(4')); 74.4 (C(2')); 70.1 (C(3')); 63.1 (C(5')); 54.7 ($OCH_3$); 26.7 ($^{13}CH_3$); 25.3 ($CH_3$, tBu); -4.61 ($SiCH_3$); -5.3 ($SiCH_3$) ppm.

*5'-O-(4,4'-dimethoxytrityl)-2'-O-(t-butyldimethylsilyl)-N-6-$^{13}$C-methyladenosine 3'-[(2-cyanoethyl)-(N,N-diisopropyl)]phosphoramidite (5)*

Compound **4** (920 mg, 1.32 mmol) was dried under high vacuum over night and dissolved in 10 mL of dry tetrahydrofuran under argon atmosphere. To this solution was added simultaneously 2-cyanoethyl-N,N-diisopropylchlorophosphoramidite (1.1 eq, 343 mg, 1.45 mmol) and N,N-diisopropylethylamine (5.0 eq, 1.15 ml, 6.58 mmol) and stirred 2 hours at room temperature. After TLC analysis (EtOAc + 1 % $NEt_3$) indicated ful conversion the reaction was quneched by addition of methanol (1 ml). The solution was diluted with chloroform and washed once with half saturated sodium bicarbonate solution. The organic layer was dried over anhydrous sodium sulfate, filtered and evaporated to dryness. The crude product purified with silica gel flash chromatography using a gradient from 2:8 to 7:3 ethyl acetate/n-hexane (+2 % triethylamine) to give **5** as a colorless foam. The product was dried in high vacuum.

Yield: 750 mg (0.83 mmol, 63 %)

TLC: EtOAc + 1% $NEt_3$; $R_f$ = 0.7

$^1$H-NMR (300 MHz, $CDCl_3$, 25°C): δ 8.30 (d, 1H, C(2)*H*); 7.96 (d, 1H, C(8)*H*); 7.50-6.76 (m, 13H, C*H* arom); 5.98 (m, 1H, C(1')*H*); 5.86 (s, 1H, N*H*); 5.07 (m, 1H, C(2')*H*); 4.40 (m, 1H, C(3')*H*); 4.33 (m, 1H, C(4')*H*); 3.92 (m, 1H, POC*H₂*); 3.78 (s, 6H, OC*H₃*); 3.63 (m, 1H, POC*H₂*); 3.59 (m, 2H, C*H* iPr); 3.55 (m, 1H, C(5')*H*); 3.31 (C(5'')*H*); 3.18 (d, 3H, $^1J_{CH}$ = 138.4 Hz, $^{13}CH_3$); 2.65 (t, 1H, $CH_2$CN); 2.30 (t, 1H, $CH_2$CN); 1.21-1.00 (m, 12H, C*H*3 iPr); 0.76 (d, 9H, C*H*3 tBu); -0.05 (d, 3H, Si$CH_3$); -0.20 (d, 3H, Si$CH_3$) ppm.

$^{13}$C-NMR (75 MHz, $CDCl_3$, 25°C): δ 153.1 (C(2)); 139.0 (C(8)); 132.0-113.3 (C arom); 88.4 (C(1')); 84.1 (C(4')); 74.9 (C(2')); 73.4 (C(3')); 63.8 (C(5')); 58.3 (POCH2); 55.5 (OCH3); 43.5 (CH iPr); 27.7 (13CH3); 26.2 (CH3 tBu); 25.3 (CH3 iPr), 20.3 (CH2CN); -3.7 (SiCH3); -4.6 (SiCH3) ppm.

$^{31}$P-NMR (121 MHz, CDCl3, 25°C): δ 151.5, 149.5 ppm.

**Supplementary Note 5. Synthesis of $^{13}$C8,$^{13}$C2-labeled m$^6$dA phosphoramidite.**

Detailed description of synthetic protocols is shown in Supplementary Fig. 1b.

*2',3',5'-Tribenzoyl-(2,8-$^{13}$C)-6-chloroadenosine (2)*

Compound **1** (2.00 g, 3.43 mmol) was suspended in N,N-dimethylaniline (1.05 eq, 453 µl, 3.60 mmol) and $POCl_3$ (21 eq, 6,6 ml, 72.10 mmol) and stirred for 7 minutes at room temperature under argon atmosphere. The resulting solution was refluxed for 13 minutes at 120 °C using a preheated oil bath. After cooling the flask to room temperature using an ice bath the mixture was poured over crushed ice and stirred together with dichloromethane (approx 50 ml) in an 250 ml erlenmeyer flask. The phases were separated and the aqueous layer extracted trice with dichloromethane. The combined organic layers were washed twice with 2M HCl, twice with saturated bicarbonate solution, once with brine, dried over $Na_2SO_4$, filtered and evaporated to give compound **2** as brown foam. TLC analysis ($CH_2Cl_2$/MeOH = 9/1) indicated complete conversion of the starting material and the product was used for the next step without further purification.

Yield: quantitative

TLC: $CH_2Cl_2$/MeOH = 9/1; $R_f$ = 0.8

*(2,8-$^{13}$C)-N-6-methyladenosine (3)*

The brownish foam of **2** was treated with a 1:1 mixture of methylamine solution (20 ml, 33 wt% in absolute ethanol) and methylamine solution (20 ml, 40 wt% in water) and stirred at room temperature for 48 hours. After TLC analysis ($CH_2Cl_2$/MeOH = 9/1) showed complete conversion of the starting material all solvents were evaporated and the oily residue was coevaporated trice with methanol.

Adsorption of the residue onto silica from methanol and subsequent silica gel flash column chromatography using a gradient from 0 to 20% of methanol in ethyl acetate yielded compound **3** as white solid. The product was dried in high vacuum.

Yield: 670 mg (2.37 mmol, 69 % referred to **1**)

TLC: $CH_2Cl_2$/MeOH = 9/1; $R_f$ = 0.1

$^1$H-NMR (300 MHz, DMSO-$d_6$, 25°C): δ 8.33 (d, 1H, $^1J_{CH}$ = 213.3 Hz, $^{13}$C(8)*H*); 8.21 (d, 1H, $^1J_{CH}$ = 199.4 Hz, $^{13}$C(2)*H*); 7.82 (s, 1H, N*H*); 5.87 (dd, 1H, C(1')H); 5.43 (d, 1H, C(2')O*H*); 5.41 (d, 1H, C(5')O*H*); 5.18 (d, 1H, C(3')O*H*); 4.61 (dd, 1H, C(2')*H*); 4.14 (dd, 1H, C(3')*H*); 3.96 (m, 1H, C(4')*H*); 3.73-3.50 (m, 2H, C(5')*H*; C(5'')*H*); 2.96 (s, 3H, NHC*H*$_3$) ppm.

$^{13}$C-NMR (75 MHz, DMSO-$d_6$, 25°C): δ 152.4 ($^{13}$C(2)); 139.6 ($^{13}$C(8)); 87.9 (C(1')); 85.9 (C(4')); 73.5 (C(2')); 70.6 (C(3')); 61.6 (C(5')); 26.8 ($NHCH_3$) ppm.

*3',5'-O-(1,1,3,3-tetra-isopropyldisiloxane-1,3-diyl)-(2,8-$^{13}$C)-N-6-methyladenosine (4)*

Compound **3** (1.24 g, 4.38 mmol) was dissolved in anhydrous pyridine (15 ml) under argon atmosphere and cooled to 0°C using an ice bath. Then 1,1,3,3-tetraisopropyl-1,3-dichlorodisiloxane (1.1 eq, 1,52g, 4.82 mmol) was added dropwise and the solution was stirred for 3 hours at room temperature. After full conversion of the starting material was indicated by TLC analysis (EtOAc) the reaction was quenched with methanol and all solvents were evaporated. The residue was dissolved in dichloromethane, washed twice with 5% citric acid, twice with saturated sodium bicarbonate solution, dried over $Na_2SO_4$, filtered and evaporated to dryness. The crude product was applied to a silica gel column with methylene chloride and eluted using a gradient from 60 to 80% of ethyl acetate in hexane to give compound **4** as white foam. The product was dried in high vacuum.

Yield: 1.47g (2.80 mmol, 64 %)

TLC: EtOAc; $R_f$ = 0.5

$^1$H-NMR (300 MHz, DMSO-$d_6$, 25°C): δ 8.19 (d, 1H, $^1J_{CH}$ = 213.1 Hz, $^{13}$C(8)*H*); 8.16 (d, 1H, $^1J_{CH}$ = 205.3 Hz, $^{13}$C(2)*H*); 7.83 (s, 1H, N*H*); 5.88 (d, 1H, C(1')H); 5.62 (d, 1H, O*H*); 4.80 (dd, 1H, C(3')*H*); 4.51 (m, 1H, C(2')*H*); 4.00 (m, 1H, C(4')*H*); 4.1-3.87 (m, 2H, C(5')*H*, C(5'')*H*); 2.95 (s, 3H, N(6)C*H*$_3$); 1.07-0.99 (m, 28H, i-Pr C*H*$_3$, C*H*) ppm.

$^{13}$C-NMR (75 MHz, DMSO-$d_6$, 25°C): δ 152.9 ($^{13}$C(2)); 139.2 ($^{13}$C(8)); 89.6 (C(1')); 81.2 (C(4')); 74.1 (C(2')); 70.2 (C(3')); 61.1 (C(5')); 27.6 (N(6)$CH_3$); 1.0 (i-Pr) ppm.

*2'-O-(1-imidazolyl)thiocarbonyl-3',5'-O-(1,1,3,3-tetra-isopropyldisiloxane-1,3-diyl)- (2,8-$^{13}$C)-N-6-methyladenosine (5)*

Compound **4** (1.47 g, 2.80 mmol) was dissolved in anhydrous 1,2-dichloro ethane (18 ml) under an argon atmosphere. Then a spatula tip 4-(dimethylamino)pyridine and 1,1′-thiocarbonyldiimidazole (1.5 eq, 750 mg, 4.19 mmol) were added and the reaction was stirred at 85°C under reflux for 2 hours. After TLC analysis ($CH_2Cl_2$/MeOH = 95/5) indicated complete conversion of the starting material the solution was diluted with dichloromethane and washed once with saturated sodium bicarbonate solution. The organic layer was dried over anhydrous sodium sulfate, filtered and evaporated. Final purification using silica gel flash chromatography with a gradient from 0% to 4% methanol in dichloromethane yields compound **5** as pale yellow foam. The product was dried in high vacuum.

Yield: 1.62 g (2.55 mmol, 91 %)

TLC: $CH_2Cl_2$/MeOH = 95/5; $R_f$ = 0.4

$^1$H-NMR (300 MHz, $CDCl_3$, 25°C): δ 8.33 (d, 1H, $^1J_{CH}$ = 201.4 Hz, $^{13}$C(2)*H*); 8.38 (s, 1H, C(2)*H* imidazol); 7.84 (d, 1H, $^1J_{CH}$ = 211.6 Hz, $^{13}$C(8)*H*); 7.67 (t, 1H, C*H* imdidazol); 7.08 (m, 1H, C*H* imidazol); 6.41 (d, 1H, C(2')*H*); 6.12 (d, 1H, C(1')*H*); 5.84 (d, 1H, N*H*); 5.57 (dd, 1H, C(3')*H*); 4.11 (m, 1H, C(4')*H*); 4.22-4.03 (m, 2H, C(5')*H*, C(5'')*H*); 3.21 (d, 3H, NHC*H*3); 1.15-0.93 (m, 28H, iPr) ppm.

$^{13}$C-NMR (75 MHz, $CDCl_3$, 25°C): δ 153.7 ($^{13}$C(2)); 138.7 ($^{13}$C(8)); 136.8 (C() imidazol); 130.9 (C(4) imidazol); 117.8 (C(5) imidazol); 87.2 (C(1')); 83.8 (C(2')); 82.5 (C(4')); 69.8 (C(3')); 60.5 (C(5')); 27.4 ($NHCH_3$); 19.1-11.5 (iPr) ppm.

*3',5'-O-(1,1,3,3-tetra-isopropyldisiloxane-1,3-diyl)-(2,8-$^{13}$C)-N-6-methyl-2'-deoxyadenosine (6)*

Compound **5** (1.62 g, 2.55 mmol) was dissolved in anhydrous and degassed toluene (50 ml) under argon atmosphere. Then, tributyltin hydride (1.5 eq, 1.03 ml, 3.82 mmol) and azobisisobutyronitrile (0.2 eq, 84 mg, 0.51 mmol) was added and the reaction was stirred at 75°C. After 1 ½ hours TLC analysis (EtOAc) showed complete conversion of the starting material and the solvent was evaporated. The residue was dissolved in dichloromethane and washed twice with saturated sodium bicarbonate solution, dried over anhydrous sodium sulfate, filtered and evaporated. The crude product was applied to a silica gel column with methylene chloride and eluted using a gradient from 50 to 80% of ethyl acetate in hexane to give compound **4** as white foam. The product was dried in high vacuum.

Yield: 1.14 g (2.24 mmol, 88 %)

TLC: EtOAc; $R_f$ = 0.6

$^1$H-NMR (300 MHz, $CDCl_3$, 25°C): δ 8.39 (d, 1H, $^1J_{CH}$ = 200.5 Hz, $^{13}$C(2)*H*); 7.97 (d, 1H, $^1J_{CH}$ = 211.3 Hz, $^{13}$C(8)*H*); 6.27 (dd, 1H, C(1')*H*); 5.85 (d, 1H, N*H*); 4.96 (m, 1H, C(3')); 4.04 (m, 2H, C(5')*H*, C(5'')*H*); 3.89 (m, 1H, C(4')*H*); 3.19 (s, 3H, $NHCH_3$); 2.76-2.56 (m, 2H, C(2')*H*, C(2'')*H*); 1.13-1.00 (m, 28H, iPr) ppm.

$^{13}C$-NMR (75 MHz, $CDCl_3$, 25°C): δ 153.1 ($^{13}C(2)$); 138.0 ($^{13}C(8)$); 85.2 (C(4')); 83.0 (C(1')); 70.0 (C(3')); 61.9 (C(5')); 27.5 ($NHCH_3$); 19.2-11.4 (iPr) ppm.

*(2,8-$^{13}C$)- N-6-methyl-2'-deoxyadenosine (7)*

Compound **6** (1.14 g, 2.24 mmol) was dissolved in anhydrous tetrahydrofuran (20 ml) under argon atmosphere. Then triethylamine trihydrofluoride (1.3 eq, 474 µl, 2.91 mmol) was added and the reaction was stirred at 45°C for 2 hours. After TLC analysis ($CH_2Cl_2$/MeOH = 9/1) showed complete conversion of the starting material all solvents were removed. The crude residue was dissolved in methanol, adsorbed on silica, applied onto a silica gel column and the product was eluted using a gradient of 0 to 20% of methanol in ethyl acetate. The product was dried in high vacuum.

Yield: 590 mg (2.21 mmol, 98 %)

TLC: $CH_2Cl_2$/MeOH = 9/1; $R_f$ = 0.1

$^1H$-NMR (300 MHz, DMSO-$d_6$, 25°C): δ 8.32 (d, 1H, $^1J_{CH}$ = 213.1 Hz, $^{13}C(8)H$); 8.22 (d, 1H, $^1J_{CH}$ = 199.4 Hz, $^{13}C(2)H$); 7.77 (s, 1H, N*H*); 6.35 (m, 1H, C(1')*H*); 5.30 (d, 1H, C(3')O*H*); 5.24 (dd, 1H, C(5')O*H*); 4.41 (m, 1H, C(3')*H*); 3.88 (m, 1H, C(4')*H*); 3.61 (m, 1H, C(5')*H*); 3.53 (m, 1H, C(5'')*H*); 2.95 (s, 3H, $NHCH_3$); 2.72 (m, 1H, C(2')*H*); 2.54 (m, 1H, C(2'')*H*) ppm.

$^{13}C$-NMR (75 MHz, DMSO-$d_6$, 25°C): δ 151.9 ($^{13}C(2)$); 138.7 ($^{13}C(8)$); 87.9 (C(4')); 83.6 (C(1')); 70.8 (C(3')); 61.7 (C(5')); 39.2 (C(2')); 26.8 ($NHCH_3$) ppm.

*5'-O-(4,4'-dimethoxytrityl)-(2,8-$^{13}C$)- N-6-methyl-2'-deoxyadenosine (7)*

Compound **6** (590 mg, 2.21 mmol) was dissolved in anhydrous pyridine (20 ml) under argon atmosphere. Then 4,4'-dimethoxytrityl chloride (1.2 eq, 898 mg, 2.65 mmol) was added in three portions within 30 minutes. The reaction was stirred at room temperature overnight. After full conversion was indicated by TLC analysis ($CH_2Cl_2$/MeOH = 95/5) the reaction was quenched with methanol and all solvents were evaporated. The oily residue was coevaporated trice with dichloromethane, dissolved in dichloroethane and washed twice with 5% citric acid and twice with saturated sodium bicarbonate solution. The combined organic layers were dried over anhydrous sodium sulfate, filtered and evaporated. The crude product was applied to a silica gel column with methylene chloride and eluted using a gradient from 60 to 100% of ethyl acetate in hexane to give compound **7** as white foam.

Yield: 920 mg (1.62 mmol, 73 %)

TLC: $CH_2Cl_2$/MeOH = 95/5; $R_f$ = 0.5

$^1H$-NMR (300 MHz, DMSO-$d_6$, 25°C): δ 8.23 (d, 1H, $^1J_{CH}$ = 212.2 Hz, $^{13}C(8)H$); 8.17 (d, 1H, $^1J_{CH}$ = 199.2 Hz, $^{13}C(2)H$); 7.73 (s, 1H, N*H*); 7.36-6.73 (m, 13H, C*H* arom); 6.36 (m, 1H, C(1')*H*); 5.35 (d, 1H,

C(3')O*H*); 4.48 (m, 1H, C(3')*H*); 3.98 (m, 1H, C(4')); 3.72 (s, 3H, OC$H_3$); 3.71 (s, 3H, OC$H_3$); 3.16 (d, 2H, C(5')*H*, C(5'')*H*); 2.95 (s, 3H, NHC*H*3); 2.88 (m, 1H, C(2')*H*); 2.32 (m, 1H, C(2'')*H*) ppm.
<u>$^{13}$C-NMR (75 MHz, DMSO-d$_6$, 25°C):</u> δ 152.3 ($^{13}$C(2)); 138.8 ($^{13}$C(8)); 131.4-111.0 (C(arom)); 85.7 (C(4')); 83.1 (C(1')); 70.5 (C(3')); 64.0 (C(5')); 56.2-53.1 ($OCH_3$); 38.4 (C(2')); 26.6 ($NHCH_3$) ppm.

*<u>5'-O-(4,4'-dimethoxytrityl)-(2,8-$^{13}$C)-N-6-methyl-2'-deoxyadenosine3'-[(2-cyanoethyl)-(N,N-diisopropyl)]phosphoramidite(8)</u>*

Compound **6** (920 mg, 1.62 mmol) was dried under high vacuum overnight and then dissolved in anhydrous tetrahydrofuran (20 ml) under argon atmosphere. Then N,N-diisopropylethylamine (5 eq, 1.41 ml, 8.08 mmol) and subsequently 2-cyanoethyl-N,N-diisopropylchlorophosphoramidite (1.3 eq, 497 mg, 2.10 mmol) were added to the solution and the mixture was stirred for 2 hours at room temperature. After full conversion was indicated by TLC analysis ($CH_2Cl_2$/MeOH = 95/5) the reaction was quenched with methanol. The mixture was diluted with dichloromethane and washed twice with half saturated sodium bicarbonate solution. The organic layer was dried over anhydrous sodium sulfate, filtered and evaporated to dryness. The crude product was applied to a silica gel column with dichloromethane and eluted using a gradient from 6:4 to 1:0 ethyl acetate/n-hexane (+ 1 % triethylamine) to give **8** as a colorless foam. The product was dried in high vacuum.
<u>Yield:</u> 980 mg (1.27mg, 79 %)
<u>TLC:</u> $CH_2Cl_2$/MeOH = 95/5; $R_f$ = 0.6
<u>$^1$H-NMR (300 MHz, DMSO-d$_6$, 25°C):</u> δ 8.24 (d, 1H, $^1J_{CH}$ = 212.5 Hz, $^{13}$C(8)*H*); 8.13 (d, 1H, $^1J_{CH}$ = 199.3 Hz, $^{13}$C(2)*H*); 7.74 (s, 1H, N*H*); 7.37-6.73 (m, 13H, C*H* arom); 6.37 (m, 1H, C(1')*H*); 4.79 (m, 1H, C(3')*H*); 4.10 (m, 1H, C(4')*H*); 3.71 (s, 6H, OC$H_3$); 3.66 (m, 2H, POC$H_2$); 3.55 (m, 2H, C*H* iPr); 3.22 (m, 2H, C(5')*H*, C(5'')*H*); 3.07 (m, 1H, C(2')*H*); 2.94 (s, 3H, NHC$H_3$); 2.77 (t, 1H, C$H_2$CN); 2.67 (t, 1H, C$H_2$CN); 2.47 (m, 1H, C(2'')*H*); 1.17-1.01 (m, 12H, C$H_3$ iPr) ppm.
<u>$^{13}$C-NMR (75 MHz, DMSO-d$_6$, 25°C):</u> δ 152.3 ($^{13}$C(2)); 139.1 ($^{13}$C(8)); 131.2-111.0 (C(arom)); 84.7 (C(4')); 83.2 (C(1')); 72.8 (C(3')); 63.2 (C(5')); 58.2 ($POCH_2$); 56.3-52.6 ($OCH_3$); 42.4 (CH iPr); 37.1 (C(2')); 26.9 ($NCH_3$); 25.9-22-4 ($CH_3$ iPr); 19.6 $CH_2$CN) ppm.
<u>$^{31}$P-NMR (121 MHz, CDCl3, 25°C):</u> δ 148.9, 148.3 ppm.
<u>ESI-MS:</u> 770.37 $[M+H]^+$; 792.35 $[M+Na]^+$; 808.32 $[M+K]^+$ m/z.

### Supplementary Figures

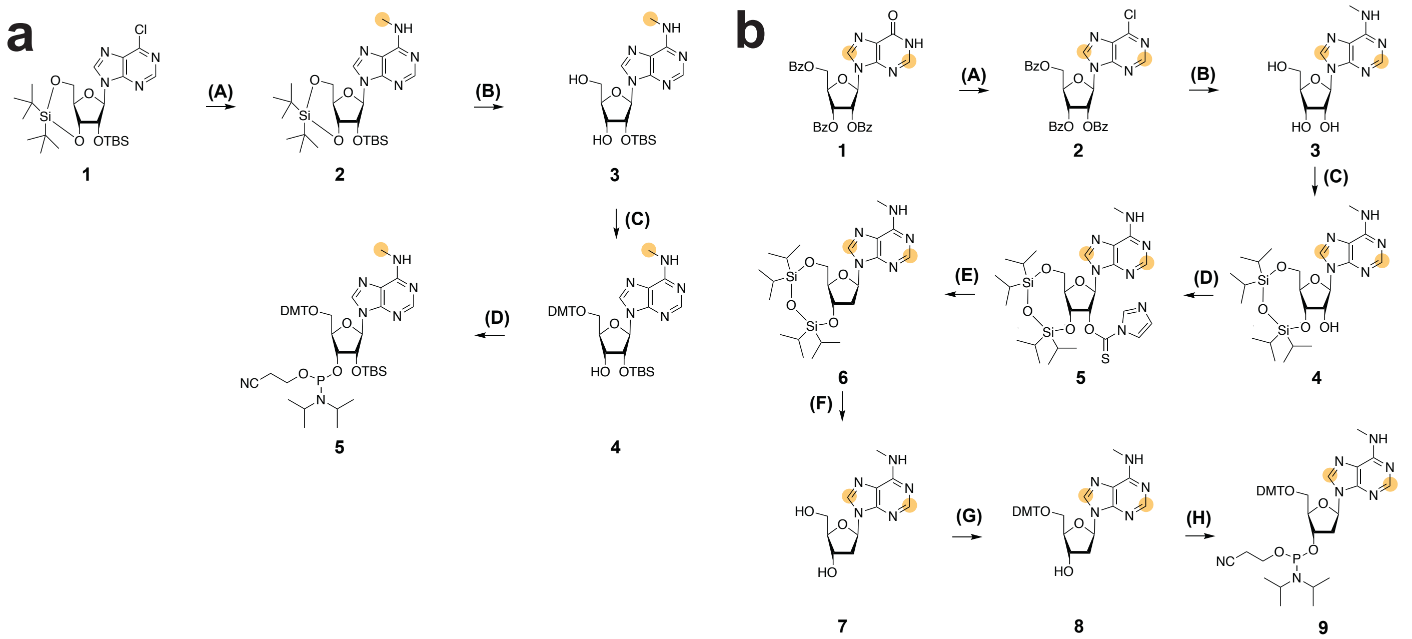


**Supplementary Fig. 1.** Synthesis of $^{13}$C labeled phosphoramidites. **a**, Synthesis of ($^{13}CH_3$)-m$^6$A RNA phosphoramidite. **(A)** $^{13}$C-MeNH$_2$-HCl, Et$_3$N in DMF, 3 d, RT; **(B)** HF-Py in pyridine, 3 h, 0°C; **(C)** DMT-Cl, DMAP in pyridine, 3 h, RT; **(D)** CEP-Cl, DIPEA in THF, 2 h, RT. **b**, Synthesis of $^{13}$C8,$^{13}$C2-labeled m$^6$dA phosphoramidite. **(A)** N,N-Dimethylaniline in POCl3, 120°C 13 min; **(B)** 1/1 methylamine (33 wt% in absolute ethanol) and methylamine (40 wt% in water), 48 h, RT; **(C)** TIPDSiCl$_2$ in pyridine, 3 h, RT; **(D)** TCDI, DMAP in DCE, 85°C 2h **(E)** Bu$_3$SnH, AIBN in toluene, 75°C 1 ½ h; **(F)** TEA-HF in THF, 45°C 2h; **(G)** DMT-Cl in pyridine, overnight RT; **(h)** CEP-Cl, DIPEA in THF, 2 h, RT.

### References


1. Engel, J.D. & von Hippel, P.H. Effects of methylation on the stability of nucleic acid conformations: studies at the monomer level. *Biochemistry* **13**, 4143-58 (1974).
2. Shi, H. et al. NMR Chemical Exchange Measurements Reveal That N(6)-Methyladenosine Slows RNA Annealing. *J Am Chem Soc* **141**, 19988-19993 (2019).
3. Berman, H.M. et al. The Protein Data Bank. *Nucleic Acids Res* **28**, 235-42 (2000).
4. Abou Assi, H. et al. 2'-O-Methylation can increase the abundance and lifetime of alternative RNA conformational states. *Nucleic Acids Res* (2020).
5. Dethoff, E.A., Petzold, K., Chugh, J., Casiano-Negroni, A. & Al-Hashimi, H.M. Visualizing transient low-populated structures of RNA. *Nature* **491**, 724-8 (2012).
6. Chu, C.C., Plangger, R., Kreutz, C. & Al-Hashimi, H.M. Dynamic ensemble of HIV-1 RRE stem IIB reveals non-native conformations that disrupt the Rev-binding site. *Nucleic Acids Res* **47**, 7105-7117 (2019).
7. Cisse, II, Kim, H. & Ha, T. A rule of seven in Watson-Crick base-pairing of mismatched sequences. *Nat Struct Mol Biol* **19**, 623-7 (2012).
8. Bang, J., Bae, S.H., Park, C.J., Lee, J.H. & Choi, B.S. Structural and dynamics study of DNA dodecamer duplexes that contain un-, hemi-, or fully methylated GATC sites. *J Am Chem Soc* **130**, 17688-96 (2008).
9. Gueron, M. & Leroy, J.L. Studies of base pair kinetics by NMR measurement of proton exchange. *Methods Enzymol* **261**, 383-413 (1995).
10. Chu, C.C., Liu, B., Plangger, R., Kreutz, C. & Al-Hashimi, H.M. m6A minimally impacts the structure, dynamics, and Rev ARM binding properties of HIV-1 RRE stem IIB. *PLoS One* **14**, e0224850 (2019).
